# Optogenetic control reveals differential promoter interpretation of transcription factor nuclear translocation dynamics

**DOI:** 10.1101/548255

**Authors:** Susan Y. Chen, Lindsey C. Osimiri, Michael Chevalier, Lukasz J. Bugaj, Andrew H. Ng, Jacob Stewart-Ornstein, Lauren T. Neves, Hana El-Samad

## Abstract

The dynamic translocation of transcription factors (TFs) in and out of the nucleus is thought to encode information, such as the identity of a stimulus. A corollary is the idea that gene promoters can decode different dynamic TF translocation patterns. Testing this TF encoding/promoter decoding hypothesis requires tools that allow direct control of TF dynamics without the pleiotropic effects associated with general perturbations. In this work, we present CLASP (Controllable Light Activated Shuttling and Plasma membrane sequestration), a tool that enables precise, modular, and reversible control of TF localization using a combination of two optimized LOV2 optogenetic constructs. The first sequesters the cargo in the dark at the plasma membrane and releases it upon exposure to blue light, while light exposure of the second reveals a nuclear localization sequence that shuttles the released cargo to the nucleus. CLASP achieves minute-level resolution, reversible translocation of many TF cargos, large dynamic range, and tunable target gene expression. Using CLASP, we investigate the relationship between Crz1, a naturally pulsatile TF, and its cognate promoters. We establish that some Crz1 target genes respond more efficiently to pulsatile TF inputs than to continuous inputs, while others exhibit the opposite behavior. We show using computational modeling that efficient gene expression in response to short pulsing requires fast promoter activation and slow inactivation and that the opposite phenotype can ensue from a multi-stage promoter activation, where a transition in the first stage is thresholded. These data directly demonstrate differential interpretation of TF pulsing dynamics by different genes, and provide plausible models that can achieve these phenotypes.

## Introduction

Transcription factors (TFs) are key mediators in the transmission of information from the internal and external environment of the cell to its genome. Understanding how TFs encode information about the environment in order to coordinate transcriptional programs remains one of the most pressing problems in molecular and systems biology. Many studies have explored how modulation of TF concentration, TF post-translational modification, and combinatorial TF control can yield differential gene regulation (Czyz et al., 1993; Sadeh et al., 2011; Springer et al., 2003), therefore explaining many important aspects of TF function and their information encoding capacity. These mechanisms, however, may not fully account for the complexity of signal multiplexing that is carried out by TFs. As a result, it has been proposed that TFs might also encode information in their spatio-temporal dynamics, using for example different patterns of nuclear shuttling for different inputs.

A number of studies have attempted to elucidate this TF dynamic encoding hypothesis by eliciting different TF dynamic patterns using various environmental inputs and assessing the consequences (Batchelor et al., 2011; Covert et al., 2005; Gotoh et al., 1990; Hoffmann et al., 2002; Nelson et al., 2004; Nguyen et al., 1993; Purvis and Lahav, 2013; Purvis et al., 2012; Tay et al., 2010; Traverse et al., 1992; Werner et al., 2008). For example, it was shown that p53 exhibits fixed concentration pulses in response to gamma radiation, but implements only one amplitude- and duration-dependent continuous pulse in response to UV (Batchelor et al., 2011). These two pulsing regimes have different physiological outcomes, with the former leading to cell cycle arrest and the latter leading to cell death (Purvis et al., 2012). Other studies programmed different TF nuclear translocation patterns by gaining control of a signaling node upstream of the TF. A prominent example of this approach is the modulation of Msn2 dynamics by regulating protein kinase A (PKA). Inhibition of an analog-sensitive PKA by a small-molecule resulted in Msn2 translocation to the nucleus (Hansen and O’Shea, 2013, 2015a, 2015b; Hao and O’Shea, 2011; Hao et al., 2013). With this method, it was shown that genes in the Msn2 regulon can be differentially modulated by the amplitude, duration, and frequency of Msn2 nuclear translocation pulses.

In the budding yeast *Saccharomyces cerevisiae*, there are approximately 200 known TFs, two-thirds of which are constitutively localized to the nucleus; the remaining one-third are located in the cytoplasm during exponential growth in complete media (Chong et al., 2015). At least nine of these basally cytoplasmic TFs transiently localize into the nucleus in response to various stress conditions (Dalal et al., 2014). Furthermore, different environmental conditions elicit a range of pulsing characteristics for these TFs that differ in their duration, amplitude, and frequency (Dalal et al., 2014) (Figure S1), suggesting that reversible TF nuclear localization may encode regulatory information. This information is then decoded by downstream target genes in order to produce an appropriate response (Granados et al., 2018).

Control of TF localization through modulation of upstream regulators with small molecules or chemicals has been an essential method to put forward such a hypothesis of TF dynamic encoding (AkhavanAghdam et al., 2016; Cai et al., 2008; Hansen and O’Shea, 2013, 2015a, 2015b; Hao and O’Shea, 2011; Lin and Doering, 2016; Purvis et al., 2012). However, this method produces pleiotropic effects that can be hard to untangle. For example, PKA controls many transcriptional regulators in addition to Msn2. As a result, modulating its activity with a small molecule may yield gene expression changes that are not solely caused by Msn2 translocation dynamics but are instead the result of combinatorial gene regulation by other PKA-responsive TFs such as Msn4 (AkhavanAghdam et al., 2016; Garmendia-Torres et al., 2007) and Dot6 (Pincus et al., 2014).

Therefore, to causally and quantitatively probe the relationship between TF nuclear localization dynamics and transcriptional activity, a method by which TFs can be specifically, quickly, and reversibly localized to the nucleus is needed. Specificity is necessary to allow direct regulation of TF nuclear localization without pleiotropic effects, while speed and reversibility are necessary to recapitulate the minutes-level resolution with which TFs translocate into and out of the nucleus in response to environmental inputs. Ideally, this method would also work modularly with many TF cargos, including TFs that are basally nuclear.

Here, we present CLASP, an optimized optogenetic tool that can exert precise, modular, and reversible control of transcription factor localization. CLASP uses two LOV2 light-responsive domains derived from *Avena sativa* to sequester a cargo at the plasma membrane in the dark and target it to the nucleus in response to blue light. We demonstrate how CLASP can be used as a general strategy to control many TF cargos without any further optimization. We exploit the characteristics of CLASP to control the localization of Crz1, a pulsatile TF, and show unambiguously that its target promoters have different abilities to differentially interpret pulsatile dynamics. Using data-backed computational modeling, we explore the principles by which they can do so. Our studies reveal that a more efficient response to short pulsed inputs can be achieved by a simple two-state promoter model with fast activation and slow shut-off. By contrast, to achieve more efficient gene expression from continuous inputs than from pulsed inputs, a more complicated model with at least two activation steps or thresholding needs to be invoked. Furthermore, to achieve this property in combination with a graded dose response, a promoter model needs to minimally combine two activation steps and thresholding, with a dependence of both activation steps on the TF. These results, made possible through the productive use of CLASP in iteration with computational modeling, pave the way for more thorough understanding of the general principles by which gene promoters can interpret TF dynamics.

## Results

### Construction and optimization of CLASP, a dual-LOV2 optogenetic strategy for control of nuclear shuttling

Some optogenetic tools that rapidly translocate protein cargos to the nucleus have been developed (Niopek et al., 2014; Redchuk et al., 2017; Yumerefendi et al., 2015). A number of these tools utilized LOV2, a light-responsive protein often isolated from *A.sativa*, to uncage a nuclear localization sequence (NLS) in response to blue light. Uncaging of this NLS caused the translocation of the optogenetic molecule to the nucleus along with any appended protein cargo. Light Activated Nuclear Shuttle (LANS) is an example of this strategy (Yumerefendi et al., 2015) (Figure 1A). While effective for some applications, the architecture of this class of optogenetic tools may cause leaky nuclear localization for some basally cytoplasmic TFs. An example of this phenotype is Msn2, which, in many cells, exhibited constitutive nuclear localization when fused to LANS even in the absence of light stimulation (Figure S2). Moreover, tools such as LANS cannot be used to regulate localization of basally nuclear TFs, as there is no mechanism for preventing the endogenous nuclear localization of these TF cargos.

**Figure 1.**
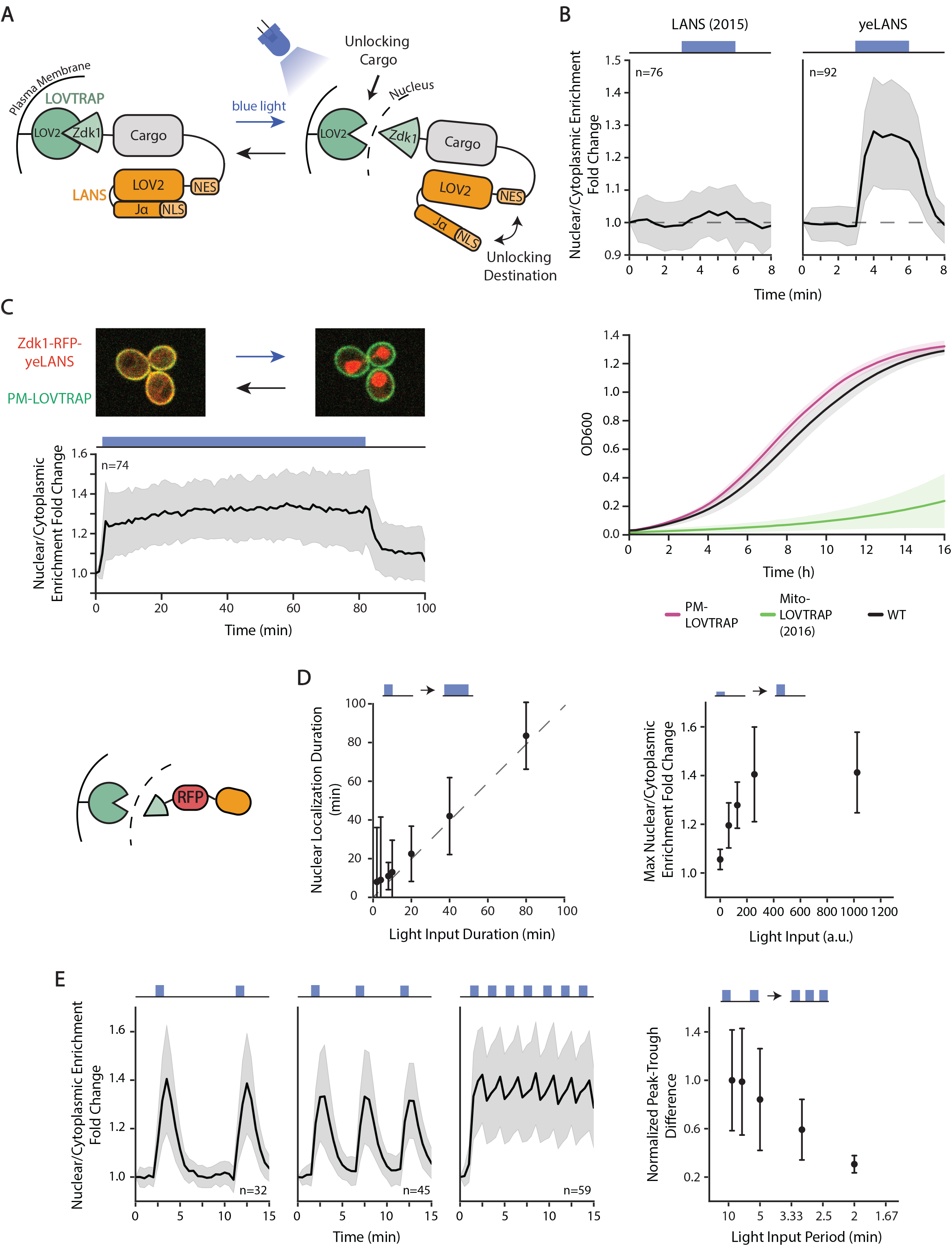
Design, Optimization, and Characterization of CLASP. **A)** Schematic illustrating CLASP mechanism. **B)** Optimization of LANS NLS **(top panels)** and LOVTRAP localization **(bottom panel)**. Top panels show mean value of nuclear/cytoplasmic enrichment fold change for original NLS and optimized NLS (yeLANS) as a function of time when given a pulse of blue light. Nuclear/Cytoplasmic enrichment fold change is calculated relative to the nuclear/cytoplasmic enrichment at t=0. Bottom panel shows mean of OD600 in 3 growth experiments for original LOVTRAP targeted to mitochondria in addition to the optimized plasma membrane targeted LOVTRAP. **C) (top panel)** Confocal microscopy image showing mScarlet-CLASP localization at the plasma membrane in the dark (left) and in the nucleus (right) after 3 minutes of light exposure. **(bottom panel)** Quantification of mean nuclear/cytoplasmic enrichment fold change of mScarlet-CLASP as a function of time in response to a prolonged light input (80 minutes, 1024 a.u. light input amplitude). Black line represents the mean of 74 cells. **D)** Quantification of the response of mScarlet-CLASP to light inputs with different dynamic characteristics. Left plot shows median time to return to within 25% of basal nuclear/cytoplasmic enrichment for light pulses of different durations and constant 1024 a.u. amplitude. Median is used to minimize the effect of outliers. The dotted line is Y=X line. Right plot shows the mean response to one minute light pulses of different amplitudes. **E)** Nuclear/cytoplasmic enrichment fold change of mScarlet-CLASP in response to light pulsing with different periods. Left three graphs show mean enrichment fold change as a function of time in response to pulsed light inputs (1 minute light given in a 9, 5, or 2 minute period, respectively) with 1024 a.u. amplitude. Right plot quantifies median peak-to-trough difference (normalized to the median peak-to-trough difference generated by the shortest period). Median is used to minimize the effect of outliers. Error bars and shaded area, except where noted, represent standard deviation to show the spread of the data. For all panels, n represents the number of cells tracked and light input regimes are depicted on top of panels. Cartoon (**left of D**) represents mScarlet-CLASP. yeLANS -- yeast enhanced LANS, PM-LOVTRAP -- Plasma Membrane LOVTRAP, Mito-LOVTRAP -- Mitochondrial LOVTRAP.

A different optogenetic tool, LOVTRAP, a LOV2-based tool for protein sequestration, could be used for rapid translocation of cargo with less leaky basal localization. LOVTRAP is composed of a LOV2 fused to the mitochondria and Zdk1, a small peptide that is fused to the protein cargo. The interaction of LOV2 and Zdk1 in the dark sequesters the cargo to the surface of the mitochondria (Wang et al., 2016) (Figure 1A). However, LOVTRAP alone does not contain targeting information, and hence cannot direct the cargo to the nucleus on demand. Therefore, to enable both robust and targeted optogenetic control of many different cargos, we sought to use LOVTRAP in concert with LANS. The idea of combining optogenetic sequestration and nuclear localization was previously investigated (Redchuk et al., 2017; Yumerefendi et al., 2018). However, the resulting tools either required complex dual color stimulation (Redchuk et al., 2017), thereby limiting the number of fluorescent proteins that could be used in a cell, or did not demonstrate modularity for different cargos (Yumerefendi et al., 2018). These tools also lacked optimization for use in yeast.

To construct a modular and specific tool for yeast protein nuclear translocation, we first tackled optimization of the published LANS and LOVTRAP constructs. Fluorescently-tagged (mCherry) LANS (Yumerefendi et al., 2015) displayed only a moderate increase (3.4%) in nuclear/cytoplasmic enrichment in response to blue light (Figure 1B, upper left panel). This increase was much weaker than that seen for transcription factors in response to stress inputs (Figure S1, 20-50% increase). Additionally, the published LOVTRAP tool used a TOM20 mitochondrial targeting tag that caused a strong growth defect in yeast at high expression levels (Figure 1B, lower panel). LOVTRAP sequestration had previously been shown to perform best when the mitochondria-bound LOV2 trap was expressed in excess of the Zdk1; as a result, these high expression levels were necessary for trapping many protein cargos, causing the growth defect to be an issue.

To improve LANS localization properties, we replaced the published LANS NLS with a small library of yeast NLS peptides (Kosugi et al., 2009). We then screened blue light induced nuclear localization of mCherry-LANS constructs that had any one of these different NLS sequences. We identified a number of NLS sequences that showed an improvement in nuclear/cytoplasmic enrichment in response to blue light (Figure S3A), including an NLS that increased the fold change by eight-fold. We chose this NLS sequence to move forward as a yeast enhanced LANS (yeLANS) (Figure 1B). Next, to rectify the growth defect associated with LOVTRAP sequestration to the mitochondria, we swapped the mitochondrial TOM20 tag with a plasma membrane Hs-RGS2 tag (Heximer et al., 2001) to create pm-LOVTRAP. This modification rescued the growth defect of LOVTRAP even at high expression levels (Figure S3B).

Finally, we combined yeLANS and pm-LOVTRAP to form CLASP (Controllable Light Activated Shuttling and Plasma membrane sequestration), a construct comprised of two AsLOV2 domains. The first AsLOV2 domain is fused to the plasma membrane and sequesters a Zdk1 fused to the N-terminus of the cargo (for example, a TF). The second AsLOV2 domain is fused to the C-terminus of the cargo. This AsLOV2 domain is preceded by a nuclear export sequence (NES) and has a nuclear localization sequence (NLS) embedded in the Jα helix. Blue light causes a conformational change in both AsLOV2 domains, yielding the simultaneous unlocking of cargo and its targeting to the nucleus (Figure 1A). Strains harboring CLASP did not experience any growth defect (Figure S3C).

We first tested CLASP with a red fluorescent protein (mScarlet) as a cargo. Confocal microscopy showed that mScarlet-CLASP was successfully sequestered at the membrane in the dark and translocates to the nucleus in response to blue light. Furthermore, widefield microscopy showed that nuclear localization could be maintained stably for at least 80 minutes (Figure 1C). Varying the duration of the light input demonstrated that CLASP could also track shorter light inputs (Figure S4). On average, mScarlet-CLASP nuclear localization extended four minutes longer than the duration of the input light pulse, illustrating its rapid shut-off time (Figure 1D, Supp Fig 4). The maximum nuclear/cytoplasmic enrichment achieved by mScarlet-CLASP was also graded as a function of light amplitude; when subjected to one minute pulses of increasing amplitude (64-1024 a.u.), enrichment increased commensurately for a wide range and saturated after 256 a.u. of light (Figure 1D, Table S4).

Finally, to test the ability of CLASP to respond to repeated light pulses and probe its dependence on their period, we subjected the cells to one minute pulses of blue light repeated every 2-9 minutes (Figure 1E, left 3 panels show one minute pulses every 2, 5, or 9 minutes). These experiments revealed that mScarlet-CLASP followed these pulses faithfully until the pulses became too rapid, that is, when the next light pulse occurred during the time required for nuclear exit (4 minutes). This effect occurred when pulses were repeated every 2 minutes, at which point nuclear localization became almost continuous at a high level. Quantification of the mean peak-to-trough difference in nuclear localization of single cell traces for different periodic light inputs showed a clear dependence on the period of the light pulse (Figure 1E).

Overall, our data indicate that mScarlet-CLASP could be rapidly, reversibly, and repeatedly localized to the nucleus as frequently as every five minutes and that the duration and the magnitude of this translocation could be robustly controlled.

### CLASP achieves precise, modular control of TF nuclear translocation and activation of target genes

The usefulness of CLASP depends on its ability to successfully control translocation of TF cargos while maintaining their function. Our next step was therefore to test the ability of CLASP to quickly and reversibly control the translocation of three basally cytoplasmic transcription factors to the nucleus. We chose a synthetic transcription factor, SynTF, constructed from Cys_2_-His_2_ zinc finger domains and a VP16 activation domain (Khalil et al., 2012), as well as Msn2, the principal transcription factor in the environmental stress response (Gasch et al., 2000), and Pho4, the principal transcription factor in the phosphate starvation response (Vardi et al., 2014). Both Msn2 and Pho4 have been known to translocate to the nucleus in response to stress (Dalal et al., 2014; Vardi et al., 2014). The three TF cargos were also tagged with a C-terminal RFP (mScarlet) for visualization.

For all three TFs, TF-CLASP achieved its maximal nuclear localization in response to light within one minute of blue light exposure. Like the mScarlet cargo, the TF cargos reversibly translocated to the nucleus as frequently as every five minutes when induced with a one minute pulse of light. Furthermore, a sustained light input produced continuous nuclear localization of the TFs, indicating that CLASP was capable of maintaining robust nuclear localization of associated TF cargos for an extended period of time (Figure 2A). The maximum nuclear/cytoplasmic enrichment fold change achieved with CLASP for Msn2 as a cargo was similar to that of Msn2 with a strong osmotic shock using 0.95M Sorbitol (Figure S1, Hoffmann et al., 2002).

**Figure 2.**
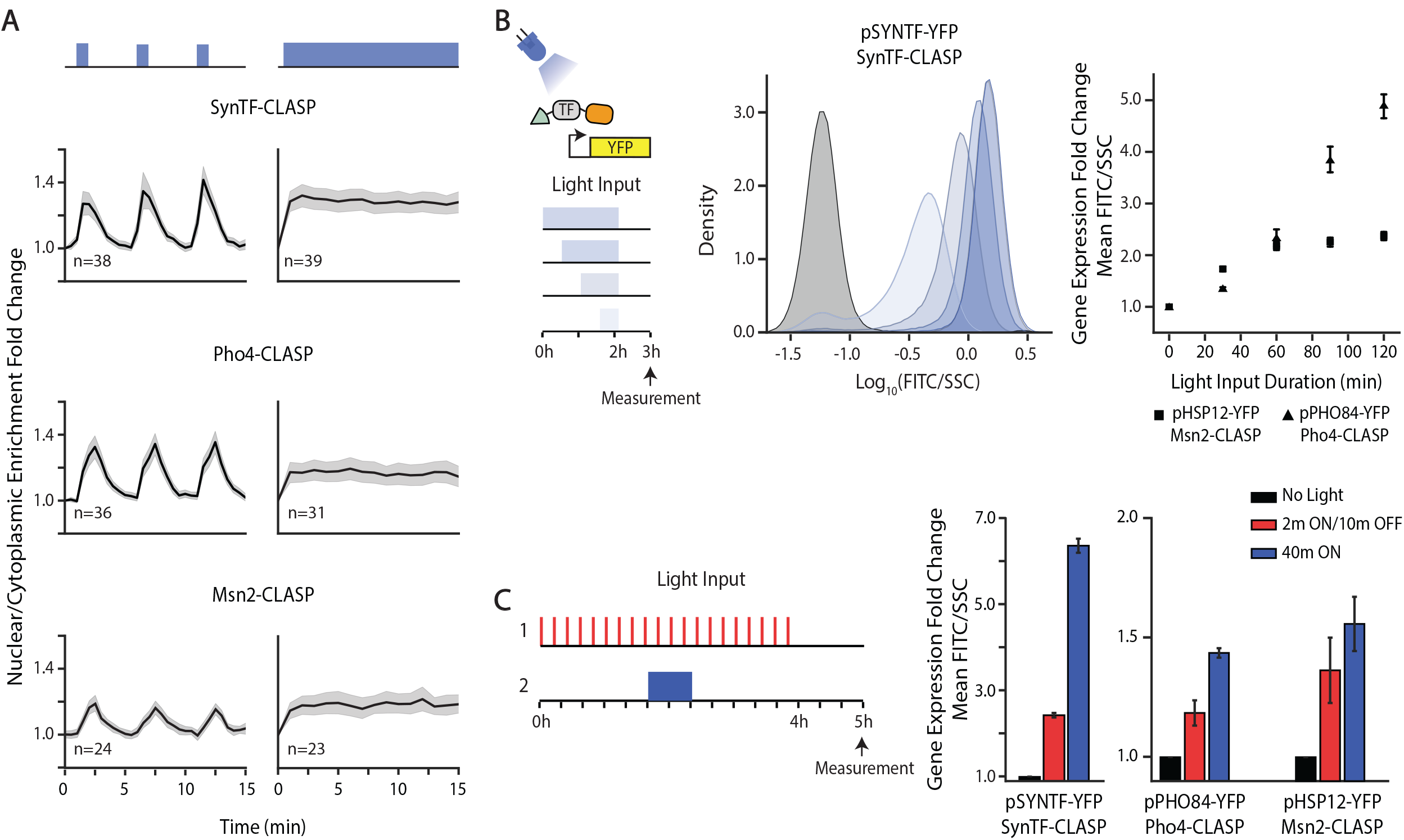
CLASP can be used to control localization of many transcription factor cargos. **A)** Nuclear/cytoplasmic enrichment fold change in response to pulsed **(left panels)** and continuous light **(right panels)** for several TF-CLASP cargos. Graph shows mean of single-cell traces for transcription factors tagged with CLASP. Light is delivered for one minute at the start of each five-minute period or continuously. Shaded gray area represents 95% confidence interval and light inputs are represented in blue above graphs. **B)** Fluorescent reporter expression due to TF-CLASP localization. Left panel shows a schematic of the experiment – the TF is localized to the nucleus for 0.5, 1, 1.5 or 2hrs. A fluorescent reporter is measured via flow cytometry one hour after light shut-off. Center panel shows the population response of pSYNTF-YFP (promoter downstream of SynTF-CLASP) for inputs shown on the left. Darker blue shades correspond to longer light duration. Black histogram corresponds to no light. Right panel shows quantification of the YFP fold change as a function of light duration for promoters responsive to other TF-CLASP constructs following the same experimental protocol. Fluorescence readings are normalized by side scatter and then normalized to the 0m dose for each strain to show fold change. Error bars represent standard error of the mean for 9 biologically independent replicates. **C)** Fluorescent reporter response to pulsatile versus continuous localization of different TF-CLASP constructs. TF-CLASP constructs are given either 20 two-minute pulses of light or 1 forty-minute pulse of light, as depicted in the schematic on the left. Reporter expression is measured via flow cytometry one hour after light shut-off. Right panels show quantification of YFP fold change in response to pulsed light input, continuous light input, or no input. Error bars represent standard error of the mean for 9 biologically independent replicates. In all panels, strains are induced with a given amplitude of light (SynTF-CLASP -- 1024 a.u.; Msn2-CLASP -- 2048 a.u., Pho4-CLASP -- 4095 a.u.).

To test whether nuclear localization of the TFs led to concomitant gene expression, we constructed promoter fusions expressing YFP with promoters that were responsive to SynTF (pSYNTF-YFP), Msn2 (pHSP12-YFP), and Pho4 (pPHO84-YFP). We then exposed these strains to fixed-amplitude light inputs (Figure S5A) of increasing duration (0.5-2 hours) and measured YFP fluorescence via flow cytometry. For all three TFs, increasing the duration of the light input led to increased downstream reporter gene expression, illustrating that the TF was still functional despite its fusion to CLASP. Notably, SynTF-CLASP yielded more than 20-fold activation of pSYNTF-YFP with only 2 hours of light activation (Figure 2B). Gene expression in the dark downstream of the three TF-CLASP constructs was similar to basal expression, and was also commensurate after light induction to gene expression generated by a constitutively nuclear TF (Figure S5B-D, Supplementary Text).

Next, we explored whether CLASP could control localization of transcription factors such as Gal4, which was basally nuclear. Gal4-CLASP was successfully sequestered to the plasma membrane in the dark and reversibly translocated to the nucleus in response to light. Nuclear translocation of Gal4-CLASP also activated expression from pGAL1, a Gal4-responsive promoter (Figure S6), indicating that CLASP was able to control TFs irrespective of their endogenous nuclear localization.

Finally, we sought to demonstrate that different TF dynamic translocation patterns generated with CLASP could yield different gene expression outputs. Several transcription factors, such as Pho4 following phosphate starvation, translocate into the nucleus in response to a stress input and reside there continuously until the response was completed (Vardi et al., 2014). Others, including Msn2 following a 0.4% glucose input, have been known to translocate into the nucleus with episodic and repeated pulses in response to an activating input (Dalal et al., 2014). Moreover, Msn2 has also been known to translocate with sustained pulses in response to osmotic shock (Figure S1B). As a result, we sought to explore the gene expression consequences of pulsing relative to continuous localization of the three CLASP-fused TFs (SynTF, Msn2 and Pho4). We delivered two light inputs that had different dynamic patterns but the same cumulative light duration of 40 minutes. In the first case, light was switched ON for 40 minutes, and in the second, light was given in 20 episodic pulses (2 minutes ON/10 minutes OFF) (Figure 2C). Delivery of the same cumulative light input and measurement at the end of the time course were necessary controls to compare efficiency of response of pulsed input relative to continuous inputs. YFP fluorescence was measured for both inputs after 5 hours using flow cytometry. These data showed that continuous nuclear residence of SynTF-CLASP, Msn2-CLASP, and Pho4-CLASP produced more gene expression than pulsed translocation, indicating that these promoters respond more efficiently to continuous inputs than pulsed inputs. This directly demonstrates that TF nuclear translocation dynamics could affect downstream reporter gene expression, an idea that we wanted to explore in more depth.

### CLASP control of the Crz1 TF reveals that its target genes differ in their efficiency of response to short pulses

To further explore the modes of decoding of TF dynamics by promoters in a biologically meaningful setting, we chose to focus on Crz1, the main TF in the calcineurin-Crz1 signaling pathway that responds to calcium stress. Crz1 has been shown to exhibit two modes of pulsatile nuclear translocation in response to calcium chloride (CaCl_2_) stress -- a single long initial pulse (40-60 min) and subsequent episodic repeated pulsing (1-4 min) (Figure S7A). We reasoned that continuous nuclear localization and pulsing of Crz1 could be interpreted differently by different target genes, a behavior that could be revealed and studied by controlling its localization using CLASP.

Crz1 has been shown to undergo phosphorylation on multiple residues to activate gene expression in calcium stress (Stathopoulos-Gerontides et al., 1999). Therefore, to survey the response of Crz1 target genes to dynamic inputs using CLASP, we needed to adopt a variant of Crz1 that bypassed this regulation, an endeavor that could be necessary for studying the effects of many TFs with CLASP. We therefore built a strain in which Crz1*, an alanine mutant with 19 S/T to A substitutions of Crz1, was used as a CLASP cargo. Crz1* was basally nuclear and bypassed the post-translational modification required of Crz1 (Figure S7B-C, Stathopoulos-Gerontides et al., 1999). To verify that Crz1* preserved the transcriptional profile of wild type Crz1, we carried out mRNA sequencing of cell populations in which the wild type allele of Crz1 was knocked out and Crz1* was expressed from a constitutive pADH1 promoter. We compared the up-regulated genes of the Crz1* strain (where Crz1* is basally nuclear) with genes upregulated by Crz1-yeLANS under CaCl_2_ stress. We found a good overlap between the two gene sets, with appropriate enrichment of GO terms that were indicative of a response to calcium stress (Figure S7C). By probing individual Crz1 target genes with fluorescent reporters, we also found that light-induced Crz1*-CLASP, but not light-induced Crz1-CLASP (Figure S7D), was able to elicit appreciable gene expression. For example, Crz1*-CLASP driving pPUN1-YFP, a canonical Crz1 responsive promoter, achieved similar gene expression fold change (1.8) as pPUN1-YFP in calcium stress (1.7) (Figure S7F). Importantly, Crz1*-CLASP did not cause increased gene expression in the absence of light, indicating that CLASP was able to successfully sequester the nuclearly localized Crz1* outside of the nucleus in the dark (Figure S7E).

We next identified six Crz1 gene targets (Yps1, Ena1, Mep1, Put1, Cmk2, Gyp7) for follow up studies. We used the promoters of these genes, which have also been canonically used in the literature (Stathopoulos and Cyert, 1997; Yoshimoto et al., 2002), to build YFP-expressing promoter fusions, each in a strain with Crz1*-CLASP tagged with mCherry for visualization (Figure 3A). We subjected these cells to two distinct types of inputs that mimic natural Crz1 translocation: 2 minute short repeated pulses with different periods or one continuous pulse of varying duration (Figure 3A). We checked that extended light exposure did not cause a growth defect in the Crz1 overexpression strain (Figure S7G). We then measured the nuclear enrichment of mCherry-tagged Crz1*-CLASP continuously at 30 second intervals. We also measured gene expression from all six YFP promoter fusions at 5 hours for all inputs given (Figure 3A). Every input (pulsatile or continuous) generated a given nuclear occupancy, which we calculated as the integral of the measured Crz1*-CLASP nuclear enrichment time traces. A given nuclear occupancy was associated with a commensurate gene expression value (measured at five hours), and these values were plotted against each other for the two input regimes for each of the six promoters. The resulting plot for all nuclear occupancy values are referred to as the Gene Output - Nuclear Occupancy plot (Output-Occupancy plot for short). Exploration of gene expression as a function of nuclear occupancy allowed a comparison on equal footing of the overall integrated responses to pulsed and continuous inputs. The Crz1-responsive promoters showed a spectrum of qualitative and quantitative behaviors in the Output-Occupancy plots (Figure 3A-C, Figure S8). For pGYP7-YFP, like the promoters shown in Figure 2, a pulsed input generated lower gene expression output than a continuous input of the same nuclear occupancy for all values tested, a phenotype that we termed efficient response to continuous inputs (Figure 3B). For pCMK2-YFP, pulsed and continuous inputs generated almost identical gene expression output. However, for pYPS1-YFP, pulsed inputs produced higher gene expression output at all Crz1*-CLASP nuclear occupancy values tested, a phenotype that we termed efficient response to pulsed inputs. These phenotypes were qualitatively reproducible despite slight quantitative day to day variability in gene expression between experiments (Figure S8). The difference in output between pulsed and continuous input as a function of nuclear occupancy was quantified as the ratio of slope of the two lines in the Output-Occupancy plot, termed the slope ratio (Figure 3A). This metric showed that the six Crz1 responsive promoters spanned a range that is bracketed by pYPS1-YFP (slope ratio > 1) and pGYP7-YFP (slope ratio <1), going from a more efficient response to pulsed than continuous inputs to the opposite phenotype (Figure 3C). These phenotypes must reflect different promoter properties since all promoter fusions generated the same YFP as the protein output. We next turned to data-backed computational modeling to systematically explore and interpret these behaviors.

**Figure 3.**
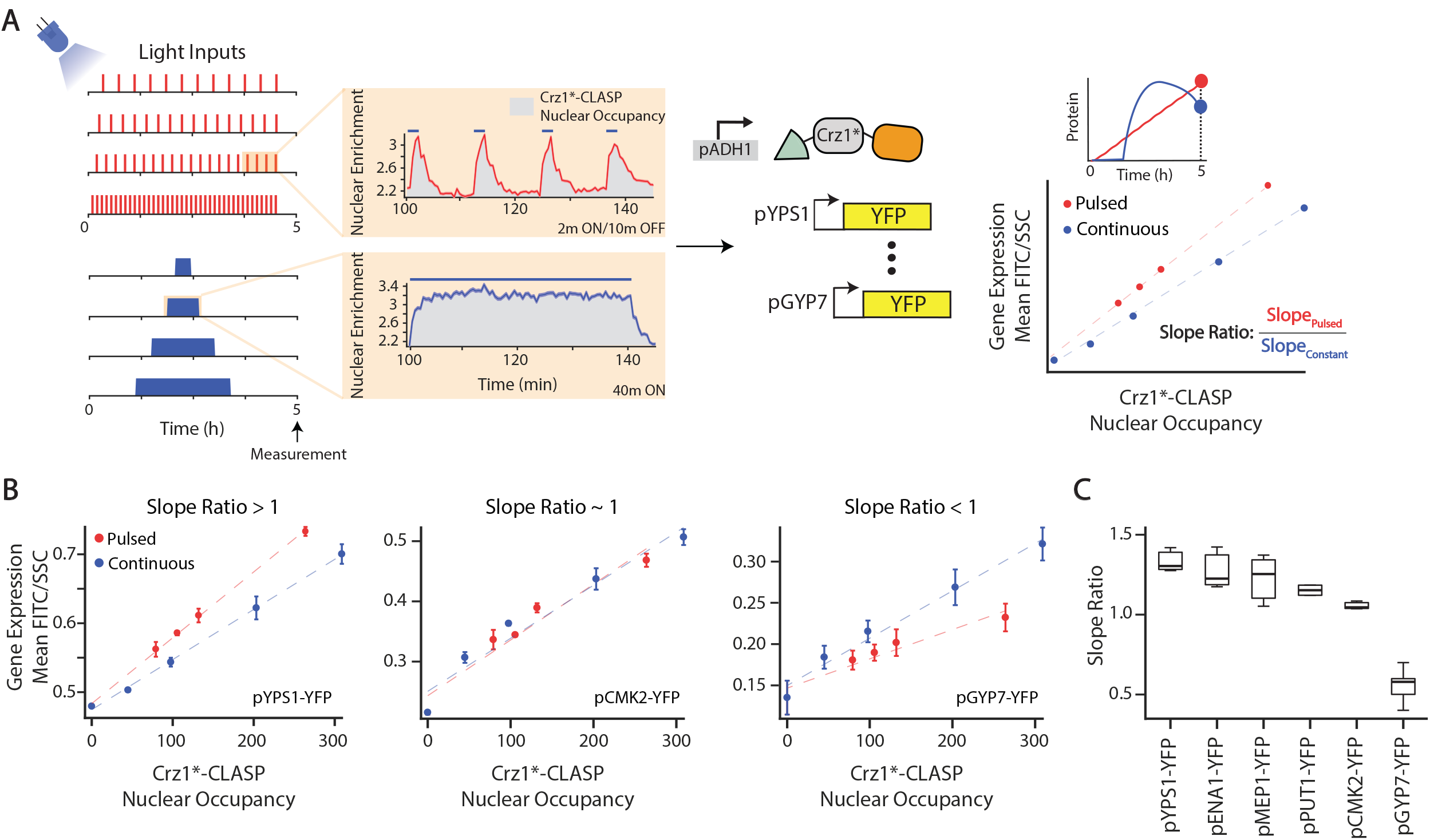
Crz1 target genes show differing interpretation to Crz1*-CLASP short nucleoplasmic pulses. **A)** Schematic of experimental setup used. Two types of light inputs are given to cells expressing Crz1*-CLASP: 2 minute pulses with increasing period (20, 15, 12, and 6 minute periods) and single pulses with increasing duration (20, 40, 80, 120 minutes). Light-induced Crz1*-CLASP nuclear localization is measured with fluorescence microscopy. The mean of single cell fluorescences are plotted (solid red for pulsed input or blue line for continuous inputs), with the shaded area representing 95% confidence interval (red or blue shading). Crz1*-CLASP nuclear occupancy (x-axis in rightmost panel) is quantified as the area under the nuclear localization traces (gray shading in middle panel). Gene expression (mean FITC/SSC) is measured for 6 promoter fusions of target gene driving a fluorescent protein (YFP) at 5 hours after light input. A schematic shows gene expression values for different light input regimes are plotted as a function of nuclear occupancy, generating the Output-Occupancy plot referred to in the text. Each point in the plot is an endpoint measurement of gene expression, as highlighted by the YFP time course schematic above. Red circles represent output-occupancy for short 2 minute pulses with increasing period, and blue circles represent that for continuous single pulse with increasing durations. A best fit line (red for pulsed inputs and blue for continuous inputs) is fit through the data points for the pulsed and continuous inputs. For each Output-Occupancy plot we define the slope ratio as the ratio of the slope of the pulsed to continuous best fit lines. **B)** Output-Occupancy plot for three representative Crz1 target promoters pYPS1-YFP, pCMK2-YFP, and pGYP7-YFP. The error bars are standard deviation of at least 3 biological replicates. **C)** Slope ratios for 6 Crz1 target genes plotted in order of highest to lowest slope ratio. In all panels, Crz1*-CLASP is induced with a 512 a.u. light input.

### Efficient response to short pulses by promoters can be explained by a simple model with fast promoter activation and slow shut-off

To understand the general determinants of the behavior of the Crz1-responsive promoters, we first built a simple and parsimonious model of gene expression. The model consisted of a promoter that occupied two states, OFF (p_off_) and ON (p_on_), with the rate constants k_on_ and k_off_ describing the transition between the two states. The rate of promoter activation depended on the concentration of nuclear TF. The activated promoter then produced YFP mRNA at rate β_1_ and β_0_, which represented the basal activity of the promoter, and the YFP mRNA produced the YFP protein at rate β_2_. mRNA and protein degraded with rates γ_1_ and γ_2_, respectively (Figure 4A). Since the data represented a YFP promoter fusion, the protein degradation rate (γ_2_), protein production rate (β_2_) and mRNA degradation rate (γ_1_) were assumed to be the same for all promoters. Hence, all remaining parameters that differed among the different promoters were only related to promoter properties. Moreover, γ_1_ and γ_2_ were set to plausible values corresponding to YFP characteristics from the literature (Christiano et al., 2014; Wang et al., 2002). The model was simulated with the same pulsatile or continuous nuclear localization input regimes used in the experiments and the output was the YFP protein value at 5 hours, mimicking the experimental setup and data collection procedures.

**Figure 4.**
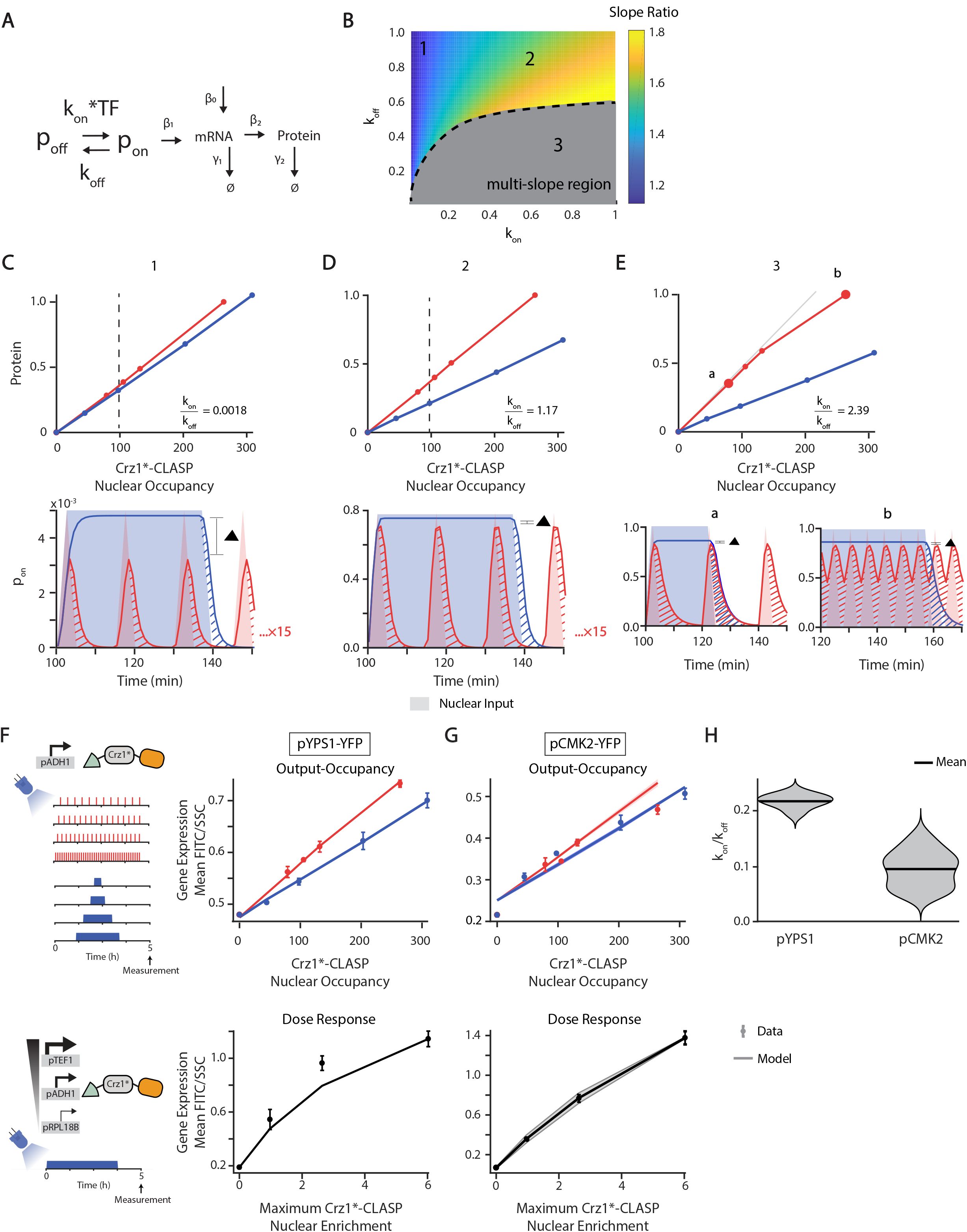
Efficient response to short pulses by promoters can be explained by a simple model with fast promoter activation and slow shut-off. **A)** Schematic of a two-state promoter model, where the input is Crz1*-CLASP nuclear localization (TF) and the output is fluorescent protein level (Protein). The promoter turns ON with rate constant k_on_ and OFF with rate constant k_off_. **B)** Heatmap of slope ratio (defined in Figure 3 and main text) resulting from the model in (A) as a function of k_on_ and k_off_, which both vary from 0.0001-1. The ratio k_on_/k_off_ increases in a clockwise direction on the heatmap. Points 1 and 2 highlight the increasing slope ratio in the linear region. Point 3 resides in the multi-slope region (gray) of the Output-Occupancy plot, and hence is not quantified by slope ratio. In this heatmap, β_1_ varies from 0.0001-10, β_0_ from 0.000001-0.01 and β_2_ from 0.0001-10. The parameter γ_1_ is set to 0.05 and γ_2_ to 0.0083. **C-E) (upper panels)** Output-Occupancy plots generated by the model for different parameter sets that correspond to points 1, 2 and 3 in the heatmap of panel B. The slope ratio for point 1 is 1.1 with a k_on_/k_off_ equal to 0.0018 (C). The slope ratio for point 2 is 1.73 with a k_on_/k_off_ equal to 1.17 (D). The kon/koff for point 3 is 2.39 (E). **(Lower panels)** Examples of time courses of promoter activity (p_on_) for a light input that produces the equivalent of 40 minutes (dotted line in upper panel) in nuclear localization either continuously or in short pulses. Solid lines are the p_on_ pulses while shading denotes nuclear localization. The red and blue hashes represent residual promoter activity beyond the nuclear localization input. The red residual promoter activity is repeated 15 times while the blue residual activity is repeated one time. The ▲bar denotes the difference between the amplitudes generated by the 2 minute pulsed and 40 minute continuous inputs. For panel (E), p_on_ is plotted for two nuclear localization values (denoted **a** and **b** in upper panel) to illustrate a regime in which p_on_ does not completely shut off between repeated pulses. **F) (upper panel)** Output-Occupancy plots for pYPS1-YFP. Circles are experimentally measured values. Solid lines are the mean of the 15 parameter sets that fit the data and shaded areas are the standard deviation of the mean. **(lower panel)** Parameters that fit the Output-Occupancy are used to predict the dose response of pYPS1-YFP (solid black line and gray shading are the mean and standard deviation, respectively, generated by the model). The gray circles are the experimentally measured dose response. **G)** Same as in (F) but repeated for pCMK2-YFP with results from 489 parameter sets that maximize fit to the experimental data. Dose response model predictions are for these parameter sets. **H)** k_on_/k_off_ ratio of the parameter sets that fit both the Output-Occupancy and dose response data for pYPS1-YFP and pCMK2-YFP. The black line denotes mean and the gray cloud denotes the distribution of the k_on_/k_off_ parameter values.

To explore a wide range of parameter regimes, we randomly sampled 10000 parameters (k_on_, k_off_, β_1_, β_2_, β_0_) and generated Output-Occupancy plots and the slope ratio metric for every combination. A plot of slope ratio as a function of each sampled parameter set showed that there was no relationship between β_1_, β_2_ or β_0_ and slope ratio (Figure S9A). Instead, β_1_, β_2_, and β_0_ scaled the range of the output curves. We therefore focused on the influence of k_on_ and k_off_ on slope ratio, plotting the values of that metric as a heatmap in the k_on_−k_off_ plane (Figure 4B). Broadly, there were two clear patterns. First, all parameter combinations tested produced higher gene expression output from pulsed inputs than continuous inputs for all nuclear occupancy values (slope ratio was always >= 1) (Figure 4A-E). Second, the separation between the continuous and pulsed input curves in the Output-Occupancy plots was dictated by the values of k_on_ and k_off_, with the separation increasing with increasing k_on_/k_off_. At very large k_on_/k_off_, the pulsed output became multi-sloped (gray region in Figure 4B).

Since the translation and degradation rates in this model were fixed, the observed differences at the protein level were determined by the differences in the promoter activity, p_on_, which in turn was dictated by k_on_ and k_off_. Hence, to gain intuition about the model results, we focused on the promoter activity, p_on_, in three regimes marked 1, 2, and 3 in Figure 4B, which provided snapshots of promoter activity as k_on_ increased and k_off_ decreased. In the first regime (1), k_on_ was much smaller than k_off_ so that a pulsed input did not induce full ON switching of the promoter within the duration of a short pulse, while p_on_ could reach its maximum possible value during a continuous longer input. This caused an amplitude difference in p_on_ between pulsed and continuous inputs (Figure 4C). However, a relatively slower k_off_ implied that the promoter stayed ON for a period of time beyond the duration of the input, and when repeated for every pulse, this residual activity could counteract the amplitude deficiency (Figure 4C, hatched region under p_on_ curves). Hence, in this k_on_/k_off_ regime, the protein output of the short pulsed input was slightly higher than that of the continuous input. In the second regime (2 in Figure 4B), a larger k_on_ caused the promoter to turn fully ON within one short pulse, and a small k_off_ again caused the promoter to switch OFF slowly, maximizing the gain from every pulse (Figure 4D). As a result, the cumulative promoter activity generated by a short pulsed input was large, and the difference in protein output generated by a pulsed compared to a continuous input of matched nuclear occupancy was also large (Figure 4D). In the third regime (3 in Figure 4B), at an extreme where k_on_ ≫ k_off_, the promoter was ON for the whole duration of the short pulsed input (4 hours of repeated pulses of 2 minutes each, for example), with little OFF switching of the promoter during repeated pulses. On the other hand, a continuous input, with the same cumulative nuclear residence as the short pulsed input, would generate a promoter that was ON approximately only for the duration of the pulse plus the switching off time of the promoter (Figure 4E, region b lower panel). The change in the slope of Output-Occupancy plot occurred when the promoter did not shut completely OFF between two consecutive pulses (compare plots for point (a) and (b) in Figure 4E upper and lower panel). While the model identified this regime, which could be applicable to some promoters, none of the promoters we explored showed strong multi-slope Output-Occupancy curves. Specifically, pYPS1-YFP showed a clear Output-Occupancy relationship with one slope, making it likely that this promoter operated in the k_on_/k_off_ regime in regime 2.

To test whether the model could provide a quantitative fit to the pYPS1-YFP and pCMK2-YFP data, we further fit the parameters to the Output-Occupancy data, specifying fits to be model predictions that maximized the fit through the data points within the error bars for pYPS1-YFP (15 parameter sets) and for pCMK2-YFP (489 parameter sets) (Figure 4F-G). We then subjected these fits to cross-validation, asking whether the parameters with the accompanying model could reproduce the dose response of pYPS1-YFP and pCMK2-YFP. We measured these dose responses in an independent experiment using strains in which Crz1*-CLASP was expressed at different levels using a suite of constitutive promoters of different strengths (pRPL18B, pADH1, and pTEF1; Lee et al., 2015). These strains, which also harbored either pYPS1-YFP or pCMK2-YFP, were subjected to continuous light input over four hours, leading to maximum nuclear localization in all strains. Different strains had different amounts of nuclear Crz1*-CLASP when localized with light, therefore measurement of YFP in each strain provided a different point on the dose response (Figure 4F, G). We subjected the model to the same treatment in silico, and produced computational predictions of the dose response curves for all parameter sets that fit the Output-Occupancy data for pYPS1-YFP or pCMK2-YFP (Figure 4F, G, H). Computational and experimental predictions were in strong qualitative agreement.

Taken together, our data show that a simple promoter model can provide a straightforward scenario in which a promoter can respond more efficiently to repeated short nuclear pulses of a TF than a continuous input. A tight iteration of modeling and experiment further revealed that this behavior is dependent on the dynamics of promoter activity produced by fast promoter activation coupled with its slow inactivation. Fast promoter activation leads to complete activation during a short pulse, and slow promoter turnoff leads to accumulation of expression over the course of repeated pulses, providing insight into how even simple promoters can readily decode dynamic inputs.

### Efficient response to continuous inputs by promoters can be explained by a model with a thresholded transition between non-transcribing promoter states

The simple model from the previous analysis could not produce the pGYP7-YFP phenotype (Figure S10A-B). When the output difference between the pulsed and continuous inputs was small (k_on_ ≪ k_off_) in this model, the output of the pulsed input was always higher than the continuous input. This was because while decreasing k_on_ reduced the output of the pulsed input, it also reduced the dynamic range of the output in response to a continuous input to a point where k_on_ was so small that the promoter was barely activated and the much faster k_off_ quickly shut off promoter activity, resulting in a promoter that was essentially unresponsive to either inputs (Figure S9B).

In order to identify a minimal model that explained the pGYP7-YFP phenotype, we explored eight elaborations of the simple promoter switching model from Figure 4 using a sequence of fitting and cross-validation. In this process, each model was first fit to the Output-Occupancy data in Figure 3; one of the eight models failed to fit. Models that fit the Output-Occupancy data were further fit to the dose response of pGYP7-YFP, which was collected in the same way as for pCMK2 and pYPS1. The pGYP7-YFP dose response was remarkably linear, and four models failed to fit it (Figure 5A-D). For the 3 remaining models, the dose response data served to further constrain parameter sets. For those refined parameters, we cross-validated the models on the data from an additional experiment in which we expressed Crz1*-CLASP from a stronger promoter (pTEF1 versus pADH1), and measured gene expression following a cumulative light induction of 40 minutes administered either as pulsed or continuous input. Following these rounds of fitting and cross-validation (Figure S10C-R), only two of the models surveyed were able to explain all the data we collected (Figure 5A-D, Figure S10O).

**Figure 5.**
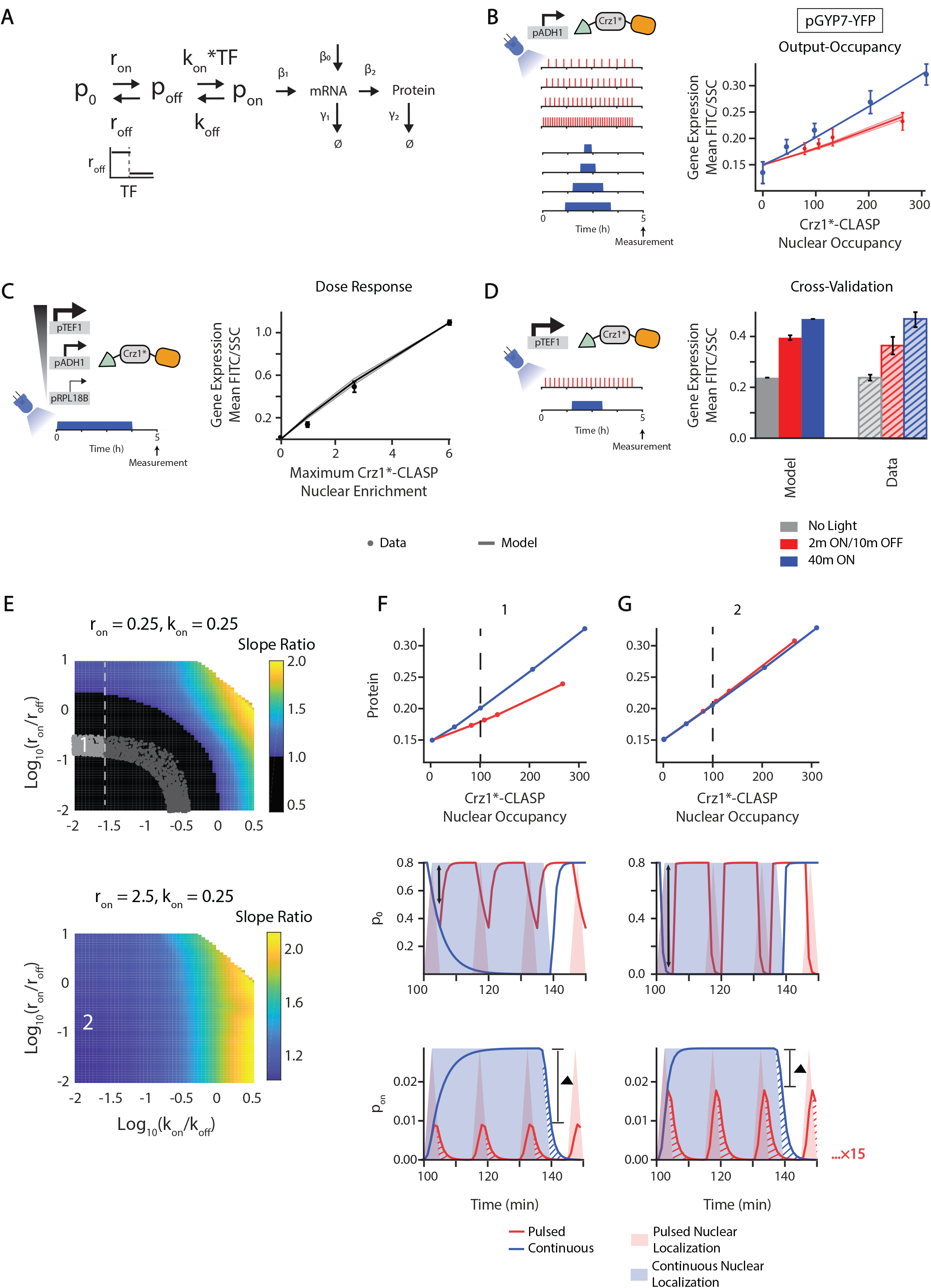
Efficient response to continuous inputs by promoters can be explained by a model with a thresholded transition between non-transcribing promoter states. **A)** Schematic of the three-state model where r_off_, the inactivation rate constant from p_0_ to p_off_, is thresholded by TF concentration and where the activation from p_off_ to p_on_ is linearly dependent on TF. **B) (left panel)** Schematic of experimental setup. **(right panel)** Output-Occupancy plot for pGYP7-YFP. Circles are experimentally measured values while lines denote the mean model output for 96 parameter sets that fit the data points within the error bars, the same metric as used in Figure 4. The solid line denotes the mean and shaded areas denote the standard deviation of the model outputs for these parameter sets. Parameters were sampled (r_on_ from 0.1-100, r_off_ from 0.1-100, k_on_ from 0.0001-1, k_off_ from 0.0001-1, β_1_ from 0.0001-10, β_0_ from 0.000001-0.01, threshold from 0-0.5) or set (β_2_ = 0.06, γ_1_ = 0.05, γ_2_ = 0.0083). **C) (left panel)** Schematic of experimental setup. **(right panel)** Dose response plot for pGYP7-YFP. The parameters that fit the Output-Occupancy data were used to further fit the dose response of pGYP7-YFP using a least squared error criterion (25 parameter sets). Solid black line is the mean generated by the model. The gray circles are the experimentally measured dose response. **D) (left panel)** Schematic of experimental setup. **(right panel)** The parameters that fit the Output-Occupancy are subjected to cross-validation using an experiment where Crz1*-CLASP expression is increased (expressed from a pTEF1 promoter), and cells are exposed to either short (2 minutes ON/10 minutes OFF) or continuous input (40 minutes of light). The model generated outputs (solid gray, red, and blue bars) are plotted with the experimental data (dashed gray, red, and blue bars). The gray bars correspond to no light input. **E) (top panel)** Heatmap shown in the log_10_(k_on_/k_off_)-log_10_(r_on_/r_off_) plane of slope ratio of Output-Occupancy relationship resulting from the model in (A). Parameters are sampled (r_off_ from 0.0025-25, k_off_ from 0.0025-25) or set (r_on_ =0.25, k_on_ = 0.25, β_1_ = 0.0001, β_2_ =0.06, γ_1_ =0.05, γ_2_ =0.0083, threshold= 0.5, β_0_ =0.000001). Point 1 highlights a parameter set that fits the output-occupancy, dose response, and cross-validation datasets. Black region is where slope ratio < 1. Gray dotted line indicates when log_10_(k_on_/k_off_) ∼= −1.5, at which point the dose response changes from linear to nonlinear with increase in the log_10_(k_on_/k_off_) value. All parameters that show a qualitative fit to Output-Occupancy data are displayed as light and dark gray dots. The light gray dots represent parameter sets where all pGYP7-YFP data are quantitatively fit. **(bottom panel)** Heatmap of slope ratio as in (B) with a r_on_ = 2.5, 10 times larger than than in (B). k_on_ is also set to 0.25. Parameters are sampled (r_off_ from 0.025-250, k_off_ from 0.0025-25) or set (β_2_ = 0.0001, β_2_ = 0.06, γ_1_ = 0.05, γ_2_ = 0.0083, threshold = 0.5, β_0_ = 0.000001). Point 2 highlights the effect of increasing both r_on_ and r_off_ while maintaining the ratio log_10_(r_on_/r_off_). **F-G) (upper panels)** Output-Occupancy plots generated by the model for different parameter sets that correspond to points 1 and 2 in the heatmaps of panel E. The slope ratio for point 1 is 0.51 with log_10_(k_on_/k_off_) = −1.58 and log_10_(r_on_/r_off_) = −0.89. The slope ratio for point 2 is with log_10_(k_on_/k_off_) = −1.58 and log_10_(r_on_/r_off_) = −0.89. Point 2 is chosen to highlight the effect of increasing both r_on_ and r_off_ while maintaining the ratio log_10_(r_on_/r_off_). **(middle panels)** Example of a time course of promoter state p_0_ for a light input that produces the equivalent of 40 minutes (dotted line in upper panel) in nuclear localization either continuously or in short pulses. Solid lines are the p_0_ pulses while shading denotes nuclear localization. The black double arrows denote the maximum depletion of the p_0_ state for the pulsed input. **(lower panels)** Example of a time course of promoter activity p_on_ for a light input that produces the equivalent of 40 minutes (dotted line in upper panel) in nuclear localization either continuously or in short pulses, similar to middle panels. The red and blue hashes represent residual promoter activity beyond the nuclear localization input. The red residual promoter activity is repeated 15 times while the blue residual activity is repeated one time. The ▲ bar denotes the difference between the amplitudes generated by the 2 minute pulsed and 40 minute continuous input.

The two models were structurally similar--they both extended the simple two-state model to contain another promoter state, thereby necessitating transition through an unproductive promoter state (p_off_) before the promoter can be fully activated. Therefore, in these models, the first transition occurred reversibly between promoter state p_0_ and a non-transcribing state p_off_ with rate constants r_on_ and r_off_, while a second transition stage occurred between p_off_ and p_on_ with rate constants k_on_ and k_off_. Both models also necessitated a linear dependence on TF in the second transition stage, whose effect was to prevent the dose response from exhibiting a thresholded behavior. Finally, the two models necessitated a thresholded interaction in the first promoter transition stage, but differed in where it was applied -- in one model, r_on_ was a thresholded function of TF, while in the other model, it was r_off_ that was thresholded by TF (Figure 5A, Figure S10O). The threshold on either r_on_ or r_off_ prevented short pulsed inputs from fully transitioning the promoter from the p_0_ state, essentially creating a filter for short inputs. Detailed descriptions of all models and their exploration can be found in the Supplementary Text and Figure S10.

To gain more insight into the pGYP7-YFP phenotype, we further explored the 3-state, r_off_ threshold model for many parameter values (Figure 5A). We sampled the model parameters by fixing r_on_ and k_on_ to values that fit the data from Figure 5B-D and varied r_off_ and k_off_ within a range of four logs. We then generated Output-Occupancy plots for every parameter set and computed its corresponding slope ratio metric, which we plotted in the log_10_(k_on_/k_off_) − log_10_(r_on_/r_off_) plane (Figure 5E). Overall, we found that this model can generate both higher expression with a continuous input (slope ratio <1, black region in Figure 5E, top panel) and higher expression with short pulses (slope ratio> 1, colored region on Figure 5E, top and bottom panel).

Quantitatively, there seemed to be three parameter constraints for this promoter model to respond more efficiently to a continuous input than a pulsed one. First, the rate of transition from p_0_ to p_off_ should be slow; second, r_off_ should be fast relative to r_on_; third, k_off_ should be fast relative to k_on_. An analysis of the 3-state r_on_ threshold model demonstrated similar requirements (Figure S11). When r_on_ and r_off_ were increased tenfold, there were no parameter combinations that generated higher expression for continuous inputs than short pulses (Figure 5E, bottom panel, Figure 5F-G, top panel). The difference in the protein outputs between the pulsed and continuous inputs was determined by the amplitude differences of promoter activity p_on_ (Figure 5F-G, middle panel), which was in turn dictated by the amplitudes of depletion from p_0_ for the short pulsed and continuous inputs (Figure 5F-G, middle panel). A slow transition from p_0_ prevented the quick, full depletion of this state before a short pulse ended, while p_0_ was fully depleted for the continuous input (Figure 5F, middle panel). In contrast, when r_on_ and r_off_ were fast, this difference disappeared as the transition from p_0_ was now able to reach the same maximal amplitude in the duration of the short input (Figure 5G, middle panel). Hence, the incomplete depletion of the p_0_ state in the duration of the short pulsed input accounted for the difference in protein outputs between the short pulsed and continuous inputs.

The requirement that the value of r_off_ be large relative to r_on_ was motivated by the fact that r_off_ dictated how quickly the promoter state transitioned back to the initial OFF state p_0_ after the end of a short pulse. When the value of r_off_ decreased relative to r_on_ (Figure S12A-B), the depletion of p_0_ could proceed to completion during a short pulse (Figure S12B, middle panel), and the resulting maximum amplitudes of the active promoter state p_on_ were comparable for a pulsed or continuous input (Figure S12B, bottom panel). Lastly, as k_off_ was decreased while keeping all other parameters constant, the p_on_ to p_off_ switching also slowed, and promoter activity continued unabated between two pulses, hence maximizing the gain of promoter activity from every input pulse and causing stronger gene expression from pulses than from a continuous input (Figure S12C). This was in essence the same mechanism as described in Figure 4. In summary, slow transition from the initial OFF state to the secondary OFF state prevented the short pulsed input from achieving a quick depletion of the initial OFF state, essentially creating a filter for short inputs. This analysis reveals that differences in dynamic changes in promoter activity can result in more efficient response to a continuous TF input than a pulsed input.

Finally, in addition to the constraints above, we found that a threshold of log_10_(k_on_/k_off_) ∼> −1.5 seemed to demarcate the transition between a linear and nonlinear promoter dose response in the parameter regime probed (light gray points, Figure 5E, left panel), therefore imposing quantitative bounds on this promoter model to exhibit a graded dose response as seen in the data.

## Discussion

In this work, we presented a general optogenetic tool that circumvents some functional caveats of previous methodologies, and used it productively to investigate in a systematic way how promoters are able to differentially respond to pulsed versus continuous transcription factor localization. Our tool, CLASP, was inspired by previous optimization efforts that attempted to make optogenetic control more precise and malleable (Gautier et al., 2010; Shimizu-Sato et al., 2002; Strickland et al., 2012; Toettcher et al., 2010, 2013). However, we pushed our optimization and characterization efforts further, producing a tool that was functional for all cargos tested, had a large dynamic range and no detectable deleterious impact on the cell. We capitalized on this tool to ask a simple and profound biological question -- can genes differentiate between transcription factor inputs that differ only in their dynamic patterns? CLASP allowed us to directly test and provide a definitive demonstration of this phenomenon for a number of TF-promoter pairs. We then explored mechanistic underpinnings of this behavior using the transcription factor Crz1, whose response to calcium stress is naturally pulsatile.

The precise and robust operation of CLASP allowed us to approach this investigation methodically, establishing through rounds of computational modeling and experimentation two classes of models that could explain how Crz1 promoters may respond differentially to different Crz1 pulsing inputs. For promoters that responded more efficiently to short pulsed inputs, we demonstrated that their behaviors could be simply explained by a two-state model of the promoter (ON or OFF), transitioning between them with first order kinetics. If the ON rate, which is dependent on the nuclear TF concentration, is fast and the OFF rate is slow, then the output from short pulses is larger than for a continuous TF input of the same nuclear occupancy. By contrast, for promoters that responded more efficiently to continuous than pulsed TF inputs, a more involved model needed to be invoked. Interestingly, for pGYP7, this behavior coincided with an additional property -- a linear dose response. To explain the efficient response to continuous inputs in isolation, either a thresholded step in a two-state model or a model with additional promoter states was sufficient. To satisfy a linear dose response in isolation, k_off_/k_on_ of a simple two-state model must be relatively large. Yet, in order to satisfy both properties simultaneously, an involved model that fulfilled the requirements for each individual property and that included a dependence on the TF at each promoter stage was needed.

What possible benefits of this differential interpretation, and of the dose response linearity, might exist for the cell? Under stress, Crz1 undergoes an initial long 40-60 minute nuclear localization, followed by pulsing in the “maintenance” phase of the calcium response. It is possible that differential interpretation of these dynamic inputs by different sets of genes is used as a mechanism to temporally program the response, with cohorts of genes activating strongly in the first long pulse and then to a lesser degree with the subsequent pulsatile episode, while others do the opposite. Moreover, it has been observed that Crz1 pulses with different amplitudes in the “maintenance” phase (Cai et al., 2008) and hence a linear dose response may be useful for preserving the efficient response to a continuous input over a range of pulse amplitudes. Previous work that probed whether pulses of Crz1 were frequency modulated as a function of the calcium chloride input found that promoters downstream of Crz1, including pGYP7, respond linearly to increasing levels of calcium chloride and interpreted this linear relationship to be the outcome of the frequency modulation (Cai et al., 2008). In this work, we established that pGYP7 was inherently linear as a function of Crz1 input for a broad range of Crz1 levels (Figure 5). It will be interesting to explore how much the linearity observed in response to calcium chloride can be attributed to frequency modulation, or alternatively, ask whether it can be solely explained by the linearity of the promoter itself.

Our studies presented concise phenomenological promoter models which were filtered through rounds of cross validation and explained all the data collected. These models correspond to well-understood mechanisms of promoter regulation. The 2-state kinetic model that describes the pYPS1-YFP behavior may represent a simple promoter activated by a TF. The two promoter models that describe pGYP7-YFP, in which a thresholding step in promoter transition occurs either in the ON (r_on_) or the OFF (r_off_) rates, can represent more complex biological mechanisms. The 3-state promoter, on the other hand, can correspond to closed (p_0_), open but non-transcribing (p_off_), and open and transcribing (p_on_) states of the promoter. Additionally, the TF-thresholded transitions of r_on_ can represent transcription factor activation of chromatin remodelers (Dillon and Festenstein, 2002; Spitz and Furlong, 2012), while thresholding in the r_off_ rate constant can represent transcription factor inhibition of heterochromatin formation either by physical occlusion of nucleosomes (Lickwar et al., 2012; Platt et al., 2013) or inhibition of deacetylation of nucleosomes (Cheng et al., 2011; Gaupel et al., 2014; Steinfeld et al., 2007). Available nucleosome occupancy data for Crz1 target genes support the models that describe pYPS1-YFP and pGYP7-YFP by showing a negative correlation between nucleosome occupancy and responsiveness to short pulses in our data, with genes that respond efficiently to short pulses exhibiting lower nucleosome occupancy (Figure S13). This correlative data aligns with the idea that a more efficient response to continuous inputs requires additional promoter regulation, such as a TF-gated promoter transition between non-transcribing promoter states, compared to promoters that respond efficiently to short inputs. Still, more mechanistic studies, such as RNA FISH for observation of promoter dynamics, are needed to pinpoint the biochemical mechanisms that underlie these models.

While we used CLASP to explore fundamental aspects of promoter dynamics, many investigations that extend this work in broader directions are readily possible. For example, we showed that CLASP can sequester and translocate basally nuclear TFs (Figure S6), and therefore can be used with a variety of cargos to mimic dynamic TF knockouts in the presence of their activating environmental inputs, or combinatorial TF regulation using a leave-one-out strategy. More generally, CLASP is likely transplantable to other cells and organisms. In mammalian cells, NFAT, the Crz1 homolog, is known to respond to immunological stimuli with nuclear pulsing (Kar et al., 2016). We envision using CLASP in this context to explore whether NFAT target genes also use differential decoding to modulate different expression programs and determine general conserved principles.

Finally, while our studies focused on transcriptional regulation at the promoter level, many opportunities for further decoding of TF information can be implemented through modulation and control of translation and degradation of mRNA and protein (Figure S9C). It will be fascinating to study the bounds of complexity explored by endogenous genes through combinatorial tuning of all steps of gene expression to implement sophisticated dynamic decoding capabilities.

## Acknowledgements

This work was supported by NIH grant R01GM119033 (awarded to H.E-.S), the National Science Foundation (NSF) Graduate Research Fellowship (GRF) (awarded to S.Y.C.), National Defense Science & Engineering Graduate (NDSEG) Fellowship, Paul and Daisy Soros Fellowship for New Americans, and National Institute for General Medical Sciences (NIGMS) Initiative for Maximizing Student Development (IMSD) Fellowship (awarded to L.O). H.E-.S is an investigator in the Chan Zuckerberg Biohub.

## Author Contributions

S.Y.C., L.C.O., and H.E-.S. conceived of the study. S.Y.C., L.C.O., A.H.N., and J.S-.O. constructed parts. L.J.B. designed the Optoplate. S.Y.C. and L.C.O. collected data, and S.Y.C., L.C.O., and L.T.N. processed data. S.Y.C., L.C.O., M.C., and H.E-.S. interpreted results, wrote and edited the manuscript.

## Methods

### Contact for reagent and resources sharing

Further information and requests for resources and reagents should be directed to and will be fulfilled by the Lead Contact, Hana El-Samad. To request reagents, please submit a form to UCSF at https://ita.ucsf.edu/researchers/mta. All plasmids will also be deposited on Addgene and can be requested from there.

### Experimental model and subject details

#### Saccharomyces Cerevisiae

##### Plasmid and strain construction

Hierarchical golden gate assembly was used to assemble plasmids for yeast strain construction using the method in Lee et al. BsaI, BsmBI, and NotI cut sites were removed from individual parts to facilitate downstream assembly and linearization. Parts were either generated via PCR or purchased as gBlocks from IDT. These parts were then assembled into transcriptional units (promoter-gene-terminator) on cassette plasmids. These cassettes were assembled together to form multi-gene plasmids for insertion into the yeast genome at the TRP, URA, or LEU locus. Cassettes were digested with NotI and then transformed into yeast as described in Lee S et al., 2013 or Lee ME et al., 2015.

##### Yeast strains, media, and growth conditions

The base *S. cerevisiae* strain used for experimentation was W303α or BY4741. Base strain for each engineered strain is noted in the strain list. From these base strains, knockout of endogenous transcription factors was done with a one-step replacement using a plasmid that contains 40 base pair overlaps in the 5’ and 3’ UTR of the transcription factor (Gardner and Jaspersen, 2014). The 40 base pair overhangs flank the *Candida Albicans* HIS selectable marker.

Single colonies were picked from auxotrophic SD (6.7 g/L Bacto-yeast nitrogen base without amino acids, 2 g/L complete supplement amino acid mix, 20 g/L dextrose) agar plates. For microscopy and growth measurement studies, colonies were picked into 1 ml SDC media. For flow cytometry studies, colonies were picked into 1 ml YPD (yeast extract, peptone, 2% glucose) or SDC (6.7 g/L Bacto-yeast nitrogen base without amino acids, 2 g/L complete supplement amino acid mix, 20 g/L dextrose) media. Colonies were grown overnight from 30°C to saturation. Prior to the start of an experiment, cells were diluted into 1-3 ml of SDC and grown for 4 hours to an OD of 0.05-0.1 prior to the start of an experiment. A TECAN Spark 10M plate reader (TECAN, Mannedorf, Switzerland) was used for growth measurements.

### Method details

#### Microscopy and blue light delivery

Cells were imaged in 96-well Matriplates (MGB096-1-2-LG-L; Brooks Life Science Systems, Spokane, WA). For widefield microscopy, blue light optogenetic stimulation of samples was done using a custom built “optoPlate” as described in Bugaj et al (Bugaj et al., 2018). Individually addressable LEDs (in 96-well format) were controlled by an Arduino Micro microcontroller and programmed with different dynamic light patterns using custom Arduino scripts. Custom adapters for fitting optoPlate on to 96-well matrix plates were designed in AutoCad and 3D printed. For confocal microscopy, blue light stimulation was done using GFP laser illumination. A Nikon Ti inverted scope, with mercury arc-lamp illumination using RFP (560/40 nm excitation, 630/75 nm emission; 572/35 nm excitation, 632/60 nm emission; both manufactured by Chroma, Bellows Falls, VT) and near-infrared FP (640/30 nm excitation, 700/35 nm emission; Chroma, Bellows Falls, VT) filters, was used for widefield microscopy imaging of samples. Images were taken with an Andor EMCCD camera. Automated imaging was controlled and coordinated by custom Matlab (MathWorks, Natick, MA) software interfaced with the μmanager software suite (Edelstein et al., 2010). Confocal microscopy of samples took place on a Nikon Ti inverted scope with a Yokogawa CSU-22 spinning disk confocal scanner unit; cells were excited with laser illumination for Cy3 (561 nm, 100 mW Coherent OBIS; ET610/60nm emission filter) and Cy5 (640 nm,100 mW Coherent OBIS; ET700/75nm emission filter). Imaging was controlled with Nikon Elements 5.02 build 1266 (Nikon Instruments, Melville, NY).

#### Quantification of nuclear localization

The software tool “ilastik” was used for image segmentation to determine nuclear occupancy (Sommer et al., 2011). Time lapse images of the IRFP nuclear marker were used to identify nuclei objects. Nuclear occupancy of each nucleus was defined as the mean pixel intensity of this nucleus. Cell tracking of nuclear/cytoplasmic enrichment was done using automated yeast cell tracking software implemented in Matlab (Doncic et al., 2013). Photobleaching was corrected for cells that underwent constant illumination and frequent imaging. This correction was done by fitting an exponential decay function to each nuclear and cytoplasmic trace and then dividing each trace by its decay function. For all microscopy analysis, “nuclear/cytoplasmic enrichment” represented the mean pixel intensity of the nucleus divided by the mean pixel intensity of the cytoplasm, and “fold change” represented division by the value of the signal at t=0. Nuclear localization duration was defined as the time from light ON to the time when the nuclear/cytoplasmic enrichment has returned to 75% of the starting (t=0) value. Normalized peak-trough difference was quantified across all pulses for single cell traces, and represented the difference between the local maximum (the peak) and the local minimum (the trough) values divided by the maximum peak-trough difference in the population-averaged traces.

#### Flow cytometry

Analysis of fluorescent protein reporter expression was performed with a BD LSRII flow cytometer (BD Biosciences) equipped with a high-throughput sampler. For steady-state measurements, cultures were diluted in TE before running through the instrument. Cultures were run on the instrument 1 hour (+/− 20 min) after optical stimulation using the optoPlate, to allow for YFP maturation. YFP (Venus) fluorescence was measured using the FITC channel and RFP (mCherry/mScarlet) was measured using the PE-Texas Red channel. For steady-state measurements, a maximum of 10,000 events were collected per sample. Fluorescence values were calculated using the height (H) measurement for the appropriate channel and normalized to cell size by dividing by side scatter (SSC-H). All analysis of flow cytometry data was performed in Python 2.7 using the package FlowCytometryTools and custom scripts.

#### Growth Assays

Growth was measured using a TECAN Spark 10M plate reader (TECAN, Mannedorf, Switzerland) using 600nm excitation. Cultures were plated into Corning 3904 96-well assay plates (Corning, Corning, NY) and grown at 30°C while shaking until saturation. Data was analyzed in Python 2.7 or Matlab using custom scripts. To quantify log phase growth rate, only the OD600 measurements which were between .1 and 1 for each strain were used. A linear regression was then fit to the natural logarithm of the log phase OD600 values (as y) and time (as x). The slope from this regression was plotted as the log phase growth rate.

#### Computational modeling

Ordinary differential equation (ODE) models of gene expression focusing on promoter kinetics were constructed. For the simple kinetic model that described an efficient response to short pulses, a model was constructed with three state variables and seven parameters. Nine models were constructed and tested for efficient response to continuous pulses. These models either contained three or five state variables with up to ten parameters. Latin hypercube sampling was done to randomly sample parameters. ODE solvers 45 and 113 in Matlab were used. Least squared error and fit within the error bars of the data were metrics used to obtain model fits. Additional details of the computational methods are described in the Supplementary Text.

#### Treatment with CaCl_2_ stress

Cells were grown at 30°C in YPD medium to saturation overnight. Cells were then diluted prior to the start of an experiment and grown for 4 hours to an OD of 0.05-0.1. For microscopy experiments, cells were plated in SDC with concanavalin A (conA) for 15 minutes to adhere them to the bottom of the glass imaging plate. Prior to imaging, the SDC was removed and replaced with a solution of SDC with 0.2M CaCl_2_. For flow cytometry experiments, cells in SDC were diluted to OD 0.1 in a media of SDC with 0.2M CaCl_2_ and grown in the media for the duration of the experiment. Prior to measurement, the 0.2M CaCl_2_ media was removed by centrifugation with 3 washes in 1X TE.

## References

1. AkhavanAghdam, Z., Sinha, J., Tabbaa, O.P., and Hao, N. (2016). Dynamic control of gene regulatory logic by seemingly redundant transcription factors.

2. Batchelor, E., Loewer, A., Mock, C., and Lahav, G. (2011). Stimulus-dependent dynamics of p53 in single cells. Mol. Syst. Biol. 7, 488.

3. Cai, L., Dalal, C.K., and Elowitz, M.B. (2008). Frequency-modulated nuclear localization bursts coordinate gene regulation. Nature 455, 485–490.

4. Cheng, C., Shou, C., Yip, K.Y., and Gerstein, M.B. (2011). Genome-wide analysis of chromatin features identifies histone modification sensitive and insensitive yeast transcription factors. Genome Biol. 12, R111.

5. Chi, Y., Huddleston, M.J., Zhang, X., Young, R.A., Annan, R.S., Carr, S.A., and Deshaies, R.J. (2001). Negative regulation of Gcn4 and Msn2 transcription factors by Srb10 cyclin-dependent kinase. Genes Dev. 15, 1078–1092.

6. Chong, Y.T., Koh, J.L.Y., Friesen, H., Duffy, S.K., Duffy, K., Cox, M.J., Moses, A., Moffat, J., Boone, C., and Andrews, B.J. (2015). Yeast Proteome Dynamics from Single Cell Imaging and Automated Analysis. Cell 161, 1413–1424.

7. Christiano, R., Nagaraj, N., Fröhlich, F., and Walther, T.C. (2014). Global Proteome Turnover Analyses of the Yeasts S. cerevisiae and S. pombe. Cell Rep. 9, 1959–1965.

8. Covert, M.W., Leung, T.H., Gaston, J.E., and Baltimore, D. (2005). Achieving stability of lipopolysaccharide-induced NF-kappaB activation. Science 309, 1854–1857.

9. Czyz, M., Nagiec, M.M., and Dickson, R.C. (1993). Autoregulation of GAL4 transcription is essential for rapid growth of Kluyveromyces lactis on lactose and galactose. Nucleic Acids Res. 21, 4378–4382.

10. Dalal, C.K., Cai, L., Lin, Y., Rahbar, K., and Elowitz, M.B. (2014). Pulsatile Dynamics in the Yeast Proteome. Curr. Biol. CB 24, 2189–2194.

11. Dillon, N., and Festenstein, R. (2002). Unravelling heterochromatin: competition between positive and negative factors regulates accessibility. Trends Genet. TIG 18, 252–258.

12. Durchschlag, E., Reiter, W., Ammerer, G., and Schüller, C. (2004). Nuclear localization destabilizes the stress-regulated transcription factor Msn2. J. Biol. Chem. 279, 55425–55432.

13. Garmendia-Torres, C., Goldbeter, A., and Jacquet, M. (2007). Nucleocytoplasmic Oscillations of the Yeast Transcription Factor Msn2: Evidence for Periodic PKA Activation. Curr. Biol. 17, 1044–1049.

14. Gasch, A.P., Spellman, P.T., Kao, C.M., Carmel-Harel, O., Eisen, M.B., Storz, G., Botstein, D., and Brown, P.O. (2000). Genomic expression programs in the response of yeast cells to environmental changes. Mol. Biol. Cell 11, 4241–4257.

15. Gaupel, A.-C., Begley, T., and Tenniswood, M. (2014). High throughput screening identifies modulators of histone deacetylase inhibitors. BMC Genomics 15, 528.

16. Gautier, A., Nguyen, D.P., Lusic, H., An, W., Deiters, A., and Chin, J.W. (2010). Genetically Encoded Photocontrol of Protein Localization in Mammalian Cells. J. Am. Chem. Soc. 132, 4086–4088.

17. Gotoh, Y., Nishida, E., Yamashita, T., Hoshi, M., Kawakami, M., and Sakai, H. (1990). Microtubule-associated-protein (MAP) kinase activated by nerve growth factor and epidermal growth factor in PC12 cells. Identity with the mitogen-activated MAP kinase of fibroblastic cells. Eur. J. Biochem. 193, 661–669.

18. Granados, A.A., Pietsch, J.M.J., Cepeda-Humerez, S.A., Farquhar, I.L., Tkačik, G., and Swain, P.S. (2018). Distributed and dynamic intracellular organization of extracellular information. Proc. Natl. Acad. Sci. 115, 6088–6093.

19. Hansen, A.S., and O’Shea, E.K. (2013). Promoter decoding of transcription factor dynamics involves a trade-off between noise and control of gene expression. Mol. Syst. Biol. 9, 704.

20. Hansen, A.S., and O’Shea, E.K. (2015a). cis Determinants of Promoter Threshold and Activation Timescale. Cell Rep. 12, 1226–1233.

21. Hansen, A.S., and O’Shea, E.K. (2015b). Limits on information transduction through amplitude and frequency regulation of transcription factor activity. ELife 4, e06559.

22. Hao, N., and O’Shea, E.K. (2011). Signal-dependent dynamics of transcription factor translocation controls gene expression. Nat. Struct. Mol. Biol. 19, 31–39.

23. Hao, N., Budnik, B.A., Gunawardena, J., and O’Shea, E.K. (2013). Tunable Signal Processing Through Modular Control of Transcription Factor Translocation. Science 339, 460–464.

24. Hoffmann, A., Levchenko, A., Scott, M.L., and Baltimore, D. (2002). The IκB-NF-κB Signaling Module: Temporal Control and Selective Gene Activation. Science 298, 1241–1245.

25. Khalil, A.S., Lu, T.K., Bashor, C.J., Ramirez, C.L., Pyenson, N.C., Joung, J.K., and Collins, J.J. (2012). A Synthetic Biology Framework for Programming Eukaryotic Transcription Functions. Cell 150, 647–658.

26. Kosugi, S., Hasebe, M., Matsumura, N., Takashima, H., Miyamoto-Sato, E., Tomita, M., and Yanagawa, H. (2009). Six classes of nuclear localization signals specific to different binding grooves of importin alpha. J. Biol. Chem. 284, 478–485.

27. Lee, M.E., DeLoache, W.C., Cervantes, B., and Dueber, J.E. (2015). A Highly Characterized Yeast Toolkit for Modular, Multipart Assembly. ACS Synth. Biol. 4, 975–986.

28. Lickwar, C.R., Mueller, F., Hanlon, S.E., McNally, J.G., and Lieb, J.D. (2012). Genome-wide protein-DNA binding dynamics suggest a molecular clutch for transcription factor function. Nature 484, 251–255.

29. Lin, Y.T., and Doering, C.R. (2016). Gene expression dynamics with stochastic bursts: Construction and exact results for a coarse-grained model. Phys. Rev. E 93, 022409.

30. Nelson, D.E., Ihekwaba, A.E.C., Elliott, M., Johnson, J.R., Gibney, C.A., Foreman, B.E., Nelson, G., See, V., Horton, C.A., Spiller, D.G., et al. (2004). Oscillations in NF-kappaB signaling control the dynamics of gene expression. Science 306, 704–708.

31. Nguyen, T.T., Scimeca, J.C., Filloux, C., Peraldi, P., Carpentier, J.L., and Van Obberghen, E. (1993). Co-regulation of the mitogen-activated protein kinase, extracellular signal-regulated kinase 1, and the 90-kDa ribosomal S6 kinase in PC12 cells. Distinct effects of the neurotrophic factor, nerve growth factor, and the mitogenic factor, epidermal growth factor. J. Biol. Chem. 268, 9803–9810.

32. Niopek, D., Benzinger, D., Roensch, J., Draebing, T., Wehler, P., Eils, R., and Di Ventura, B. (2014). Engineering light-inducible nuclear localization signals for precise spatiotemporal control of protein dynamics in living cells. Nat. Commun. 5, 4404.

33. Pincus, D., Aranda-Díaz, A., Zuleta, I.A., Walter, P., and El-Samad, H. (2014). Delayed Ras/PKA signaling augments the unfolded protein response. Proc. Natl. Acad. Sci. 111, 14800–14805.

34. Platt, J.M., Ryvkin, P., Wanat, J.J., Donahue, G., Ricketts, M.D., Barrett, S.P., Waters, H.J., Song, S., Chavez, A., Abdallah, K.O., et al. (2013). Rap1 relocalization contributes to the chromatin-mediated gene expression profile and pace of cell senescence. Genes Dev. 27, 1406–1420.

35. Purvis, J.E., and Lahav, G. (2013). Encoding and Decoding Cellular Information through Signaling Dynamics. Cell 152, 945–956.

36. Purvis, J.E., Karhohs, K.W., Mock, C., Batchelor, E., Loewer, A., and Lahav, G. (2012). p53 Dynamics Control Cell Fate. Science 336, 1440–1444.

37. Redchuk, T.A., Omelina, E.S., Chernov, K.G., and Verkhusha, V.V. (2017). Near-infrared optogenetic pair for protein regulation and spectral multiplexing. Nat. Chem. Biol. 13, 633–639.

38. Sadeh, A., Movshovich, N., Volokh, M., Gheber, L., and Aharoni, A. (2011). Fine-tuning of the Msn2/4–mediated yeast stress responses as revealed by systematic deletion of Msn2/4 partners. Mol. Biol. Cell 22, 3127–3138.

39. Shimizu-Sato, S., Huq, E., Tepperman, J.M., and Quail, P.H. (2002). A light-switchable gene promoter system. Nat. Biotechnol. 20, 1041–1044.

40. Spitz, F., and Furlong, E.E.M. (2012). Transcription factors: from enhancer binding to developmental control. Nat. Rev. Genet. 13, 613–626.

41. Springer, M., Wykoff, D.D., Miller, N., and O’Shea, E.K. (2003). Partially Phosphorylated Pho4 Activates Transcription of a Subset of Phosphate-Responsive Genes. PLOS Biol. 1, e28.

42. Stathopoulos-Gerontides, A., Guo, J.J., and Cyert, M.S. (1999). Yeast calcineurin regulates nuclear localization of the Crz1p transcription factor through dephosphorylation. Genes Dev. 13, 798–803.

43. Steinfeld, I., Shamir, R., and Kupiec, M. (2007). A genome-wide analysis in Saccharomyces cerevisiae demonstrates the influence of chromatin modifiers on transcription. Nat. Genet. 39, 303–309.

44. Strickland, D., Lin, Y., Wagner, E., Hope, C.M., Zayner, J., Antoniou, C., Sosnick, T.R., Weiss, E.L., and Glotzer, M. (2012). TULIPs: Tunable, light-controlled interacting protein tags for cell biology. Nat. Methods 9, 379–384.

45. Tay, S., Hughey, J.J., Lee, T.K., Lipniacki, T., Quake, S.R., and Covert, M.W. (2010). Single-cell NF-kappaB dynamics reveal digital activation and analogue information processing. Nature 466, 267–271.

46. Toettcher, J.E., Voigt, C.A., Weiner, O.D., and Lim, W.A. (2010). The promise of optogenetics in cell biology: interrogating molecular circuits in space and time. Nat. Methods 8, 35–38.

47. Toettcher, J.E., Weiner, O.D., and Lim, W.A. (2013). Using Optogenetics to Interrogate the Dynamic Control of Signal Transmission by the Ras/Erk Module. Cell 155, 1422–1434.

48. Traverse, S., Gomez, N., Paterson, H., Marshall, C., and Cohen, P. (1992). Sustained activation of the mitogen-activated protein (MAP) kinase cascade may be required for differentiation of PC12 cells. Comparison of the effects of nerve growth factor and epidermal growth factor. Biochem. J. 288 (Pt 2), 351–355.

49. Vardi, N., Levy, S., Gurvich, Y., Polacheck, T., Carmi, M., Jaitin, D., Amit, I., and Barkai, N. (2014). Sequential feedback induction stabilizes the phosphate starvation response in budding yeast. Cell Rep. 9, 1122–1134.

50. Wang, H., Vilela, M., Winkler, A., Tarnawski, M., Schlichting, I., Yumerefendi, H., Kuhlman, B., Liu, R., Danuser, G., and Hahn, K.M. (2016). LOVTRAP, An Optogenetic System for Photo-induced Protein Dissociation. Nat. Methods 13, 755–758.

51. Wang, Y., Liu, C.L., Storey, J.D., Tibshirani, R.J., Herschlag, D., and Brown, P.O. (2002). Precision and functional specificity in mRNA decay. Proc. Natl. Acad. Sci. U. S. A. 99, 5860–5865.

52. Werner, S.L., Kearns, J.D., Zadorozhnaya, V., Lynch, C., O’Dea, E., Boldin, M.P., Ma, A., Baltimore, D., and Hoffmann, A. (2008). Encoding NF-kappaB temporal control in response to TNF: distinct roles for the negative regulators IkappaBalpha and A20. Genes Dev. 22, 2093–2101.

53. Yoshimoto, H., Saltsman, K., Gasch, A.P., Li, H.X., Ogawa, N., Botstein, D., Brown, P.O., and Cyert, M.S. (2002). Genome-wide analysis of gene expression regulated by the calcineurin/Crz1p signaling pathway in Saccharomyces cerevisiae. J. Biol. Chem. 277, 31079–31088.

54. Yumerefendi, H., Dickinson, D.J., Wang, H., Zimmerman, S.P., Bear, J.E., Goldstein, B., Hahn, K., and Kuhlman, B. (2015). Control of Protein Activity and Cell Fate Specification via Light-Mediated Nuclear Translocation. PloS One 10, e0128443.

55. Yumerefendi, H., Wang, H., Dickinson, D.J., Lerner, A.M., Malkus, P., Goldstein, B., Hahn, K., and Kuhlman, B. (2018). Light-Dependent Cytoplasmic Recruitment Enhances the Dynamic Range of a Nuclear Import Photoswitch. Chembiochem Eur. J. Chem. Biol. 19, 1319–1325.

