## SupplementaryMaterials for "Optogenetic control reveals differential promoter interpretation of transcription factor nuclear translocation dynamics"

### Supplementary Text

#### Experimental Controls

##### Measuring the basal and constitutively nuclear gene expression of TFs

To further assess how TF-CLASP-induced expression compares to endogenous gene expression, we measured the level of reporter gene expression when the TFs were constitutively localized to the nucleus by C-terminally tagging them with the same NLS used in yeLANS (TF-NLS), or in their basal localization by C-terminally tagging them with only mScarlet. All TF-NLS, TF-mScarlet, and TF-CLASP constructs were expressed from pRPL18b. We compared this value to expression achieved when TF-CLASP was induced with 2 hours of blue light. SynTF-CLASP achieved 52% of pSYNTF-YFP expression produced through constitutive nuclear localization of SynTF (Figure S5B). Furthermore, the mean SynTF-CLASP-induced gene expression in the dark (.09) was similar to the the mean basal gene expression in a strain in which the SynTF was only tagged with mScarlet (.07) (Figure S5B). Pho4-CLASP activated pPHO84-YFP to 14% of the gene expression achieved with constitutive nuclear localization (Figure S5C) while Msn2-CLASP was more efficient at inducing pHSP12-YFP gene expression than constitutive Msn2 nuclear localization (23% greater expression, Figure S5D). Since Msn2 is subject to faster degradation in the nucleus (Chi et al., 2001; Durchschlag et al., 2004), transient localization with CLASP may be more efficient at inducing gene expression. For both pHSP12-YFP and pPHO84-YFP, reporter expression in the dark was lower in a strain that had either Msn2-CLASP or Pho4-CLASP than in their respective controls with either Msn2 or Pho4 when only tagged with mScarlet (28% and 88% lower, respectively). In fact, pPHO84-YFP showed basal bimodal expression in the constitutively expressed Pho4 strain, but not in the Pho4-CLASP strain (Figure S5C). These data suggest that CLASP can potently sequester TFs in the dark.

#### Modeling

##### Model equations and sampling details of the pYPS1-YFP and pCMK2-YFP phenotypes

**Simple Two-State Promoter Model** This model described the efficient response of pYPS1-YFP to short pulses. The model described a two-state promoter that activates mRNA production which then activates protein production and is depicted in Figure 4A. We modeled these interactions by:

$$\frac{dp_{on}}{dt} = k_{on} \cdot TF \cdot (1 - p_{on}) - k_{off} \cdot p_{on} \tag{1}$$

$$\frac{dmRNA}{dt} = \beta_0 + \beta_1 \cdot p_{on} - \gamma_1 \cdot mRNA \tag{2}$$

$$\frac{dProtein}{dt} = \beta_2 \cdot mRNA - \gamma_2 \cdot Protein \tag{3}$$

In these equations, $p_{on}$ represented promoter activity while mRNA and Protein represented concentration of mRNA and protein, respectively. Here we assumed that promoter activity was conserved such that $1 = p_{on} + p_{off}$. TF represented the concentration of nuclear transcription factor.

The model was characterized by 7 parameters. Most of the activation/inactivation and production/degradation terms were modeled by first-order mass action kinetics. The parameter, $\beta_0$, was zeroth-order, to reflect basal promoter activity. We chose this simple model form because we were interested in a parsimonious model that could explain the experimental phenotype of pYPS1-YFP. Note that the rate of promoter activation was dependent on TF concentration because Crz1 has been shown to activate genes through binding of a known promoter element, the calcineurin-dependent response element (CDRE), through its zinc finger domain (Stathopoulos-Gerontides et al., 1999).

The input to the model was the concentration of nuclear transcription factor (TF), while the output represented protein concentration (Protein). The equations were numerically solved by the ODE solver ode113 for nonstiff differential equations via MATLAB.

Parameters $k_{on}$, $k_{off}$, $\beta_0$,and $\beta_1$ were sampled over 4-5 orders of magnitude systematically and randomly using Latin Hypercube Sampling (LHS). $k_{on}$ and $k_{off}$ varied from 1e-4 to 1. $\beta_0$ was varied from 1e-6 to 1e-2. $\beta_1$ was varied from 1e-4 to 10. Parameters $\beta_2 = 0.06$,$\gamma_1 = 0.05$, and $\gamma_2 = 0.0083$ were fixed to values according to literature (Hansen and O'Shea, 2013; Wang et al., 2002).

**Parameter Search and Model Fitting** From the parameter sets sampled, the slope ratio (defined in Figure 3), a summary metric for the degree of efficiency in response to short pulses, was calculated for each parameter set. These slope ratio values were plotted as heatmaps as a function of the parameters $k_{on}$ and $k_{off}$ to demonstrate the effect of parameters on efficient response to short pulses (slope ratio > 1) or efficient response to continuous pulses (slope ratio < 1).

The model was used to fit the experimental data. Fits of the experimental data to the simple two-promoter state model (Figure 4) were obtained by the following procedure:

1. $10^4$ parameters were randomly sampled using LHS with the aforementioned parameter ranges

2. The model outputs were used to construct the Output-Occupancy plots and compared to the experimental Output-Occupancy plots. Fits were determined to be model outputs that maximize fit to the data points within error of the Output-Occupancy data. Note that the ability of the model to fit the data was the same regardless of the criteria of fit used -- whether the criteria was the model output fit within the error bars of the data or least squared error of the model to the best fit line to the data.

3. Cross-validation of parameter fits to the Output-Occupancy data were then performed using the dose response.The least squared error was used as the metric to assess fit.

From this procedure, we identified parameter sets that fit all of the experimental data for both pYPS1-YFP and pCMK2-YFP.

**Model exploration and sampling details for the pGYP7-YFP phenotype: List of models**

**Two-state models with either** $r_{off}$ **or** $r_{on}$ **thresholding**. These models involved a TF concentration gated activation, $r_{on}$, or inactivation, $r_{off}$, rate constant. The kinetic model with $r_{off}$ thresholding consisted of equations (1)-(3), but utilized the following equation instead of (1):

$$\frac{dp_{on}}{dt} = r_{on} \cdot TF \cdot (1 - p_{on}) - r^*_{off} \cdot p_{on} \qquad (1a)$$

where $r^*_{off}$ = 0 when TF ≥ *threshold*. Otherwise $r^*_{off} = r_{off}$. *threshold* is a parameter value which denoted the threshold TF concentration at which $r^*_{off}$ switches from 0 to $r_{off}$.

The kinetic model with $r_{on}$ thresholding consisted of equations (1)-(3), and utilized the following equation in place of (1):

$$\frac{dp_{on}}{dt} = r^*_{on} \cdot TF \cdot (1 - p_{on}) - r_{off} \cdot p_{on} \qquad (1b)$$

where $r^*_{on} = r_{on}$ when TF ≥ *threshold*. Otherwise $r^*_{on} = 0$.

These two models could represent a binary interaction of the transcription factor (TF) with promoter elements, where below a TF concentration, the TF had no effect on promoter activity and above a TF concentration, the promoter turned on at its maximal rate.

The same parameter ranges were sampled in this model as in the simple kinetic model. The additional parameter *threshold* was sampled randomly from TF = 0 to 2.7, the maximum value of the TF input to the model.

**Cooperative Model**. Similar to the two-state thresholded models, the cooperative model described a nonlinear relationship between TF concentration and protein output. The model was represented by the equations:

$$\frac{dmRNA}{dt} = \beta_0 + \beta_1 \cdot \frac{TF^n}{TF^n + k_d^n} - \gamma_1 \cdot mRNA \quad (4)$$

$$\frac{dProtein}{dt} = \beta_2 \cdot mRNA - \gamma_2 \cdot Protein \quad (5)$$

where n is the hill coefficient, and $k_d = \frac{k_{off}}{k_{on}}$.

The same parameter ranges were sampled in this model as in the simple kinetic model. The additional parameter n was sampled randomly from n = 0.5 to 4, a biologically relevant range (Hansen and O'Shea, 2013).

**3-State Models**. We considered five 3-state models with different relationships of TF and the rate constants for the transition between promoter states. The first such model was a **3-state model**. An additional promoter state, $p_0$, was added. The rate equations describing this model were:

$$\frac{dp_0}{dt} = r_{off} \cdot p_{off} - r_{on} \cdot p_0 \quad (6)$$

$$\frac{dp_{off}}{dt} = k_{off} \cdot p_{on} + r_{on} \cdot p_0 - (r_{off} + k_{on} \cdot TF) \cdot p_{off} \quad (7)$$

$$\frac{dp_{on}}{dt} = k_{on} \cdot TF \cdot p_{off} - k_{off} \cdot p_{on} \quad (8)$$

$$\frac{dmRNA}{dt} = \beta_0 + \beta_1 \cdot p_{on} - \gamma_1 \cdot mRNA \quad (9)$$

$$\frac{dProtein}{dt} = \beta_2 \cdot mRNA - \gamma_2 \cdot Protein \quad (10)$$

In this model, the $p_0$ and $p_{off}$ could be thought of as non-transcribing promoter states that represented nucleosome occluded and open, respectively. $p_{on}$ represented an active transcribing promoter state. The rate constants $r_{off}$ and $r_{on}$ described the transitions between the occluded and open promoter states. The promoter underwent a transition between $p_0$ and $p_{off}$ (with rate constants, $r_{on}$ and $r_{off}$ respectively), but $p_{off}$ was still not a promoter state conducive for transcription. A second transition from $p_{off}$ to $p_{on}$ (with rate constants, $k_{on}$ and $k_{off}$, respectively) was needed to start transcription.

**3-state model with** $r_{off}$ **threshold but no linear TF dependence of** $k_{on}$. This model was the same as the 3-state model with $r_{off}$ threshold, except no linear dependence of the TF in the transition from $p_{off}$ and $p_{on}$. The model was described by equations (6)-(10); the following equations replaced equations (6) and (7):

$$\frac{dp_0}{dt} = r_{off} \cdot p_{off} - r_{on} \cdot TF \cdot p_0 \tag{6d}$$

$$\frac{dp_{off}}{dt} = k_{off} \cdot p_{on} + r_{on} \cdot TF \cdot p_0 - (r_{off} + k_{on} \cdot TF) \cdot p_{off} \tag{7d}$$

**3-state model with** $r_{on}$ **threshold**. This model contained a TF concentration threshold dependence of $r_{on}$ between the occluded $p_0$ and open $p_{off}$ promoter states, and was described by equations (6) - (10); where model equations (6) and (7) were replaced by:

$$\frac{dp_0}{dt} = r_{off} \cdot p_{off} - r^*_{on} \cdot p_0$$

$$\frac{dp_{off}}{dt} = k_{off} \cdot p_{on} + r^*_{on} \cdot p_0 - (r^*_{off} + k_{on} \cdot TF) \cdot p_{off} \quad \text{(6b)}$$

(7b)

where $r^*_{on} = r_{on}$ when TF ≥ *threshold*. Otherwise $r^*_{on} = 0$.

For this model, the transcription factor modulated the rate of transition from $p_0$ to $p_{off}$ such that when TF concentration reached the threshold concentration, *threshold*, the transition rate switches from zero to a value. Biologically, this model could represent TF interaction with chromatin acetylators and other modifiers that could promote an open chromatin conformation on the promoter.

**3-state model with $r_{off}$ threshold**. Similarly, the model with a threshold dependence on the inactivating transition from the open $p_{off}$ to occluded $p_0$ states was described by the model equations (6)-(10); where equations (6) and (7) were replaced by:

$$\frac{dp_0}{dt} = r^*_{off} \cdot p_{off} - r_{on} \cdot p_0 \quad (6c)$$

$$\frac{dp_{off}}{dt} = k_{off} \cdot p_{on} + r_{on} \cdot p_0 - (r^*_{off} + k_{on} \cdot TF) \cdot p_{off} \quad (7c)$$

where $r_{off}^* = 0$ when TF ≥ *threshold*. Otherwise $r_{off}^* = r_{off}$.

For this model, the transcription factor modulated the rate of transition from $p_{off}$ to $p_0$ such that when TF concentration reached the threshold concentration, *threshold*, the transition rate switched from a value to zero. Biologically, this model could represent either physical hindrance of heterochromatin formation or TF-modulated repression of a chromatin de-acetylase. The parameters $r_{on}$ and $r_{off}$ were randomly sampled in the range from 10e-4 to 1.

**3-state model with linear TF dependence of $r_{on}$ and $k_{on.}$** This model contained a linear dependence on TF for the transitions between both $p_0$ to $p_{off}$ and $p_{off}$ to $p_{on}$. This model was described by the model equations (6)-(10); where equations (6) and (7) were replaced by:

$$\frac{dp_0}{dt} = r^*_{off} \cdot p_{off}) - r_{on} \cdot p_0 \quad (6a)$$

$$\frac{dp_{off}}{dt} = k_{off} \cdot p_{on} + r_{on} \cdot p_0 - (r^*_{off} + k_{on}) \cdot p_{off} \quad (7a)$$

**Parameter Search and Model Fitting** Parameter search and model fitting were done in the same way as for modeling of pYPS1-YFP and pCMK2-YFP in the section above, except two rounds of fitting were done with the Output-Occupancy and dose response data of pGYP7-YFP. Fits to the dose response were determined to be parameter sets whose least squared error was 0.8 standard deviations below the mean of the least squared error distribution. An experiment with a strain expressing pTEF1 driven Crz1*-CLASP exposed to short pulsed and continuous inputs, as described in the main text, was used to cross-validate the model fits.

**Detailed exploration of model fits to the pGYP7-YFP data**

The simple kinetic model that described the pYPS1-YFP and pCMK2-YFP phenotypes produced no parameter sets for which a pulsed input generated lower gene expression output than a continuous input (Figure S10A-B). Hence, we explored model elaborations, introduced in the previous section, of the simple promoter switching model.

We first tested whether the **two-state models with either** $r_{off}$ **or** $r_{on}$ **thresholding** (Figure S10E-F, G-H), or the **cooperative model** (Figure S10C-D), could generate a promoter that responded efficiently to continuous pulses. The rationale here was that if the promoter spent some time below its threshold of activation for any input, then the effect of this TF concentration thresholding would be smaller for a continuous pulse that does this once, than for a sequence of short pulses where this would be done repeatedly. In agreement with this intuition, this suite of models was able to generate Output-Occupancy plots that mirrored the pGYP7-YFP experimental data for many parameters (Figure S10 C-D, E-G, G-H, left panel). Figure S10G shows an illustrative example of this class of models, where many parameters sets (380) were shown to maximize fits through the data points within the error bars (Figure S10G, left panel). Upon further fitting with independently-obtained dose response data for pGYP7-YFP (obtained in the same way as explained above for pYPS1-YFP and pCMK2-YFP), this model however failed to fit the data, as did all models with only two promoter states (Figure S10 C-D, E-F, G-H, middle panel). The failure of these models to fit the pGYP7-YFP dose response showed a characteristic pattern -- while the pGYP7-YFP dose response was linear in the TF regime we measured, the computationally predicted dose response was thresholded given the model structure we imposed (Figure S10 C-D, E-F, G-H, middle panel).

Next, we increased the complexity of the model by adding a second layer of promoter transitions to generate the **3-state model** (Figure S10K). This model had the same linear structure as the two-state promoter model of Figure 4, and hence could not produce a more efficient response to continuous input over the pulsed one (Figure S10L).

We also tested whether a **3-state model with** $r_{off}$ **threshold but no linear TF dependence of** $k_{on}$ could produce better fits (Figure S10I). This model was indeed able to generate Output-Occupancy plots that match the pGYP7-YFP experimental data (423 parameter sets within error) (Figure S10J, left panel), but with these parameters, it again produced a thresholded dose response that failed to fit that of pGYP7-YFP (Figure S10J, middle panel).

Following these results, we reasoned that the introduction of a linear dependence on TF concentration in the immediate step before promoter activation could mitigate the effects of a threshold on an earlier promoter transition step, therefore producing a linear dose response. Hence, the **3-state model with $r_{off}$ threshold** was tested.(Figure 5A). With this addition, the model was able to generate Output-Occupancy plots that maximize fit to the experimental data for many parameters (96 parameter sets) and for a subset of those (25 parameter sets), was also able to recapitulate the pGYP7-YFP dose response (Figure 5B-C). The **3-state model with $r_{on}$ threshold** was similarly able to recapitulate the data (Figure S10 O-P), albeit with a poorer fit for the Output-Occupancy plot. Finally, we tested the **3-state model with linear TF dependence of $r_{on}$ and $k_{on}$** (Figure S10M). This model was also able to produce qualitative fits to the Output-Occupancy plot (Figure S10M, left panel) and dose response data (Figure S10M, middle panel).

To further test these three successful models and also further invalidate the discarded models, we subjected them to cross-validation using an independent experiment in a strain where Crz1*-CLASP expression was increased (now expressed from pTEF1 instead of pADH1). We subjected these cells to either short pulses (2 minutes ON/10 minutes OFF) or a continuous input (40 minutes of light) in a timespan of 4 hours, and measured pGYP7-YFP levels at 5 hours. These data revealed that the efficient response to continuous input was still preserved at the higher pTEF1 expression level. All discarded models (Figure S10, right panels) were inconsistent with these data, predicting instead a reversal of the phenotype with an increase in the TF concentration. Notably, the **3-state model with linear TF dependence of $r_{on}$ and $k_{on}$** (Figure S10N, right panel) also failed this cross-validation because the dependence on TF caused the rate of transition from $p_0$ to $p_{off}$ to increase with increased TF, and thus the efficient response to continuous inputs could only be produced for relatively low TF concentrations. Hence, only two minimal models were able to explain all the data we collected (Figure S10 O-P, Figure 5A-D).

### Supplementary Method Details

Delivery of stress inputs for microscopy

For each environmental perturbation, cells were grown overnight to saturation in YPD, diluted in prior to the experiment, and grown to an OD of 0.1. 200ul cells were plated with conA. Just before imaging, the SDC media was removed from the microscopy well and the appropriate environmental stress media was applied to the cells.The media for glucose depletion consisted of 0.67% YNB w/o AA w/ ammonium sulfate, 0.79% CSM, 0.05%

glucose. The media for Osmotic shock was composed of 0.67% YNB w/o AA w/ ammonium sulfate, 0.79% CSM, 2% glucose, and 0.95M sorbitol (Gasch et al., 2000). The phosphate depletion media contained 0.17% Pi-depleted YNB (without amino acids and ammonium sulfate), 0.1% ammonium sulfate, 2% glucose, 25mM sodium citrate (pH 4.7), 0.79% CSM.

RNAseq of Crz1 19A and 5A mutant

Single colonies were picked and grown to saturation in YPD at 30°C overnight. Cells were then diluted and grown for 4 hours to an OD of 0.3. Cells were harvested by centrifugation and frozen with liquid nitrogen. RNA was extracted using phenol chloroform (Sambrook and Russell, 2006). RNA quality was assessed using the Agilent RNA Pico kit. The Lexogen Quantseq 3' mRNA-Seq Library Prep Kit was used for RNA preparation. mRNA libraries were quantified using Qubit dsDNA HS Assay Kit and subject to single-end sequencing on an Illumina HiSeq 4000. Fastq files from illumina were aligned using STAR (Dobin et al., 2013). Downstream processing of read counts and differential gene expression was conducted using custom Matlab scripts.

Automated Flow cytometry

Cells were cultured as described above. Cells were inoculated and grown to an OD of 0.1 then diluted 1:10 for a total of 30mls for each reaction chamber. Cells were subjected to light input perturbations in the reaction chambers. Control of fluidics was achieved using LABView. The first 750 events of sample were discarded, and 2,000-10,000 events were collected per sample. Gene expression was measured using the FITC channel. Cytometer outputs were analyzed using custom matlab scripts. For more information on the automated flow cytometry hardware and LabView control of dynamic sample acquisition, see Harrigan et al., 2018.

### Supplementary Figure Captions

**Supplementary Figure 1: Approximately one-third of TFs are basally cytoplasmic in log phase and a subset are shown to exhibit transient nuclear localization. A)** LOC scores of available transcription factors from the CYCLoPs database are plotted (Chong et al 2015). The LOC score, as defined in Chong et al., 2015, is the number of cells assigned to a specific location (nucleus in this instance) over the total number of cells in any subcellular location. Increasing LOC score denotes increasing nuclear enrichment. **B)** Fold change of nuclear enrichment for a panel of stress-responsive transcription factors (Msn2, Msn4, Stb3, Dot6, and Crz1) are plotted as a function of time in response to environmental inputs

(Glucose depletion and osmotic shock). For glucose depletion, SDC media (2% glucose) is replaced with SD media with 0.05% glucose. For osmotic shock, SDC media is replaced with SDC media with 0.95M sorbitol. Imaging begins at t = 0 after addition of environmental perturbation and samples are imaged every 30 seconds. The solid black lines represent the mean of single cell traces and the shading represents the standard error of the mean.

**Supplementary Figure 2: TF-yeLANS yields different localization depending on TF.** Confocal microscopy images of yeast expressing SynTF-yeLANS and Msn2-yeLANS in the absence of blue light. Red arrows (inset) denote examples of cells that exhibit nuclear/cytoplasmic localization of Msn2-yeLANS.

**Supplementary Figure 3: LANS optimization and LOVTRAP growth comparison. A)** Mean nuclear/cytoplasmic enrichment (nuclear intensity divided by cytoplasm intensity) is plotted as a function of time. Shaded error represents standard deviation and light input regimes are illustrated above graphs. n refers to number of cells tracked and subplot headings (e.g., NLS #3) correspond to NLS peptides listed in Table S2. **B)** Comparison of Mito-LOVTRAP and PM-LOVTRAP strains. Mito-LOVTRAP and PM-LOVTRAP are expressed from pTDH3 (highest), pRPL18B (medium), and pREV1 (lowest) promoters. Strains marked with an asterisk denote those for which growth curves are plotted in Figure 1B of main text. **C)** Comparison of Zdk1-mScarlet-yeLANS + Mito-LOVTrap and CLASP. Both component (e.g. Mito-LOVTRAP and Zdk1-mScarlet-yeLANS) are expressed at the same level, using either pTDH3, pRPL18B, or pREV1 promoters. Background (control strain) denotes the WT strain with pSYNTF-YFP integrated in the LEU locus. Zdk1-mScarlet-yeLANS + Mito-LOVTRAP and mScarlet-CLASP strains also have this integration.

**Supplementary Figure 4: mScarlet-CLASP localization as a function of light input duration. A)** A zoom in of Figure 1D in the main text that shows median duration of nuclear localization as a function of light input duration; the line X = Y is denoted by the dashed line. The zoomed graph illustrates that for short pulse durations, the OFF time -- time that nuclear localization extends past the pulse -- is not linearly related to light input duration. **B)** A scatterplot that shows duration of nuclear localization as a function of light input duration. Each point represents a single cell. **C)** Mean nuclear/cytoplasmic enrichment fold change as a function of time for mScarlet-CLASP induced with blue light. Light input regimes are illustrated above graphs (indicating 0, 2, 4, 8, 10, 20, 40, or 80 minute light input). In all plots, error (bars or shading) represents standard deviation.

**Supplementary Figure 5: Characterization of TF-CLASP strains. A)** Mean FITC/SSC is plotted as a function of light intensity (a.u.) for strains that are exposed to different amplitudes of light for two hours (continuous input). Marked in red is the lowest light dose which yielded near-maximal expression for each strain (>90%); this dose is used in all microscopy and flow cytometry experiments for each strain. Error bars represent standard error of the mean for 9 biological replicates. **B-D)** Each subplot shows the probability density functions of $\log_{10}$(FITC/SSC) of gene expression of corresponding fluorescent promoter fusion for TF-CLASP, TF-NLS (constitutive nuclear localization), and TF-mScarlet (basal localization) strains. Histograms display expression from 9 biological replicates (data from replicates are pooled). TF-mScarlet strains are not exposed to light, for facile comparison to TF-CLASP (No Light) expression. TF-NLS strains are exposed to two hours of blue light (continuous input) to control for the effect of blue light on YFP fluorescence when comparing to TF-CLASP (Light) expression. For all panels, TF cargos are expressed from pRPL18B.

**Supplementary Figure 6: Gal4 can be sequestered and reversibly localized with CLASP; Gal4-CLASP is a functional TF. A)** RFP (top panels) and brightfield (bottom panels) images of mScarlet-tagged Gal4 (left panels) and mScarlet-tagged Gal4-CLASP (right panels). **B)** Gal4 nuclear enrichment is plotted as a function of time. Light input regime is illustrated above graph. Shaded gray area represents 95% confidence interval. **C)** Gene expression of pGal1-YFP resulting from Gal4-CLASP nuclear localization following two hours of blue light input. Shown is the probability density function of $\log_{10}$(FITC/SSC) of pGAL1-YFP in the dark (gray) or after light exposure (blue) .

**Supplementary Figure 7: Characterization of Crz1, Crz1-CLASP, and Crz1*-CLASP nuclear translocation and gene expression with $CaCl_2$ or blue light input. A)** Single cell traces of Crz1 nuclear enrichment over time for 3 representative cells following 0.2M $CaCl_2$. The red lines indicate nuclear localization events. **B)** Schematic of the CRZ1 Open Reading Frame (ORF). Labeled are the Nuclear Localization Sequences (NLS#1 and NLS#2) and the Nuclear Exit Sequence (NES), as well as the Serine-Rich Region (SRR), which is calcium responsive. The light pink triangles denote reported S/T phosphosites, while the dark pink triangles denote reported and characterized S/T phosphosites. The 19 dark and light pink phosphosites are mutated from S/T -> A to construct Crz1*. **C)** Heatmap of gene expression for 5657 genes. Samples in each column of the heatmap are pADH1-Crz1 with no input, pADH1-Crz1-yeLANS with 60 minutes of light, pADH1-Crz1* with no input, and pADH1-Crz1-yeLANS with 0.2M $CaCl_2$ delivered at the start of the experiment. All samples are in log phase and all measurements are taken 60 minutes after delivery of input. **D)** Gene expression (Mean FITC/SSC) of the Crz1 reporter gene pPUN1-YFP driven by either Crz1-CLASP (blue) or Crz1*-CLASP (pink) when given 30 minutes of blue light. **E)** Basal gene expression of pPUN1-YFP for different Crz1 strains: endogenous Crz1, pAdh1-Crz1 in a Crz1 KO background, pAdh1-Crz1* in a Crz1 KO background, and pAdh1-Crz1*-CLASP (without light input) in a Crz1 KO background. **F)** Probability density functions of gene expression of pPUN1-YFP, measured by FITC/SSC, in response to 0.2M $CaCl_2$ (which causes an initial Crz1 nuclear localization pulse of 40-60 minutes), 60 minutes of blue light exposure, and no input. Measurements are taken at 5 hours after delivery of input. **G)** OD600, a measurement for growth, plotted as a function of time, for pAdh1-Crz1 with (blue) and without (red) light input (intensity 512 a.u.) over a period of 24 hours, indicating that light exposure does not affect population growth. Measurements are taken every hour.

**Supplementary Figure 8: A)** Output-Occupancy plot for pYPS1-YFP. **B)** Output-Occupancy plot pCMK2-YFP. **C)** Output-Occupancy plot for pGYP7-YFP. These data are biological replicates taken from different days for the data shown in Figure 3.

**Supplementary Figure 9: Characterization of two-state promoter model of Figure 4 in main text. A)** Slope ratio as a function of parameter values for each sampled parameter. Each blue circle represents a parameter set in the set of 10000 parameter sets searched. $k_{on}$ varies from 0.0001-1, $k_{off}$ from 0.0001-1, $\beta_1$ from 0.0001-10, $\beta_2$ from 0.0001-10 and $\beta_0$ from 0.000001-0.01. $\gamma_1$ is set to 0.05 and $\gamma_2$ to 0.0083. **B)** Output-Occupancy plot for a parameter set with efficient response to short pulses. Parameter values are: $k_{on}$ = 1, $k_{off}$ = 0.8, $\beta_1$ = 0.0001, $\beta_2$ = 0.1, $\gamma_1$ = 0.05, $\gamma_2$ = 0.0083, and $\beta_0$ = 0.000001. $k_{on}$ is reduced 2x, 4x, 8x, 16x, and 32x. The red line represents the gene expression values from increasing pulsed inputs and the blue line represents gene expression values from increasing constant inputs. **C) (left panel)** Heatmap of slope ratios in the $\gamma_1$-$k_{on}$ plane. Parameters are sampled ($\gamma_1$ from 0.0001-10, $k_{on}$ from 0.0001-10) or set ($k_{off}$ = 1, $\beta_1$ = 0.0001, $\beta_2$ =0.1, $\gamma_2$ = 0.0083, $\beta_0$ =0.000001). **(right panel)** Heatmap of slope ratios resulting from the kinetic model in the $\gamma_2$-$k_{on}$ plane. Parameters are sampled ($\gamma_2$ from 0.0001-10, $k_{on}$ from 0.0001-10) or set ($k_{on}$ = 0.001, $\beta_1$ = 0.0001, $\beta_2$ =0.1, $\gamma_1$ = 0.05, $\beta_0$ =0.000001).

**Supplementary Fig 10: Exploration of various models for pGYP7 data A)** Schematic of the kinetic model, where the input is Crz1*-CLASP nuclear localization (TF) and the output is fluorescent protein level (Protein). **B) (left panel)** Output-Occupancy plot for pGYP7-YFP. Circles are experimentally measured values while lines denote the output of the model for 200 parameter sets out of 10000 that maximize fits through data points. The solid line denotes the mean and shaded areas the standard deviation of the model outputs. Parameters are sampled ($k_{on}$ from 0.0001-1, $k_{off}$ from 0.0001-1, $\beta_1$ from 0.0001-10, $\beta_0$ from 0.000001-0.01) or set ($\beta_2$ = 0.06, $\gamma_1$ = 0.05, $\gamma_2$ = 0.0083). **(middle panel)** Dose response plot for pGYP7. The parameters that fit the Output-Occupancy data are used to further fit the dose response of pGYP7-YFP using a best fit to least squared error criterion. Parameter sets below the mean of the least squared error distribution are plotted (solid black line is the mean generated by the model). The gray dots are the experimentally measured dose response. **(right panel)** The parameters that fit the Output-Occupancy are then subject to cross-validation using an experiment where Crz1*-CLASP expression is increased (expressed from a pTEF1 promoter), and cells are exposed to either short (2 minutes ON/10 minutes OFF) or continuous input (40 minutes of light). The model generated outputs (solid red and blue bar) are plotted with the experimental data (hashed red and blue bar). The gray bars are samples not exposed to light. **C)** Schematic of a model with cooperativity **D) (left panel)** Same plots as in (B), with 2481 parameter sets for this model. Parameters are sampled ($k_d$ from 0.01-100, n from 0.5-4, $\beta_1$ from 0.0001-10, $\beta_0$ from 0.000001-0.01) or set ($\beta_2$ = 0.06, $\gamma_1$ = 0.05, $\gamma_2$ = 0.0083). **(middle panel, right panel)** Plotted in the same manner as in (B, middle panel, right panel) with 35 parameter sets. **E)** Schematic of a 2-state model with thresholding on the activation constant, $r_{on}$. **F) (left panel)**. Same plots as in (B), with 148 parameter sets for this model. Parameters are sampled ($r_{on}$ from 0.1-100,$r_{off}$ from 0.1-100, $\beta_1$ from 0.0001-10, $\beta_0$ from 0.000001-0.01) or set ($\beta_2$ = 0.06, $\gamma_1$ = 0.05, $\gamma_2$ = 0.0083, threshold = 0.5). **(middle panel, right panel)** Plotted in the same manner as in (B, middle panel, right panel) with 16 parameter sets. **G)** Schematic of a two-state promoter model with a thresholded promoter inactivation constant, $r_{off}$, where the input is Crz1*-CLASP nuclear localization (TF) and the output is fluorescent protein level (Protein). **H) (left panel)** Same plots as in (B), with 380 parameter sets for this model. Parameters are sampled ($r_{on}$ from 0.0001-1, $r_{off}$ from 0.0001-1, $\beta_1$ from 0.0001-10, $\beta_0$ from 0.000001-0.01, threshold from 0-2.7) or set ($\beta_2$ = 0.06, $\gamma_1$ = 0.05, $\gamma_2$ = 0.0083). **(middle panel, right panel)** Plotted in the same manner as in (B, middle panel, right panel) with 52 parameter sets. **I)** Schematic of a 3-state model with thresholding in the inactivation constant, $r_{off}$, between the promoter off-states, $p_0$ and $p_{off}$, and no TF dependence in the step before promoter activation. **J) (left panel)** Same plots as in (B), with 423 parameter sets for this model. Parameters are sampled ($r_{on}$ from 0.1-100, $r_{off}$ from 0.1-100, $k_{on}$ from 0.0001-1, $k_{off}$ from 0.0001-1, $\beta_1$ from 0.0001-10, $\beta_0$ from 0.000001-0.01, threshold from 0-2.7) or set ($\beta_2$ = 0.06, $\gamma_1$ = 0.05, $\gamma_2$ = 0.0083). **(middle panel, right panel)** Plotted in the same manner as in (B, middle panel, right panel) with 84 parameter sets. **K)** Schematic of a 3-state model with linear dependence of transition from the $p_0$ to $p_{off}$. **L) (left panel)**. Same plots as in (B), with 1288 parameter sets for this model. Parameters are sampled ($r_{on}$ from 0.1-100,$r_{off}$ from 0.1-100, $k_{on}$ from 0.0001-1, $k_{off}$ from 0.0001-1, $\beta_1$ from 0.0001-10, $\beta_0$ from 0.000001-0.01) or set ($\beta_2$ = 0.06, $\gamma_1$ = 0.05, $\gamma_2$ = 0.0083). **(middle panel, right panel)** Plotted in the same manner as in (B, middle panel, right panel) with 16 parameter sets. **M)** Schematic of 3-state model with linear dependence on TF in both transitions from $p_0$ to $p_{off}$ and $p_{off}$ to $p_{on}$. **N) (left panel)** Same plots as in (B), with 1638 parameter sets for this model. Parameters are sampled ($r_{on}$ from 0.1-100, $r_{off}$ from 0.1-100, $k_{on}$ from 0.0001-1, $k_{off}$ from 0.0001-1, $\beta_1$ from 0.0001-10, $\beta_0$ from 0.000001-0.01) or set ($\beta_2$ = 0.06, $\gamma_1$ = 0.05, $\gamma_2$ = 0.0083). **(middle panel, right panel)** Plotted, in the same manner as in (B, middle panel, right panel) with 228 parameter. **O)** Schematic of the 3-state model with thresholding in the activation constant, $r_{on}$, between promoter off-states, $p_0$ and $p_{off}$. **P) (left panel)** Same plots as in (B), with 1649 parameter sets for this model. Parameters are sampled ($r_{on}$ from 0.1-100, $r_{off}$ from 0.1-100, $k_{on}$ from 0.0001-1, $k_{off}$ from 0.0001-1, $\beta_1$ from 0.0001-10, $\beta_0$ from 0.000001-0.01, threshold from 0-0.5) or set ($\beta_2$ = 0.06, $\gamma_1$ = 0.05, $\gamma_2$ = 0.0083). **(middle panel, right panel)** Plotted in the same manner as in (B, middle panel, right panel) with 455 parameter sets. **Q)** Schematic of the 3-state model with thresholding in the inactivation constant, $r_{off}$, between promoter off-states, $p_0$ and $p_{off}$. **R) (left panel)** Same plots as in (B), with 96 parameter sets for this model. Parameters are sampled ($r_{on}$ from 0.1-100, $r_{off}$ from 0.1-100, $k_{on}$ from 0.0001-1, $k_{off}$ from 0.0001-1, $\beta_1$ from 0.0001-10, $\beta_0$ from 0.000001-0.01, threshold from 0-0.5) or set ($\beta_2$ = 0.06, $\gamma_1$ = 0.05, $\gamma_2$ = 0.0083). **(middle panel, right panel)** Plotted in the same manner as in (B, middle panel, right panel) with 25 parameter sets.

**Supplementary Figure 11: Comparison of the 3-state models with either $r_{on}$ or $r_{off}$ thresholding in the transition from $p_0$ to $p_{off}$. A) (upper panel)** Schematic of the 3-state model with thresholding in the activation constant, $r_{on}$, between promoter off-states, $p_0$ and $p_{off}$. **(middle panel)** Heatmap of slope ratio in the $\log_{10}(k_{on}/k_{off})$-$\log_{10}(r_{on}/r_{off})$ plane. $r_{on}$ is set to 0.02 and $k_{on}$ = 0.6. Parameters are sampled ($r_{off}$ from 0.0002-0.02, $k_{off}$ from 0.002-0.2) or set ($\beta_1$ = 0.0001, $\beta_2$ =0.06, $\gamma_1$ =0.05, $\gamma_2$ =0.0083, threshold= 0.5, $\beta_0$ =0.000001). **(lower panel)** Same heatmap as in (A, middle panel) except with $r_{on}$ set to 2, and $r_{off}$ ranges from 0.02-2. **B)**

**(upper panel)** Schematic of the 3-state model with thresholding in the inactivation constant, $r_{off}$, between promoter OFF-states, $p_0$ and $p_{off}$. **(middle panel)** Same heatmap as in (A, middle panel) with $r_{on}$ is set to 0.25 and $k_{on}$ = 0.25. Parameters are sampled ($r_{off}$ from 0.0025-2.5, $k_{off}$ from 0.0025-0.25) or set ($\beta_1$ = 0.0001, $\beta_2$ =0.06, $\gamma_1$ =0.05, $\gamma_2$ =0.0083, threshold= 0.5, $\beta_0$ =0.000001). **(lower panel)** Same heatmap as in (B, middle panel) except with $r_{on}$ set to 2.5, and $r_{off}$ ranges from 0.025-25.

**Supplementary Figure 12: Additional parameter requirements of the 3-state $r_{on}$ threshold model for fitting pGYP7-YFP. A)** Heatmap in the $\log_{10}(k_{on}/k_{off})$-$\log_{10}(r_{on}/r_{off})$ plane of slope ratio of Output-Occupancy relationship, previously described in Figure 5E. **B-C) (upper panels)** Output-Occupancy plots are generated by the model for different parameter sets that correspond to points 3 and 4 in the heatmap in A. The slope ratio for point 3 is 1.05 with $\log_{10}(k_{on}/k_{off})$ = -1.58 and $\log_{10}(r_{on}/r_{off})$ = 0.6. The slope ratio for point 4 is 1.25 with $\log_{10}(k_{on}/k_{off})$ = 0.1 and $\log_{10}(r_{on}/r_{off})$ = -0.89. Point 3 is chosen to highlight the effect of decreasing $r_{off}$, while Point 4 is chosen to highlight the effect of decreasing $k_{off}$. **(middle panels)** Example of a time course of promoter state $p_0$ for a light input that produces the equivalent of 40 minutes (dotted line in upper panel) in nuclear localization either continuously or in short pulses. Solid lines are the $p_0$ pulses while shading denotes nuclear localization. The black double arrows denote the maximum depletion of the $p_0$ state for the pulsed input. **(lower panels)** Example of a time course of promoter activity $p_{on}$ for a light input that produces the equivalent of 40 minutes (dotted line in upper panel) in nuclear localization either continuously or in short pulses, similar to the (middle panels). The red and blue dashes represent residual promoter activity beyond the nuclear localization input. The red residual promoter activity is repeated 15 times while the blue residual activity is repeated one time. The ▲ bar denotes the difference between the amplitudes generated by the 2 minute pulse and 40 minute continuous input.

**Supplementary Figure 13: Correlation of nucleosome occupancy and sensitivity to pulsing. A)** Heatmap of H3 occupancy for the Crz1 target genes as specified by Yoshimoto 2002. H3 occupancy is defined as counts of H3 enrichment over the IgG antibody, which signals no pull down of histones. The dataset and determination of start sites are obtained from Sen et al., 2015 and Malabat et al., 2015, respectively. The software deepTools 2.0 is used to compute the H3 occupancy values. -1 and +1 kb from the transcription start site (TSS) is used. The positions of YPS1, CMK2, and GYP7 in the heatmap are denoted with black triangles. **B)** Slope ratios of Crz1 target genes as a function of their mean H3 nucleosome occupancy scores averaged from -1kb to the Transcription Start Site (TSS). The correlation coefficient is $r^2 = 0.26$.

### Supplementary Datasets

**Supplementary Table 1**. The NLS optimization document contains the amino acid and peptide sequences used in the screen for NLSs with maximal dynamic range.

**Supplementary Table 2**. Strain list.

**Supplementary Table 3**. Plasmid list.

**Supplementary Table 4**. Transfer function of OptoPlate light input to intensity.

### Supplementary References


1. Chi, Y., Huddleston, M.J., Zhang, X., Young, R.A., Annan, R.S., Carr, S.A., and Deshaies, R.J. (2001). Negative regulation of Gcn4 and Msn2 transcription factors by Srb10 cyclin-dependent kinase. Genes Dev. *15*, 1078–1092.
2. Durchschlag, E., Reiter, W., Ammerer, G., and Schüller, C. (2004). Nuclear localization destabilizes the stress-regulated transcription factor Msn2. J. Biol. Chem. *279*, 55425–55432.
3. Hansen, A.S., and O'Shea, E.K. (2013). Promoter decoding of transcription factor dynamics involves a trade-off between noise and control of gene expression. Mol. Syst. Biol. *9*, 704.
4. Stathopoulos-Gerontides, A., Guo, J.J., and Cyert, M.S. (1999). Yeast calcineurin regulates nuclear localization of the Crz1p transcription factor through dephosphorylation. Genes Dev. *13*, 798–803.
5. Wang, Y., Liu, C.L., Storey, J.D., Tibshirani, R.J., Herschlag, D., and Brown, P.O. (2002). Precision and functional specificity in mRNA decay. Proc. Natl. Acad. Sci. U. S. A. *99*, 5860–5865.

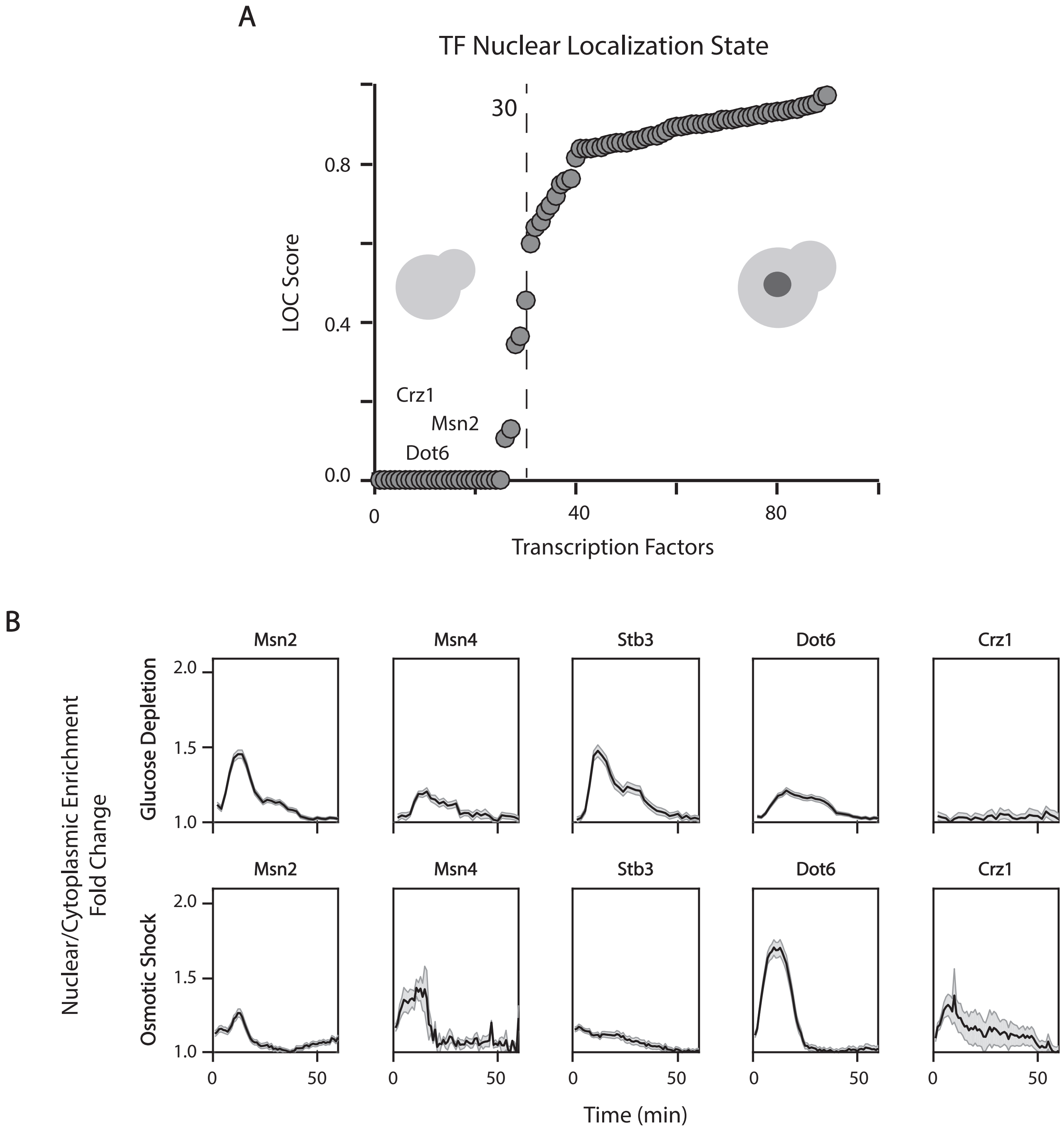


Supplementary Figure 1

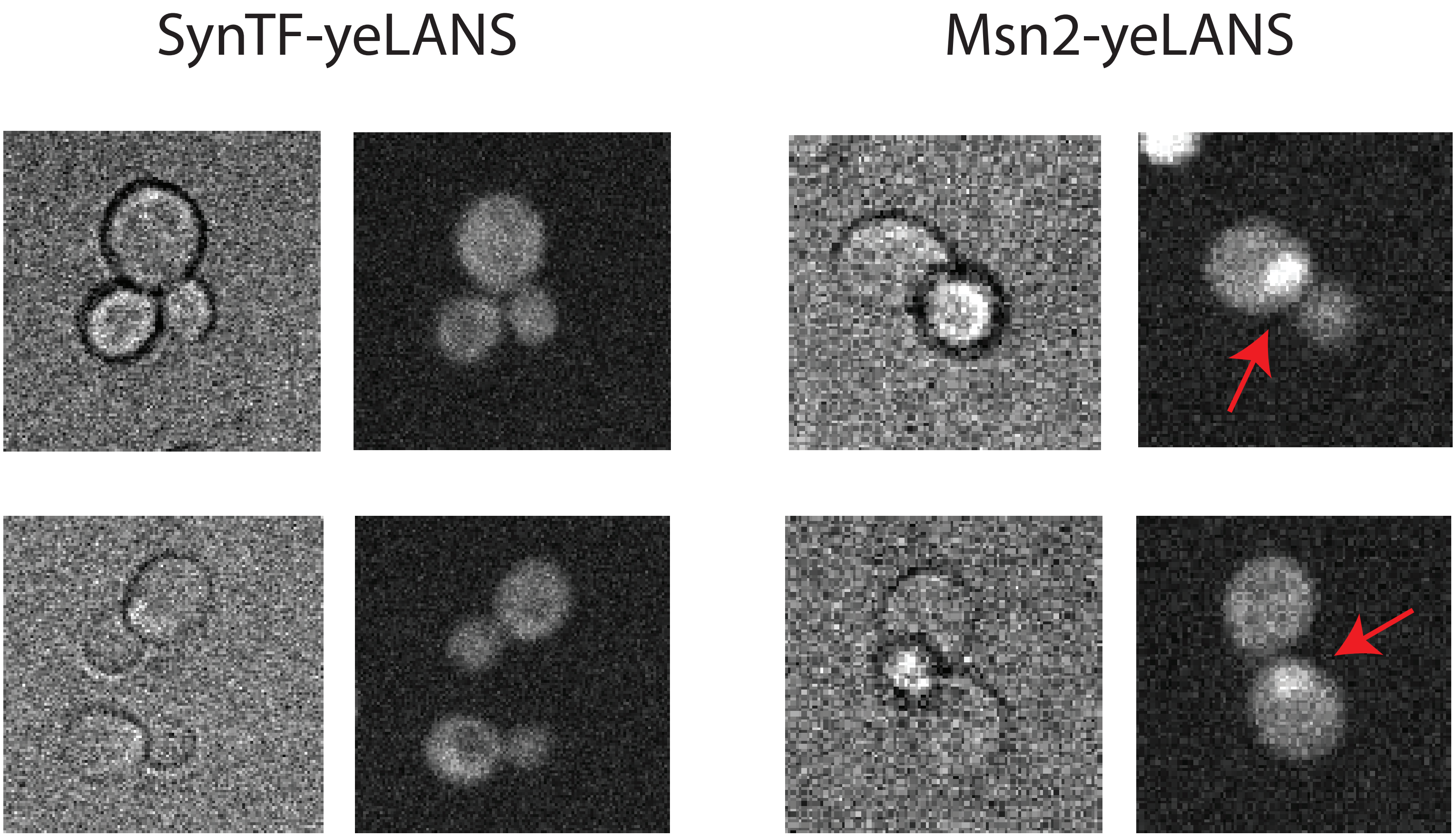


Supplementary Figure 2

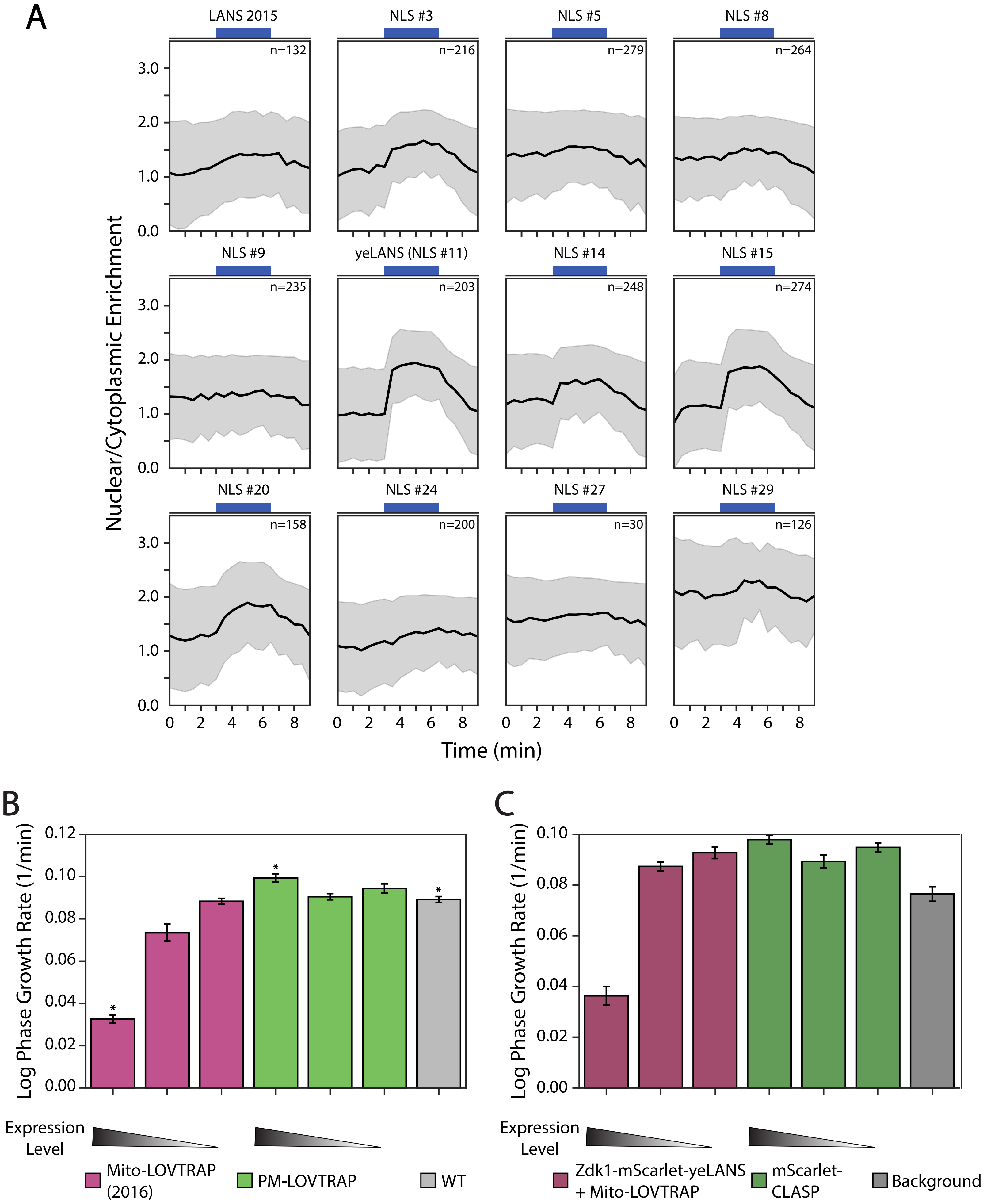


Supplementary Figure 3

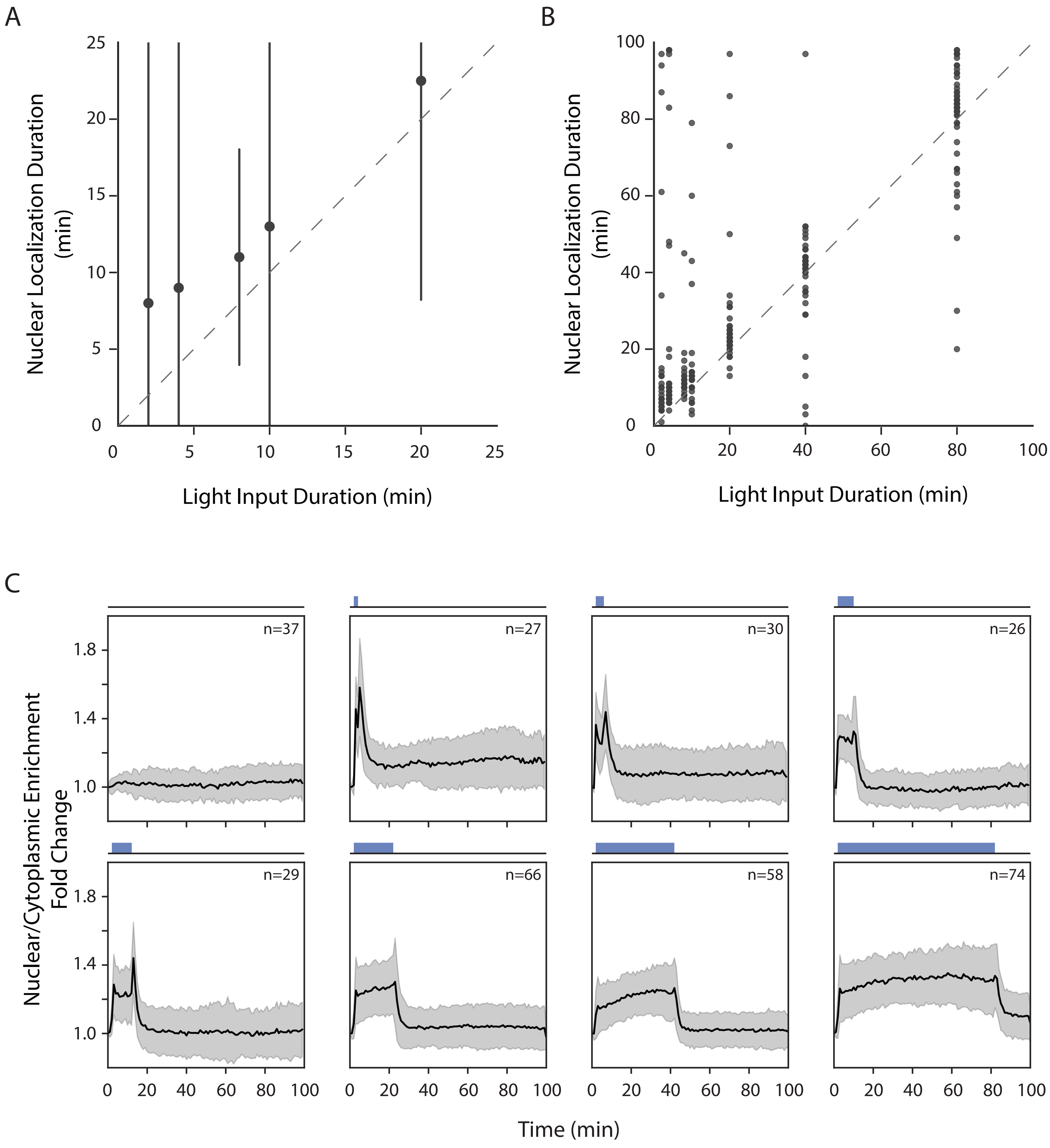


Supplementary Figure 4

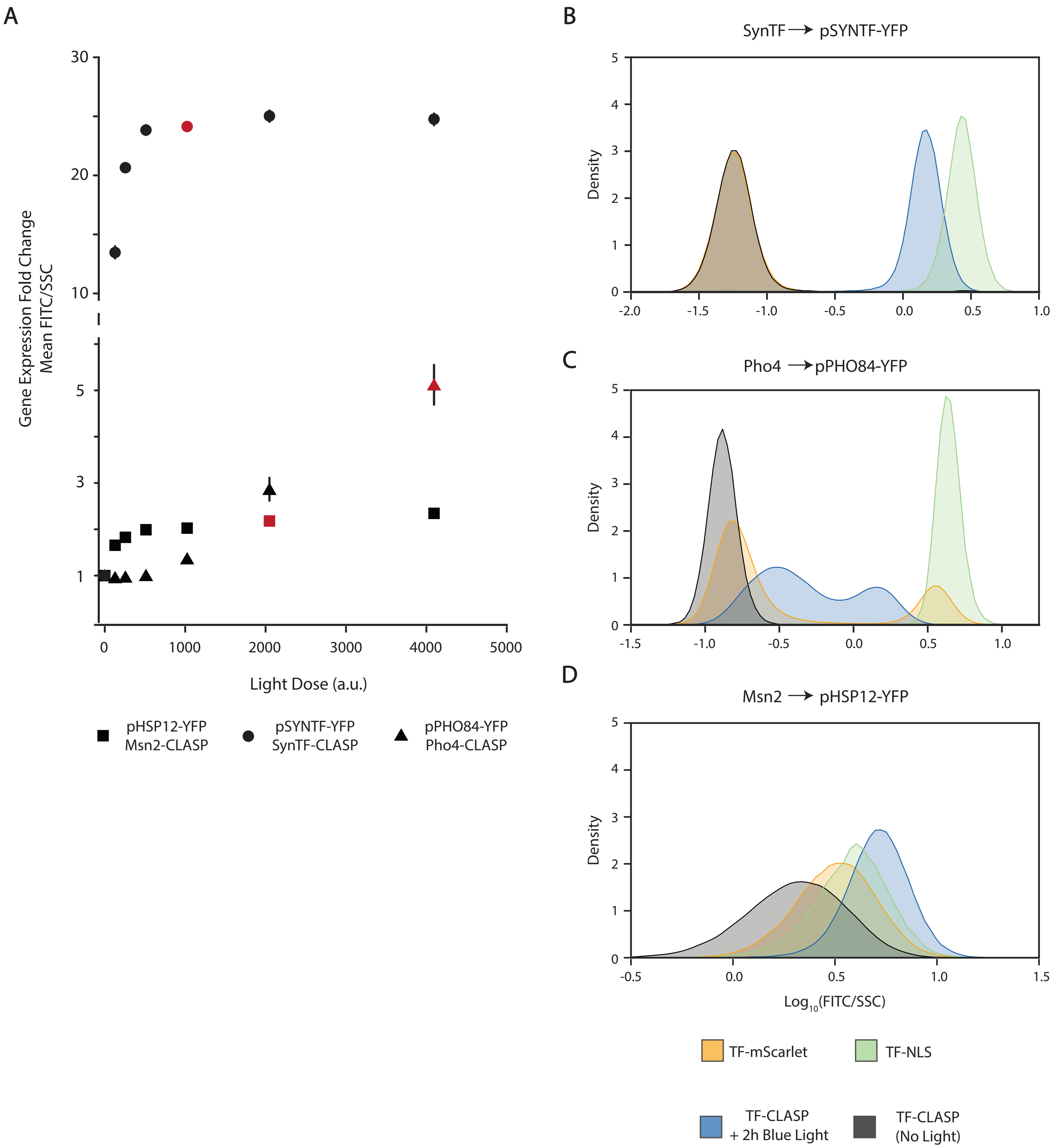


Supplementary Figure 5

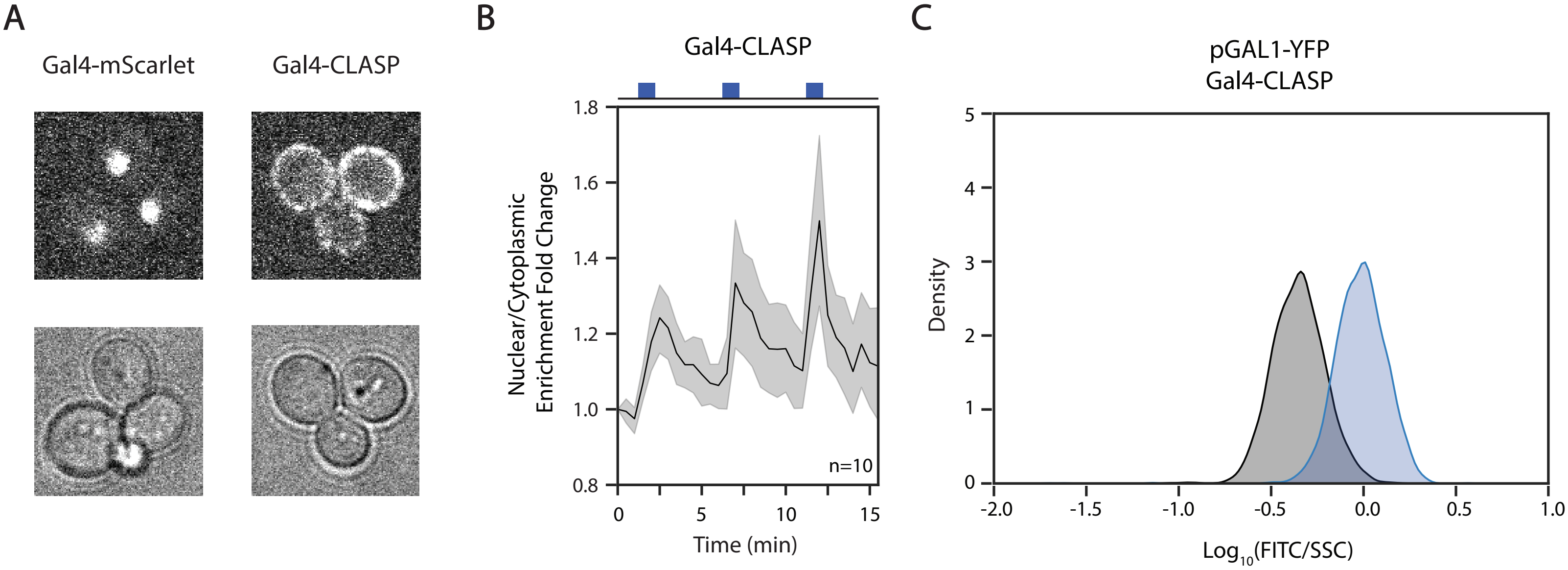


Supplementary Figure 6

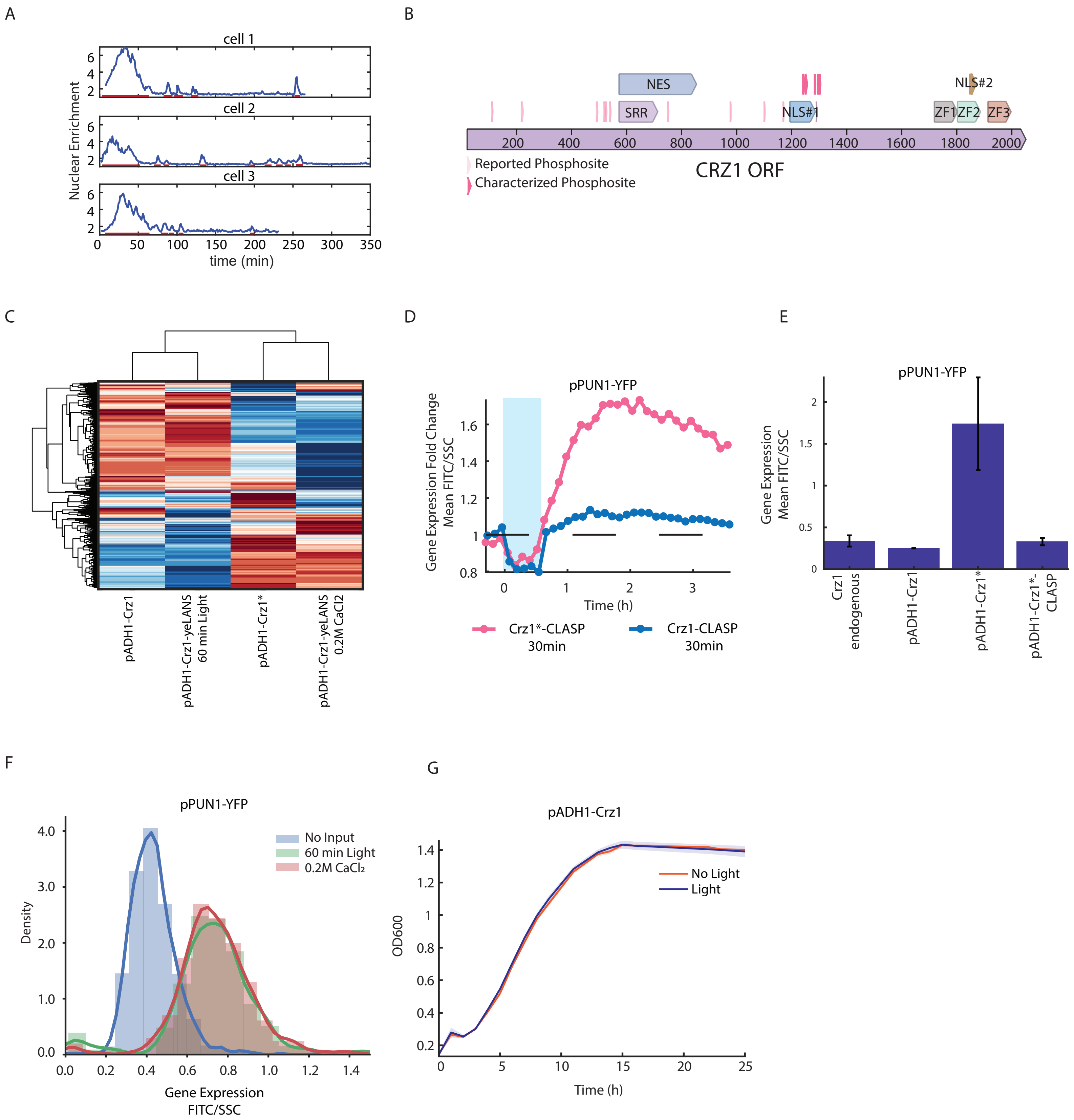


Supplementary Figure 7

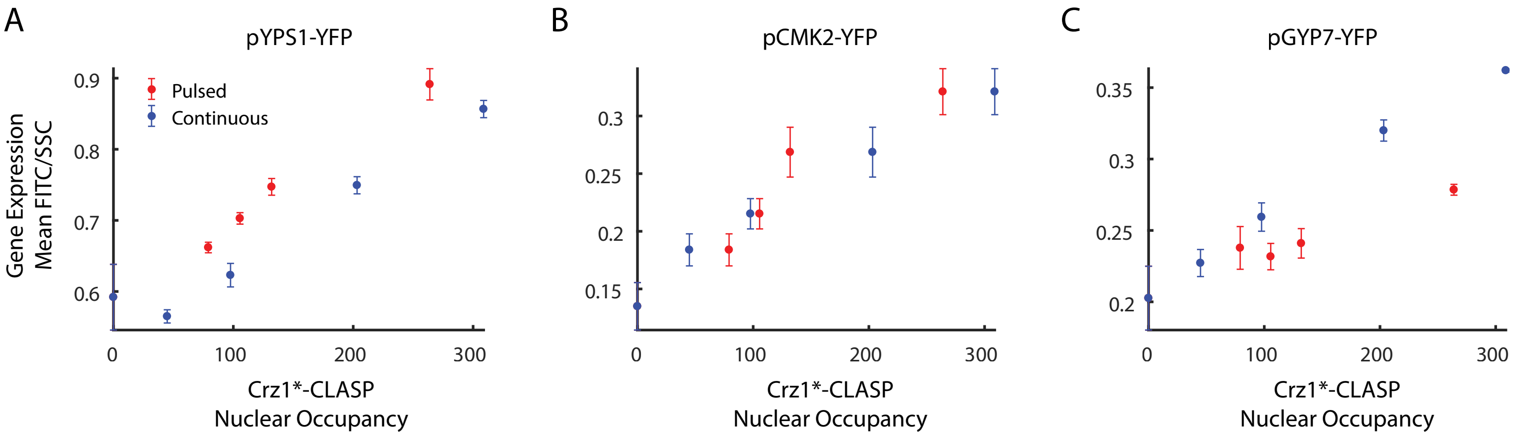


Supplementary Figure 8

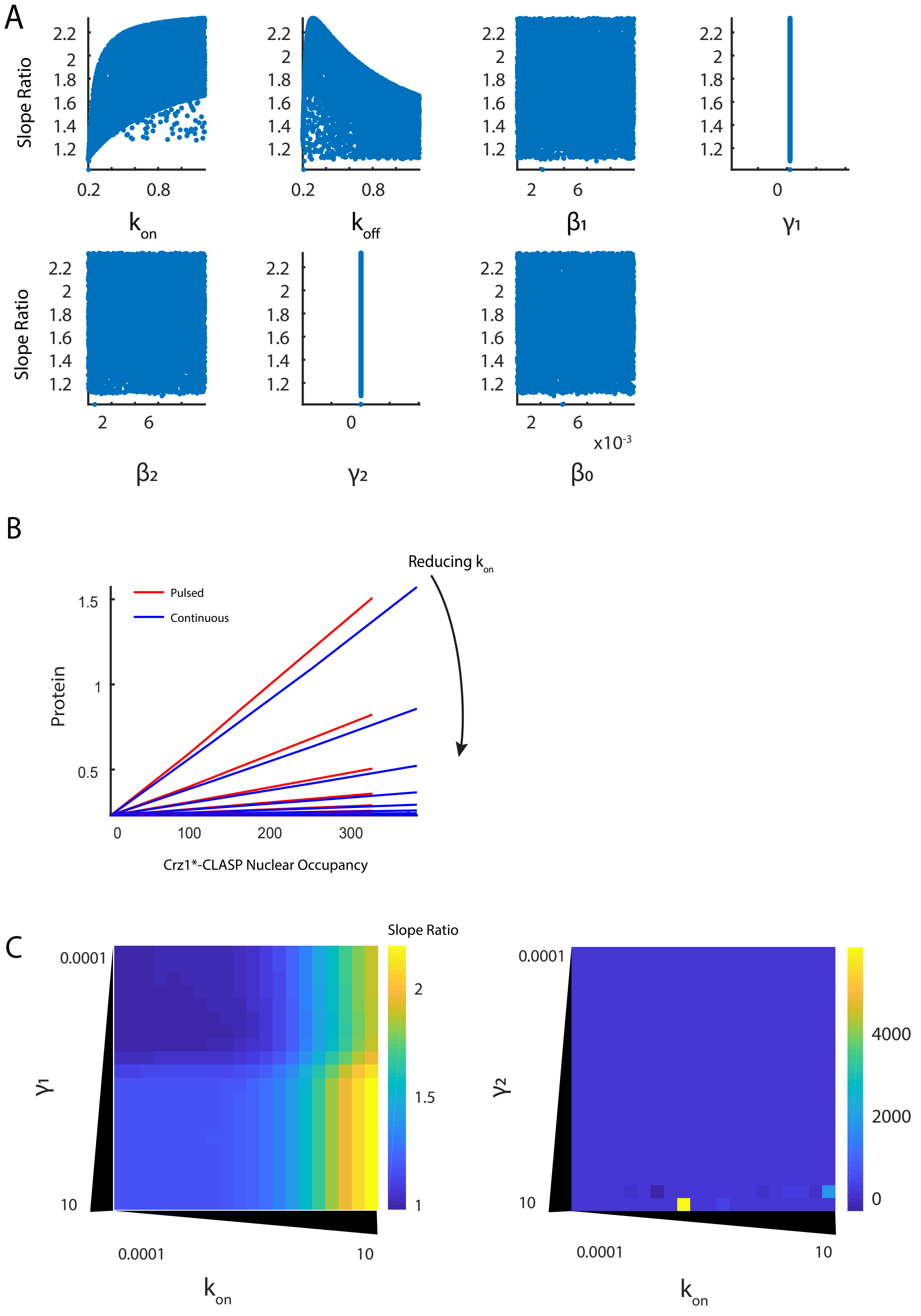


Supplementary Figure 9

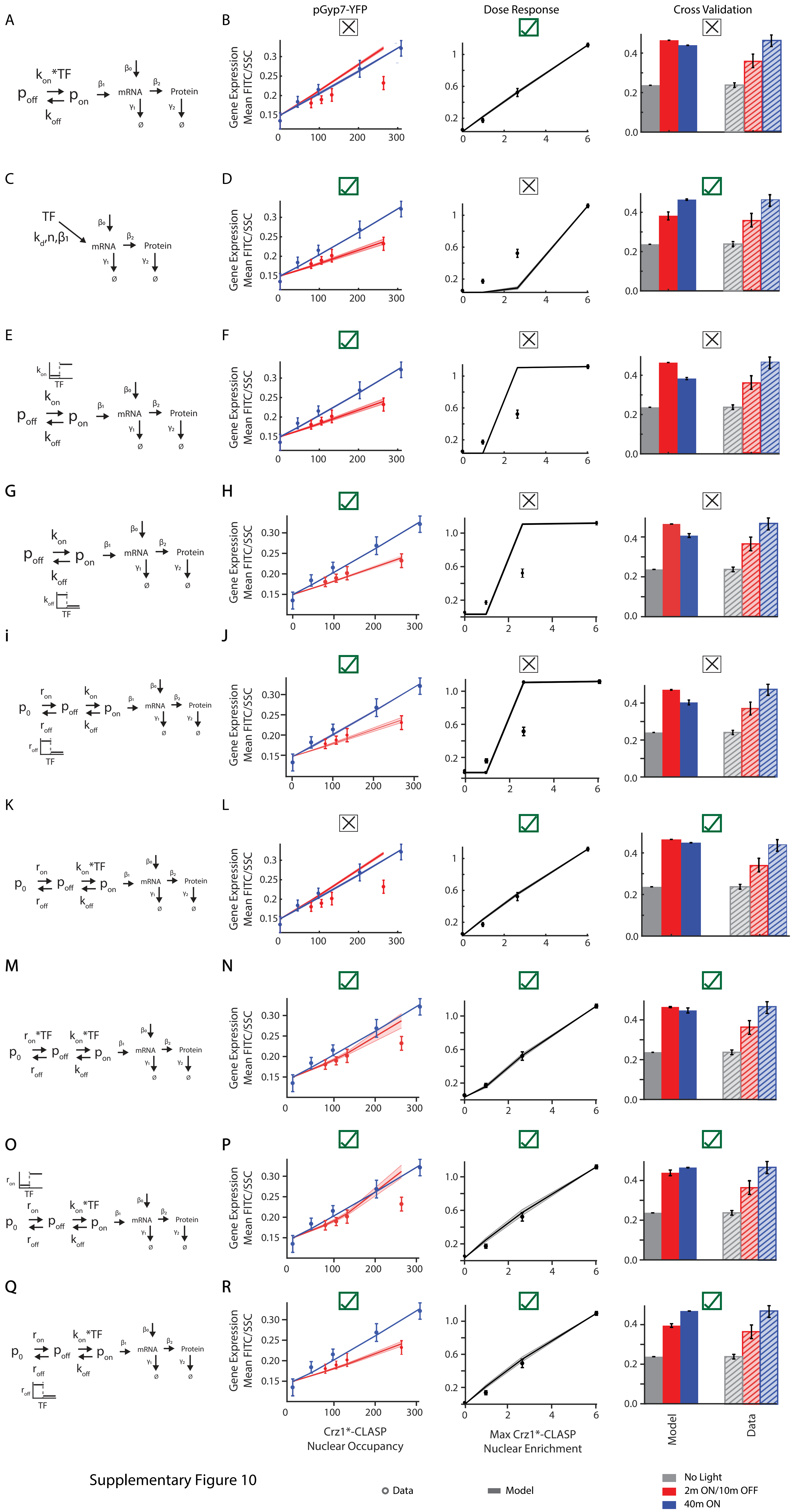


Supplementary Figure 10

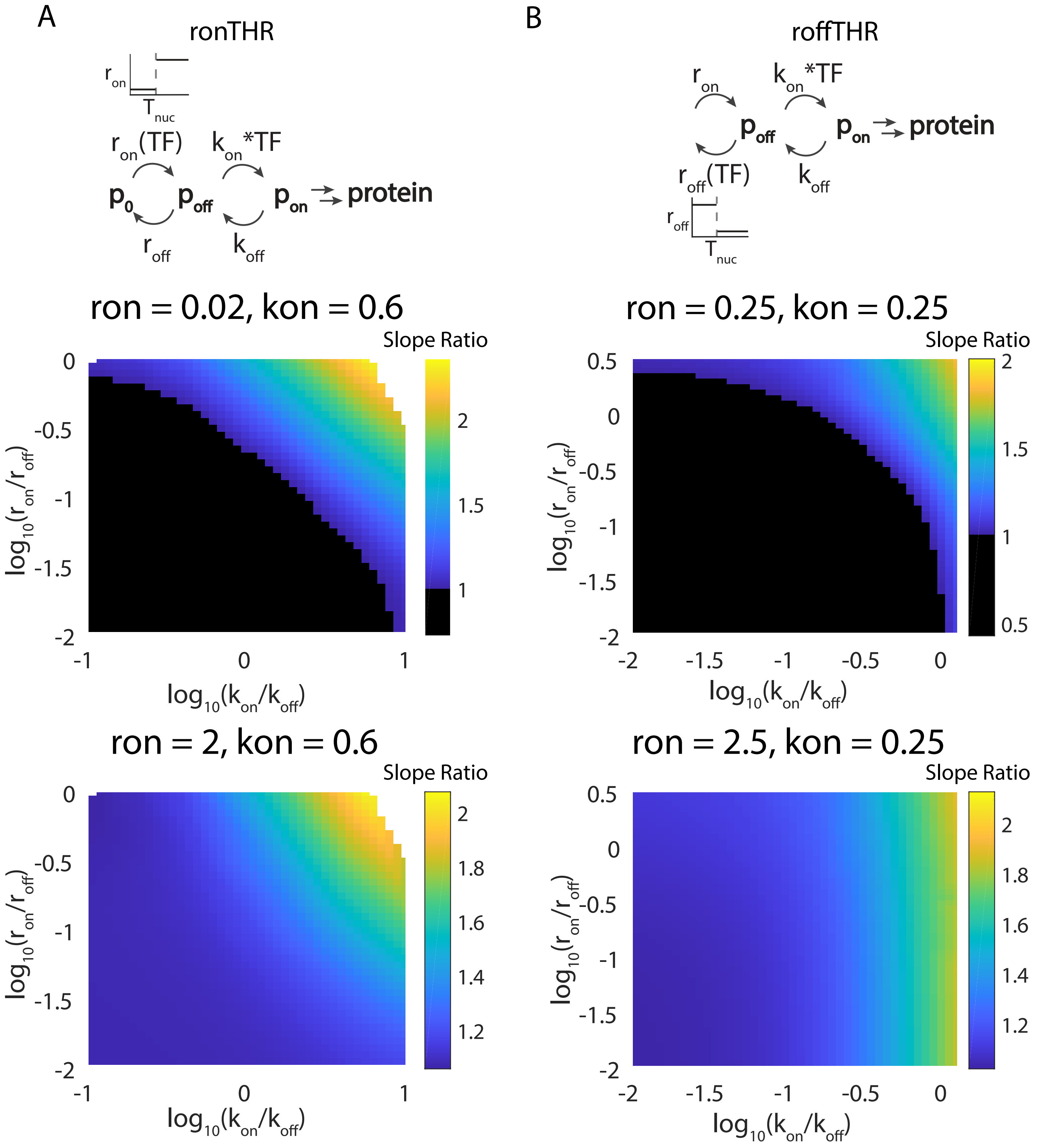


Supplementary Figure 11

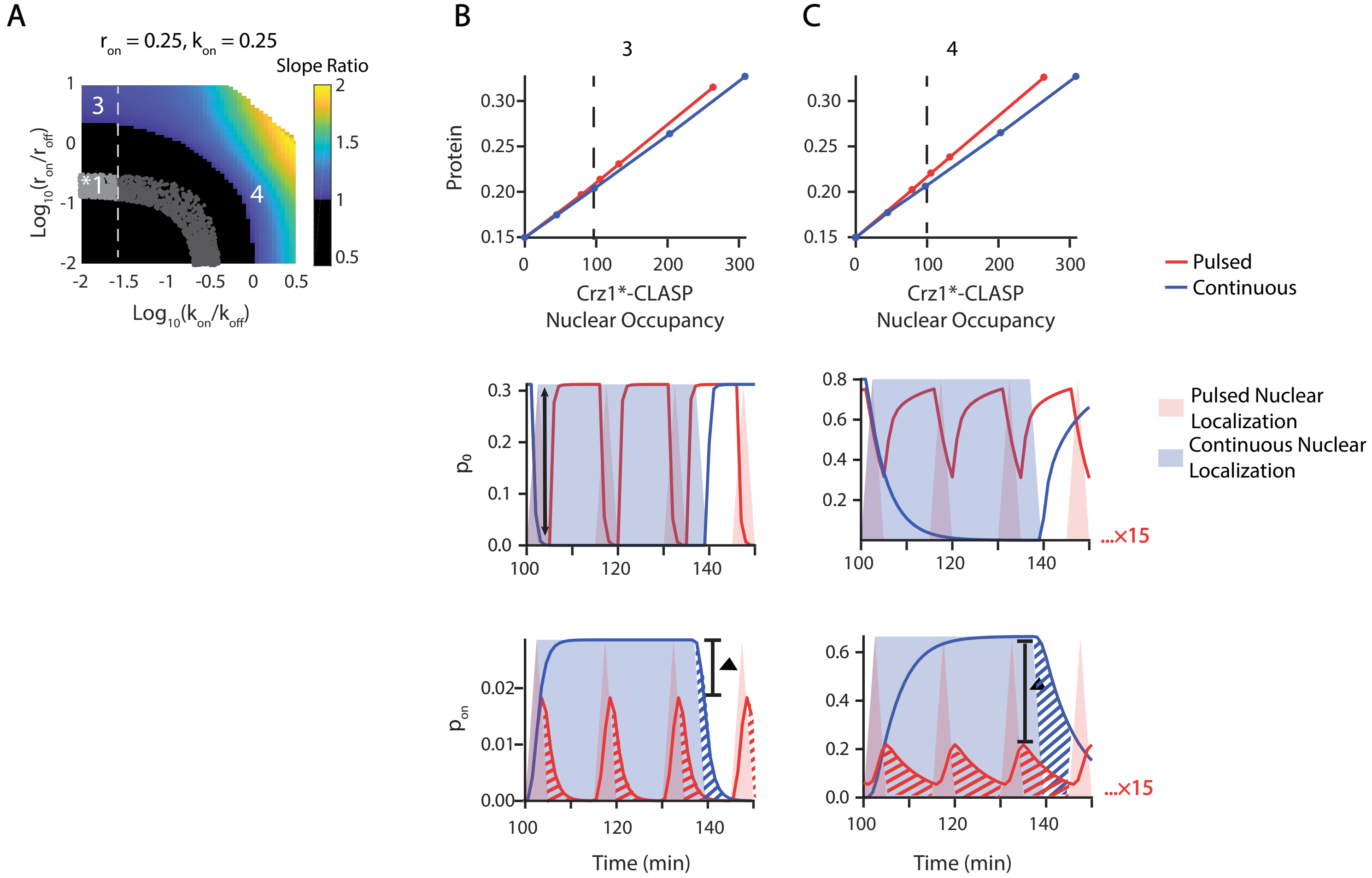


Supplementary Figure 12

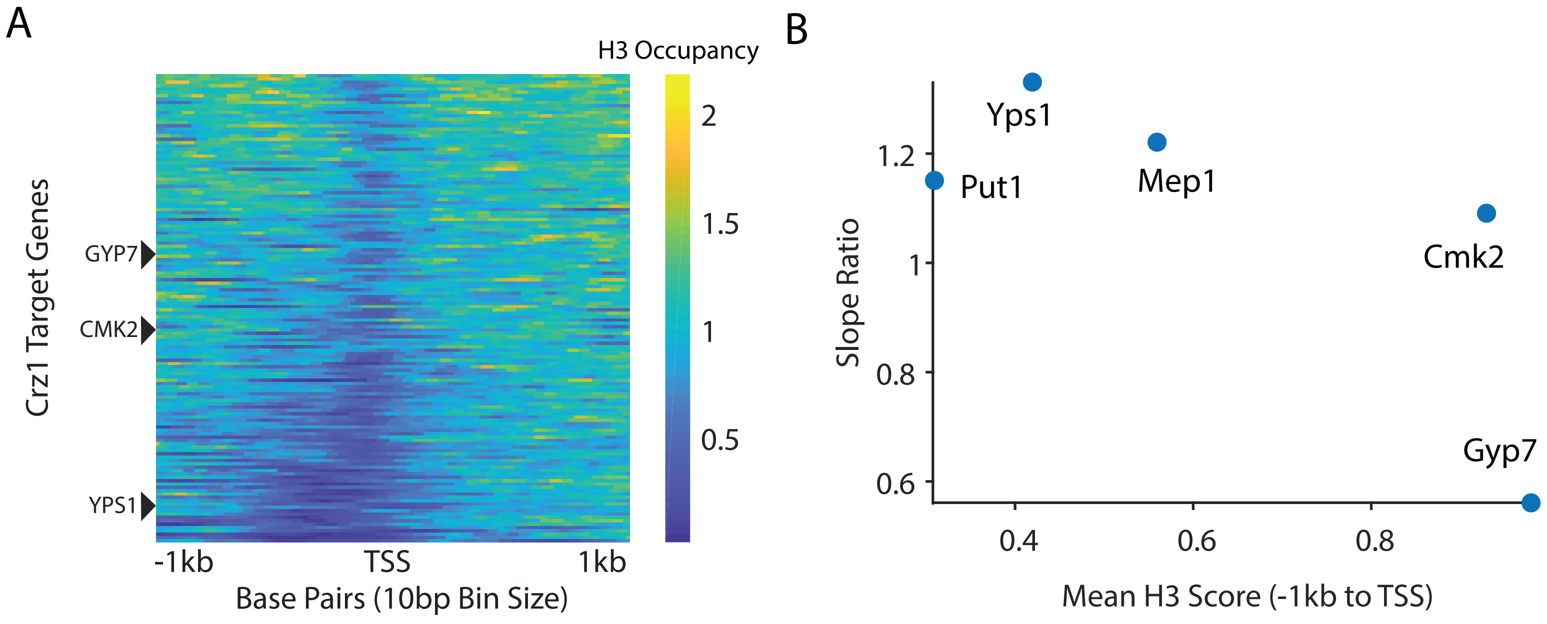


Supplementary Figure 13

| # | NLS | score (Kosugi 2009) | aa length | class |
|---|---|---|---|---|
| 0 | paaKRvKld | na | 9 | original LANS, class2 |
| 1 | IKKTAENIDMAAKRVKLD | na | 18 | original LANS variant, class2 |
| 2 | IKKTAENIDEAAKELPDANL | na | 20 | original LANS variant, class2 |
| 3 | raaKRpRtt | 10 | 9 | class2 |
| 4 | raaKRlRtt | 9 | 9 | class2 |
| 5 | paaKRpRtt | 9 | 9 | class2 |
| 8 | apaKRaRtt | 8 | 9 | class2 |
| 9 | paaKRlCtt | 9 | 9 | class2 |
| 11 | aaaKRswsmaf | 10 | 11 | class3 |
| 14 | aaaKRswvmaf | 9 | 11 | class3 |
| 15 | aaaKRswsaaf | 10 | 11 | class3 |
| 20 | KRpatlandspaaKRR | 9 | 16 | bipartite |
| 21 | KRpaeldgadnasqaaKRR | 10 | 19 | bipartite |
| 22 | KRpaeldeadnasqaaKRR | 10 | 19 | bipartite |
| 24 | KRKRwendip | | 10 | class1 |
| 27 | psRKRKRdhyav | | 12 | class1 |
| 29 | tspsRKRKwdqv | | 12 | class1 |

Supplementary Table 1

| Name | Base Strain | Genome | Description | Figure Correspondence |
|---|---|---|---|---|
| yWCD230 | BY4741 | BY4741, HIS3 repaired | Wild type strain for plasmid integration | Supp Figure 3B |
| yLO213 | BY4741 | ura3: ConS-pRPL18B-Zdk1-mScarlet-yeLANS-tADH1-Con1-pRPL18B-Hs_RGS2(33-67)_13aa-iRFP713 (iRFP)-asLOV2(404-546)-tPGK1-ConE-URA3 (pSc-native-tSc)<br>leu2: p43_8(8x)-Venus-tPGK1-LEU2 (pSc-native-tSc) | mScarlet-LANSTrap strain used for microscopy studies | Figure 1C-E; Supp Figure 3C, 4 |
| yLO133 | BY4741 | ura3: ConS-pRPL18B-Zdk1-VP16-ZF43_8-mScarlet-yeLANS-tADH1-Con1-pRPL18B-Hs_RGS2(33-67)_13aa-iRFP713 (iRFP)-asLOV2(404-546)-tPGK1-ConE-URA3 (pSc-native-tSc)<br>leu2: p43_8(8x)-Venus-tPGK1-LEU2 (pSc-native-tSc) | SynTF-LANSTrap | Figure 2; Supp Figure 5A-B |
| yLO204 | BY4741 | leu2: ConLS'-pPHO84-Venus-tPGK1-ConRE'-LEU2 (pSc-native-tSc)<br>ura3: ConS-pRPL18B-Zdk1-Pho4-mScarlet-yeLANS-tADH1-Con1-pTDH3-Hs_RGS2(33-67)_13aa-iRFP713 (iRFP)-asLOV2(404-546)-tPGK1-ConE-URA3 (pSc-native-tSc)<br>PHO4::HIS3 | Pho4-LANSTrap | Figure 2; Supp Figure 5A,C |
| yLO228 | BY4741 | ura3: ConS-pRPL18B-Zdk1-Msn2-mScarlet-yeLANS-tADH1-Con1-pTDH3-Hs_RGS2(33-67)_13aa-iRFP713 (iRFP)-asLOV2(404-546)-tPGK1-ConE'-URA3 (pSc-native-tSc)<br>MSN4::URA3 (broken with 5'FOA)<br>MSN2::HIS3<br>leu2: ConS-pHsp12-Venus-tPGK1-ConE'-LEU2 (pSc-native-tSc) | Msn2-LANSTrap | Figure 2; Supp Figure 5A,D |
| yLO240 | BY4741 | ura3: ConLS'-pTDH3-TOM20_TMD(1-39)-iRFP713 (iRFP)-asLOV2(404-546)-tPGK1-ConRE'-URA3 (pSc-native-tSc) | Mito-LOVTRAP (high expression level) | Figure 1B; Supp Figure 3B |
| yLO241 | BY4741 | ura3: ConLS'-pRPL18B-TOM20_TMD(1-39)-iRFP713 (iRFP)-asLOV2(404-546)-tPGK1-ConRE'-URA3 (pSc-native-tSc) | Mito-LOVTRAP (medium expression level) | Supp Figure 3B |
| yLO242 | BY4741 | ura3: ConLS'-pREV1-TOM20_TMD(1-39)-iRFP713 (iRFP)-asLOV2(404-546)-tPGK1-ConRE'-URA3 (pSc-native-tSc) | Mito-LOVTRAP (low expression level) | Supp Figure 3B |
| yLO243 | BY4741 | ura3: ConLS'-pTDH3-Hs_RGS2(33-67)_13aa-iRFP713 (iRFP)-asLOV2(404-546)-tPGK1-ConRE'-URA3 (pSc-native-tSc) | PM-LOVTRAP (high expression level) | Figure 1B; Supp Figure 3B |
| yLO244 | BY4741 | ura3: ConLS'-pRPL18B-Hs_RGS2(33-67)_13aa-iRFP713 (iRFP)-asLOV2(404-546)-tPGK1-ConRE'-URA3 (pSc-native-tSc) | PM-LOVTRAP (medium expression level) | Supp Figure 3B |
| yLO245 | BY4741 | ura3: ConLS'-pREV1-Hs_RGS2(33-67)_13aa-iRFP713 (iRFP)-asLOV2(404-546)-tPGK1-ConRE'-URA3 (pSc-native-tSc) | PM-LOVTRAP (low expression level) | Supp Figure 3B |
| yLO210 | BY4741 | ura3: ConS-pTDH3-Zdk1-mScarlet-yeLANS-tADH1-Con1-pTDH3-Hs_RGS2(33-67)_13aa-iRFP713 (iRFP)-asLOV2(404-546)-tPGK1-ConE-URA3 (pSc-native-tSc)<br>leu2: p43_8(8x)-Venus-tPGK1-LEU2 (pSc-native-tSc) | mScarlet-LANSTrap (high expression level) | Supp Figure 3C |
| yLO215 | BY4741 | ura3: ConS-pREV1-Zdk1-mScarlet-yeLANS-tADH1-Con1-pREV1-Hs_RGS2(33-67)_13aa-iRFP713 (iRFP)-asLOV2(404-546)-tPGK1-ConE-URA3 (pSc-native-tSc)<br>leu2: p43_8(8x)-Venus-tPGK1-LEU2 (pSc-native-tSc) | mScarlet-LANSTrap (low expression level) | Supp Figure 3C |
| yLO216 | BY4741 | ura3: ConS-pTDH3-Zdk1-mScarlet-yeLANS-tADH1-Con1-pTDH3-TOM20_TMD(1-39)-iRFP713 (iRFP)-asLOV2 (404-546)-tPGK1-ConE-URA3 (pSc-native-tSc)<br>leu2: p43_8(8x)-Venus-tPGK1-LEU2 (pSc-native-tSc) | Zdk1-mScarlet-yeLANS + mito-LOVTRAP (high expression level) | Supp Figure 3C |
| yLO219 | BY4741 | ura3: ConS-pRPL18B-Zdk1-mScarlet-yeLANS-tADH1-Con1-pRPL18B-TOM20_TMD(1-39)-iRFP713 (iRFP)-asLOV2(404-546)-tPGK1-ConE-URA3 (pSc-native-tSc)<br>leu2: p43_8(8x)-Venus-tPGK1-LEU2 (pSc-native-tSc) | Zdk1-mScarlet-yeLANS + mito-LOVTRAP (medium expression level) | Supp Figure 3C |
| yLO221 | BY4741 | ura3: ConS-pREV1-Zdk1-mScarlet-yeLANS-tADH1-Con1-pREV1-TOM20_TMD(1-39)-iRFP713 (iRFP)-asLOV2 (404-546)-tPGK1-ConE-URA3 (pSc-native-tSc)<br>leu2: p43_8(8x)-Venus-tPGK1-LEU2 (pSc-native-tSc) | Zdk1-mScarlet-yeLANS + mito-LOVTRAP (low expression level) | Supp Figure 3C |
| yLO152 | BY4741 | ura3: ConS-pRPL18B-VP16-ZF43_8-mScarlet-yeLANS-tADH1-Con1-pRPL18B-Hs_RGS2(33-67)_13aa-iRFP713 (iRFP)-asLOV2(404-546)-tPGK1-ConE-URA3 (pSc-native-tSc)<br>leu2: p43_8(8x)-Venus-tPGK1-LEU2 (pSc-native-tSc) | SynTF-yeLANS | Supp Figure 2 |
| yLO179 | BY4741 | ura3: ConS-pRPL18B-Msn2-mScarlet-yeLANS-tADH1-Con1-pTDH3-Hs_RGS2(33-67)_13aa-iRFP713 (iRFP)-asLOV2(404-546)-tPGK1-ConE'-URA3 (pSc-native-tSc)<br>MSN2::HIS3<br>leu2: ConS-pHsp12-Venus-tPGK1-ConE'-LEU2 (pSc-native-tSc) | Msn2-yeLANS | Supp Figure 2 |
| yLO171 | BY4741 | ura3: ConS-pRPL18B-VP16-ZF43_8-mScarlet-NLS11-tADH1-Con1-pRPL18B-Hs_RGS2(33-67)_13aa-iRFP713 (iRFP)-asLOV2(404-546)-tPGK1-ConE-URA3 (pSc-native-tSc)<br>leu2: p43_8(8x)-Venus-tPGK1-LEU2 (pSc-native-tSc) | SynTF-NLS11 for comparison to SynTF-LANSTrap | Supp Figure 5B |
| yLO236 | BY4741 | ura3: ConS-pRPL18B-VP16-ZF43_8-mScarlet-tADH1-Con1-pRPL18B-Hs_RGS2(33-67)_13aa-iRFP713 (iRFP)-asLOV2(404-546)-tPGK1-ConE-URA3 (pSc-native-tSc)<br>leu2: p43_8(8x)-Venus-tPGK1-LEU2 (pSc-native-tSc) | SynTF-mScarlet for comparison to SynTF-LANSTrap | Supp Figure 5B |

Supplementary Table 2

| Name | Base Strain | Genome | Description | Figure Correspondence |
|---|---|---|---|---|
| yLO224 | BY4741 | ura3: ConS-pRPL18B-Pho4-mScarlet-tADH1-Con1-pTDH3-Hs_RGS2(33-67)_13aa-iRFP713 (iRFP)-asLOV2(404-546)-tPGK1-ConE-URA3 (pSc-native-tSc)<br>leu2: ConLS'-pPHO84-Venus-tPGK1-ConRE'-LEU2 (pSc-native-tSc)<br>PHO4::HIS3 | Pho4-mScarlet for comparison to Pho4-LANSTrap | Supp Figure 5C |
| yLO225 | BY4741 | ura3: ConS-pRPL18B-Pho4-mScarlet-NLS11-tADH1-Con1-pTDH3-Hs_RGS2(33-67)_13aa-iRFP713 (iRFP)-asLOV2(404-546)-tPGK1-ConE-URA3 (pSc-native-tSc)<br>leu2: ConLS'-pPHO84-Venus-tPGK1-ConRE'-LEU2 (pSc-native-tSc)<br>PHO4::HIS3 | Pho4-NLS11 for comparison to Pho4-LANSTrap | Supp Figure 5C |
| yLO229 | BY4741 | ura3: ConS-pRPL18B-Msn2-mScarlet-NLS11-tADH1-Con1-pTDH3-Hs_RGS2(33-67)_13aa-iRFP713 (iRFP)-asLOV2(404-546)-tPGK1-ConE'-URA3 (pSc-native-tSc)<br>MSN4::URA3 (broken with 5'FOA)<br>MSN2::HIS3<br>leu2: ConS-pHsp12-Venus-tPGK1-ConE'-LEU2 (pSc-native-tSc) | Msn2-NLS11 for comparison to Msn2-LANSTrap | Supp Figure 5D |
| yLO237 | BY4741 | ura3: ConS-pRPL18B-Msn2-mScarlet-tADH1-Con1-pTDH3-Hs_RGS2(33-67)_13aa-iRFP713 (iRFP)-asLOV2(404-546)-tPGK1-ConE'-URA3 (pSc-native-tSc)<br>MSN4::URA3 (broken with 5'FOA)<br>MSN2::HIS3<br>leu2: ConS-pHsp12-Venus-tPGK1-ConE'-LEU2 (pSc-native-tSc) | Msn2-mScarlet for comparison to Msn2-LANSTrap | Supp Figure 5D |
| yLO209 | BY4741 | ura3: ConS-pRPL18B-Zdk1-Gal4-mScarlet-yeLANS-tADH1-Con1-pTDH3-Hs_RGS2(33-67)_13aa-iRFP713 (iRFP) -asLOV2(404-546)-tPGK1-ConE-URA3 (pSc-native-tSc)<br>leu2: ConS-pGal1-Venus-tPGK1-ConE-LEU2 (pSc-native-tSc)<br>GAL4::HIS5 (JSO His -- double check) | Gal4-mScarlet | Supp Figure 6A |
| yLO226 | BY4741 | ura3: ConS-pRPL18B-Zdk1-Gal4-mScarlet-tADH1-Con1-pTDH3-Hs_RGS2(33-67)_13aa-iRFP713 (iRFP)-asLOV2 (404-546)-tPGK1-ConE-URA3 (pSc-native-tSc)<br>leu2: ConS-pGal1-Venus-tPGK1-ConE-LEU2 (pSc-native-tSc)<br>GAL4::HIS5 (JSO His -- double check) | Gal4-LANSTrap | Supp Figure 6A-C |
| yAHN437 | BY4741 | leu2: p43_8(8x)-Venus-tPGK1-LEU2 (pSc-native-tSc) | Background strain | Supp Figure 3C |
| ySYC1-78 | yBMH35 | W303a TPK1/2/3-AS Nhp6a-iRFP (Kan) trp1::pAdh1-mCherry-PEF-NESLOVNLS IRFP-nuclear | Original Yumerefendi LANS construct | Supp Figure 3A |
| ySYC1-79 | yBMH35 | W303a TPK1/2/3-AS Nhp6a-iRFP (Kan) trp1::pAdh1-mCherry-PEF-NESLOVNLSvar#3 IRFP-nuclear | class 2 raaKRpRtt | Supp Figure 3A |
| ySYC1-80 | yBMH35 | W303a TPK1/2/3-AS Nhp6a-iRFP (Kan) trp1::pAdh1-mCherry-PEF-NESLOVNLSvar#5 IRFP-nuclear | class 2 paaKRpRtt | Supp Figure 3A |
| ySYC1-81 | yBMH35 | W303a TPK1/2/3-AS Nhp6a-iRFP (Kan) trp1::pAdh1-mCherry-PEF-NESLOVNLSvar#8 IRFP-nuclear | class 2 apaKRaRtt | Supp Figure 3A |
| ySYC2-1 | yBMH35 | W303a TPK1/2/3-AS Nhp6a-iRFP (Kan) trp1::pAdh1-mCherry-PEF-NESLOVNLSvar#11 IRFP-nuclear | class 3 aaaKRswsmaf | Figure1B; Supp Figure 3A |
| ySYC2-2 | yBMH35 | W303a TPK1/2/3-AS Nhp6a-iRFP (Kan) trp1::pAdh1-mCherry-PEF-NESLOVNLSvar#14 IRFP-nuclear | class 3 aaaKRswvmaf | Supp Figure 3A |
| ySYC2-3 | yBMH35 | W303a TPK1/2/3-AS Nhp6a-iRFP (Kan) trp1::pAdh1-mCherry-PEF-NESLOVNLSvar#15 IRFP-nuclear | class 3 aaaKRswsaaf | Supp Figure 3A |
| ySYC2-4 | yBMH35 | W303a TPK1/2/3-AS Nhp6a-iRFP (Kan) trp1::pAdh1-mCherry-PEF-NESLOVNLSvar#20 IRFP-nuclear | bipartite KRpatlandspaaKRR | Supp Figure 3A |
| ySYC2-5 | yBMH35 | W303a TPK1/2/3-AS Nhp6a-iRFP (Kan) trp1::pAdh1-mCherry-PEF-NESLOVNLSvar#4 IRFP-nuclear | class 2 raaKRlRtt | Supp Figure 3A |
| ySYC2-6 | yBMH35 | W303a TPK1/2/3-AS Nhp6a-iRFP (Kan) trp1::pAdh1-mCherry-PEF-NESLOVNLSvar#9 IRFP-nuclear | class 2 paaKRlCtt | Supp Figure 3A |
| ySYC2-7 | yBMH35 | W303a TPK1/2/3-AS Nhp6a-iRFP (Kan) trp1::pAdh1-mCherry-PEF-NESLOVNLSvar#24 IRFP-nuclear | class 1 KRKRwendip | Supp Figure 3A |
| ySYC2-8 | yBMH35 | W303a TPK1/2/3-AS Nhp6a-iRFP (Kan) trp1::pAdh1-mCherry-PEF-NESLOVNLSvar#27 IRFP-nuclear | class 1 psRKRKRdhyav | Supp Figure 3A |
| ySYC2-9 | yBMH35 | W303a TPK1/2/3-AS Nhp6a-iRFP (Kan) trp1::pAdh1-mCherry-PEF-NESLOVNLSvar#29 IRFP-nuclear | class 1 tspsRKRKwdqv | Supp Figure 3A |
| ySYC1-62 | yBMH35 | W303a TPK1/2/3-AS Nhp6a-iRFP (Kan) trp1::pAdh1-Dot6-mCherry (TRP1) IRFP-nuclear | stress-induced localization experiments for Dot6 | Supp Figure 1 |
| ySYC1-70 | yBMH35 | W303a TPK1/2/3-AS Nhp6a-iRFP (Kan) trp1::pAdh1-Crz1-mCherry (TRP1) IRFP-nuclear | stress-induced localization experiments for Crz1 | Supp Figure 1 |
| ySYC1-71 | yBMH35 | W303a TPK1/2/3-AS Nhp6a-iRFP (Kan) trp1::pAdh1-Stb3-mCherry (TRP1) IRFP-nuclear | stress-induced localization experiments for Stb3 | Supp Figure 1 |
| ySYC1-72 | yBMH35 | W303a TPK1/2/3-AS Nhp6a-iRFP (Kan) trp1::pAdh1-Msn2-mCherry (TRP1) IRFP-nuclear | stress-induced localization experiments for Msn2 | Supp Figure 1 |
| ySYC1-74 | yBMH35 | W303a TPK1/2/3-AS Nhp6a-iRFP (Kan) trp1::pAdh1-Msn4-mCherry (TRP1) IRFP-nuclear | stress-induced localization experiments for Msn4 | Supp Figure 1 |
| ySYC1-76 | yBMH35 | W303a TPK1/2/3-AS Nhp6a-iRFP (Kan) trp1::pAdh1-Pho4-mCherry (TRP1) IRFP-nuclear | stress-induced localization experiments for Pho4 | Supp Figure 1 |
| ySYC5-36 | ySYC5-26 | w303a nhp6a::nhp6a-iRFP(KAN) crz1::sfGFP-CandHIS ura::pADH1-zdk-CRZ1(ALA)-mCH-lans;Tdh3-RGS2(URA)<br>leu::pYps1-Venus(LEU) | promoter fusions driven by Crz1*-LANSTrap; pYps1 | Figure 3 |
| ySYC5-37 | ySYC5-26 | w303a nhp6a::nhp6a-iRFP(KAN) crz1::sfGFP-CandHIS ura::pADH1-zdk-CRZ1(ALA)-mCH-lans;Tdh3-RGS2(URA)<br>leu::pGyp7-Venus(LEU) | promoter fusions driven by Crz1*-LANSTrap; pGyp7 | Figure 3 |
| ySYC5-38 | ySYC5-26 | w303a nhp6a::nhp6a-iRFP(KAN) crz1::sfGFP-CandHIS ura::pADH1-zdk-CRZ1(ALA)-mCH-lans;Tdh3-RGS2(URA)<br>leu::pPut1-Venus(LEU) | promoter fusions driven by Crz1*-LANSTrap; pPut1 | Figure 3 |

Supplementary Table 2

| Name | Base Strain | Genome | Description | Figure Correspondence |
|---|---|---|---|---|
| ySYC5-40 | ySYC5-26 | w303a nhp6a::nhp6a-iRFP(KAN) crz1::sfGFP-CandHIS ura::pADH1-zdk-CRZ1(ALA)-mCH-lans;Tdh3-RGS2(URA) leu::pCmk2-Venus(LEU) | promoter fusions driven by Crz1*-LANSTrap; pCmk2 | Figure 3 |
| ySYC5-43 | ySYC5-26 | w303a nhp6a::nhp6a-iRFP(KAN) crz1::sfGFP-CandHIS ura::pADH1-zdk-CRZ1(ALA)-mCH-lans;Tdh3-RGS2(URA) leu::pEna1-Venus(LEU) | promoter fusions driven by Crz1*-LANSTrap; pEna1 | Figure 3 |
| ySYC5-45 | ySYC5-26 | w303a nhp6a::nhp6a-iRFP(KAN) crz1::sfGFP-CandHIS ura::pADH1-zdk-CRZ1(ALA)-mCH-lans;Tdh3-RGS2(URA) leu::pMep1-Venus(LEU) | promoter fusions driven by Crz1*-LANSTrap; pMep1 | Figure 3 |
| ySYC5-34 | ySYC5-26 | w303a nhp6a::nhp6a-iRFP(KAN) crz1::sfGFP-CandHIS ura::pADH1-zdk-CRZ1(ALA)-mCH-lans;Tdh3-RGS2(URA) leu::pPun1-Venus(LEU) | promoter fusion driven by Crz1*-LANSTrap; pPun1 | Supp Figure 11 |
| ySYC5-27 | ySYC5-11 | w303a nhp6a::nhp6a-iRFP(KAN) crz1::sfGFP-CandHIS ura::pADH1-CRZ(WT)-mCH(URA) leu::pPun1-Venus (LEU) | Crz1 WT for CaCl2 experiment; flow; rnaseq | Supp Figure 11 |
| ySYC5-32 | ySYC5-24 | w303a nhp6a::nhp6a-iRFP(KAN) crz1::sfGFP-CandHIS ura::pADH1-zdk-CRZ1(WT)-mCH-lans;Tdh3-RGS2(URA) leu::pPun1-Venus(LEU) | promoter fusion driven by Crz1-LANSTrap; pPun1 | Supp Figure 11 |
| ySYC6-3 | ySYC4-34 | w303a nhp6a::nhp6a-iRFP(KAN) crz1::sfGFP-CandHIS leu::pPun1-venus(LEU) | Pun1 promoter fusion, CrzKO | Supp Figure 11 |
| ySYC5-30 | ySYC5-13 | w303a nhp6a::nhp6a-iRFP(KAN) crz1::sfGFP-CandHIS ura::pADH1-CRZ1(ALA)-mCH(URA) leu::pPun1-Venus (LEU) | untrapped Crz1*; pPun1; flow; rnaseq | Supp Figure 11 |
| ySYC5-71 | ySYC5-69 | w303a nhp6a::nhp6a-iRFP(KAN) crz1::sfGFP-CandHIS ura::pADH1-zdk-CRZ1(5A)-mCH-lans;Tdh3-RGS2(URA) leu::pPun1-Venus(LEU) | crz1(5A); testing equivalence of crz1* and crz1(5A) | Supp Figure 11 |
| ySYC6-23 | ySYC6-7 | w303a nhp6a::nhp6a-iRFP(KAN) crz1::sfGFP-CandHIS ura::pTef1-zdk-CRZ(ALA)-mCH-lans(URA) leu::pGyp7-venus(LEU) | dose response strains (high expression level) | Figure 5 |
| ySYC6-24 | ySYC6-6 | w303a nhp6a::nhp6a-iRFP(KAN) crz1::sfGFP-CandHIS ura::pRpl18b-zdk-CRZ(ALA)-mCH-lans(URA) leu::pGyp7-venus(LEU) | dose response strains (low expression level) | Figure 5 |
| yBMH34 | | w303a Nhp6a-iRFP(Kan) | | background strain |
| yBMH35 | | w303a TPK1/2/3-AS Nhp6a-iRFP (Kan) | | background strain |
| ySYC4-34 | yBMH34 | W303a Nhp6a-iRFP(Kan) crz1::HIS(candida) | | background strain |
| ySYC5-11 | ySYC4-34 | w303a nhp6a::nhp6a-iRFP(KAN) crz1::sfGFP-CandHIS ura::pADH1-CRZ(WT)-mCH(URA) | | background strain |
| ySYC5-13 | ySYC4-34 | w303a nhp6a::nhp6a-iRFP(KAN) crz1::sfGFP-CandHIS ura::pADH1-CRZ1(ALA)-mCH(URA) | | background strain |
| ySYC5-24 | ySYC4-34 | w303a nhp6a::nhp6a-iRFP(KAN) crz1::sfGFP-CandHIS ura::pADH1-zdk-CRZ1(WT)-mCH-lans;Tdh3-RGS2(URA) | | background strain |
| ySYC5-26 | ySYC4-34 | w303a nhp6a::nhp6a-iRFP(KAN) crz1::sfGFP-CandHIS ura::pADH1-zdk-CRZ1(ALA)-mCH-lans;Tdh3-RGS2(URA) | | background strain |
| ySYC5-69 | ySYC4-34 | w303a nhp6a::nhp6a-iRFP(KAN) crz1::sfGFP-CandHIS ura::pADH1-zdk-CRZ1(5A)-mCH-lans;Tdh3-RGS2(URA) | precursor strain for Crz1(5A); pPun1 | background strain |
| ySYC6-6 | ySYC4-34 | w303a nhp6a::nhp6a-iRFP(KAN) crz1::sfGFP-CandHIS ura::pRpl18b-zdk-CRZ(ALA)-mCH-lans(URA) | | background strain |
| ySYC6-7 | ySYC4-34 | w303a nhp6a::nhp6a-iRFP(KAN) crz1::sfGFP-CandHIS ura::pTef1-zdk-CRZ(ALA)-mCH-lans(URA) | | background strain |

Supplementary Table 2

| Name | Background | Marker | Description | Notes | Addgene ID |
|---|---|---|---|---|---|
| pLO405 | | URA3 | URA3 5' Homology-ConS-pRPL18B-Zdk1-mScarlet-yeLANS-tADH1-Con1-pRPL18B-Hs_RGS2(33-67)_13aa-iRFP713 (iRFP)-asLOV2(404-546)-tPGK1-ConE-URA3 (pSc-native-tSc)-URA3 3' Homology-KanR-ColE1 | mScarlet-LANSTrap strain used in Figure 1 and Supplementary Figures; medium expression of mScarlet and pm-LOVTRAP | |
| pLO278 | | URA3 | URA3 5' Homology-ConS-pRPL18B-Zdk1-VP16-ZF43_8-mScarlet-yeLANS-tADH1-Con1-pRPL18B-Hs_RGS2(33-67)_13aa-iRFP713 (iRFP)-asLOV2(404-546)-tPGK1-ConE-URA3 (pSc-native-tSc)-URA3 3' Homology-KanR-ColE1 | SynTF-LANSTrap strain used in Figure 2 and Supplementary Figures; medium expression of SynTF and high expression of pm-LOVTRAP | |
| pLO388 | | LEU2 | LEU2 5' Homology-ConLS'-pPHO84-Venus-tPGK1-ConRE'-LEU2 (pSc-native-tSc)-LEU2 3' Homology-KanR-ColE1 | pPHO84 reporter | |
| pLO361 | | URA3 | URA3 5' Homology-ConS-pRPL18B-Zdk1-Msn2-mScarlet-yeLANS-tADH1-Con1-pTDH3-Hs_RGS2 (33-67)_13aa-iRFP713 (iRFP)-asLOV2(404-546)-tPGK1-ConE'-URA3 (pSc-native-tSc)-URA3 3' Homology-KanR-ColE1 | Msn2-LANSTrap plasmid for strain used in Figure 2 and Supplementary Figures; medium expression of Msn2 and high expression of pm-LOVTRAP | |
| pLO485 | | URA3 | URA3 5' Homology-ConLS'-pTDH3-TOM20_TMD(1-39)-iRFP713 (iRFP)-asLOV2(404-546)-tPGK1-ConRE'-URA3 (pSc-native-tSc)-URA3 3' Homology-KanR-ColE1 | High expression Mito-LOVTRAP | |
| pLO486 | | URA3 | URA3 5' Homology-ConLS'-pRPL18B-TOM20_TMD(1-39)-iRFP713 (iRFP)-asLOV2(404-546)-tPGK1-ConRE'-URA3 (pSc-native-tSc)-URA3 3' Homology-KanR-ColE1 | Medium expression Mito-LOVTRAP | |
| pLO487 | | URA3 | URA3 5' Homology-ConLS'-pREV1-TOM20_TMD(1-39)-iRFP713 (iRFP)-asLOV2(404-546)-tPGK1-ConRE'-URA3 (pSc-native-tSc)-URA3 3' Homology-KanR-ColE1 | Low expression Mito-LOVTRAP | |
| pLO488 | | URA3 | URA3 5' Homology-ConLS'-pTDH3-Hs_RGS2(33-67)_13aa-iRFP713 (iRFP)-asLOV2(404-546)-tPGK1-ConRE'-URA3 (pSc-native-tSc)-URA3 3' Homology-KanR-ColE1 | High expression PM-LOVTRAP | |
| pLO489 | | URA3 | URA3 5' Homology-ConLS'-pRPL18B-Hs_RGS2(33-67)_13aa-iRFP713 (iRFP)-asLOV2(404-546)-tPGK1-ConRE'-URA3 (pSc-native-tSc)-URA3 3' Homology-KanR-ColE1 | Medium expression PM-LOVTRAP | |
| pLO490 | | URA3 | URA3 5' Homology-ConLS'-pREV1-Hs_RGS2(33-67)_13aa-iRFP713 (iRFP)-asLOV2(404-546)-tPGK1-ConRE'-URA3 (pSc-native-tSc)-URA3 3' Homology-KanR-ColE1 | Low expression PM-LOVTRAP | |
| pLO402 | | URA3 | URA3 5' Homology-ConS-pTDH3-Zdk1-mScarlet-yeLANS-tADH1-Con1-pTDH3-Hs_RGS2(33-67)_13aa-iRFP713 (iRFP)-asLOV2(404-546)-tPGK1-ConE-URA3 (pSc-native-tSc)-URA3 3' Homology-KanR-ColE1 | mScarlet-LANSTrap plasmid for strain used in Supplementary Figures; high expression of mScarlet and PM-LOVTRAP | |
| pLO407 | | URA3 | URA3 5' Homology-ConS-pREV1-Zdk1-mScarlet-yeLANS-tADH1-Con1-pREV1-Hs_RGS2(33-67)_13aa-iRFP713 (iRFP)-asLOV2(404-546)-tPGK1-ConE-URA3 (pSc-native-tSc)-URA3 3' Homology-KanR-ColE1 | mScarlet-LANSTrap plasmid for strain used in Supplementary Figures; low expression of mScarlet and PM-LOVTRAP | |
| pLO414 | | URA3 | URA3 5' Homology-ConS-pTDH3-Zdk1-mScarlet-yeLANS-tADH1-Con1-pTDH3-TOM20_TMD(1-39)-iRFP713 (iRFP)-asLOV2(404-546)-tPGK1-ConE-URA3 (pSc-native-tSc)-URA3 3' Homology-KanR-ColE1 | Zdk1-mScarlet-yeLANS + mito-LOVTRAP plasmid for strain used in Supplementary Figures; high expression of mScarlet and mito-LOVTRAP | |
| pLO417 | | URA3 | URA3 5' Homology-ConS-pRPL18B-Zdk1-mScarlet-yeLANS-tADH1-Con1-pRPL18B-TOM20_TMD(1-39)-iRFP713 (iRFP)-asLOV2(404-546)-tPGK1-ConE-URA3 (pSc-native-tSc)-URA3 3' Homology-KanR-ColE1 | Zdk1-mScarlet-yeLANS + mito-LOVTRAP plasmid for strain used in Supplementary Figures; medium expression of mScarlet and mito-LOVTRAP | |
| pLO419 | | URA3 | URA3 5' Homology-ConS-pREV1-Zdk1-mScarlet-yeLANS-tADH1-Con1-pREV1-TOM20_TMD(1-39)-iRFP713 (iRFP)-asLOV2(404-546)-tPGK1-ConE-URA3 (pSc-native-tSc)-URA3 3' Homology-KanR-ColE1 | Zdk1-mScarlet-yeLANS + mito-LOVTRAP plasmid for strain used in Supplementary Figures; low expression of mScarlet and mito-LOVTRAP | |
| pLO316 | | URA3 | URA3 5' Homology-ConS-pRPL18B-VP16-ZF43_8-mScarlet-yeLANS-tADH1-Con1-pRPL18B-Hs_RGS2(33-67)_13aa-iRFP713 (iRFP)-asLOV2(404-546)-tPGK1-ConE-URA3 (pSc-native-tSc)-URA3 3' Homology-KanR-ColE1 | SynTF-yeLANS plasmid for strain used in Supplementary Figures; medium expression of SynTF and PM-LOVTRAP (expressed as control for LANSTrap strains) | |
| pLO366 | | URA3 | URA3 5' Homology-ConS-pRPL18B-Msn2-mScarlet-yeLANS-tADH1-Con1-pTDH3-Hs_RGS2(33-67)_13aa-iRFP713 (iRFP)-asLOV2(404-546)-tPGK1-ConE'-URA3 (pSc-native-tSc)-URA3 3' Homology-KanR-ColE1 | Msn2-yeLANS plasmid for strain used in Supplementary Figures; medium expression of Msn2 and high expression of PM-LOVTRAP (expressed as control for LANSTrap strains) | |
| pLO344 | | URA3 | URA3 5' Homology-ConS-pRPL18B-VP16-ZF43_8-mScarlet-NLS11-tADH1-Con1-pRPL18B-Hs_RGS2(33-67)_13aa-iRFP713 (iRFP)-asLOV2(404-546)-tPGK1-ConE-URA3 (pSc-native-tSc)-URA3 3' Homology-KanR-ColE1 | SynTF-NLS plasmid for strain used in Supplementary Figures; medium expression of SynTF and PM-LOVTRAP (expressed as control for LANSTrap strains) | |
| pLO411 | | URA3 | URA3 5' Homology-ConS-pRPL18B-VP16-ZF43_8-mScarlet-tADH1-Con1-pRPL18B-Hs_RGS2(33-67)_13aa-iRFP713 (iRFP)-asLOV2(404-546)-tPGK1-ConE-URA3 (pSc-native-tSc)-URA3 3' Homology-KanR-ColE1 | SynTF-mScarlet plasmid for strain used in Supplementary Figures; medium expression of SynTF and PM-LOVTRAP (expressed as control for LANSTrap strains) | |
| pLO412 | | URA3 | URA3 5' Homology-ConS-pRPL18B-Pho4-mScarlet-tADH1-Con1-pTDH3-Hs_RGS2(33-67)_13aa-iRFP713 (iRFP)-asLOV2(404-546)-tPGK1-ConE-URA3 (pSc-native-tSc)-URA3 3' Homology-KanR-ColE1 | Pho4-mScarlet plasmid for strain used in Supplementary Figures; medium expression of Pho4 and high expression of PM-LOVTRAP (expressed as control for LANSTrap strains) | |
| pLO465 | | URA3 | URA3 5' Homology-ConS-pRPL18B-Pho4-mScarlet-NLS11-tADH1-Con1-pTDH3-Hs_RGS2(33-67)_13aa-iRFP713 (iRFP)-asLOV2(404-546)-tPGK1-ConE-URA3 (pSc-native-tSc)-URA3 3' Homology-KanR-ColE1 | Pho4-NLS plasmid for strain used in Supplementary Figures; medium expression of Pho4 and high expression of PM-LOVTRAP (expressed as control for LANSTrap strains) | |
| pLO463 | | URA3 | URA3 5' Homology-ConS-pRPL18B-Msn2-mScarlet-NLS11-tADH1-Con1-pTDH3-Hs_RGS2(33-67)_13aa-iRFP713 (iRFP)-asLOV2(404-546)-tPGK1-ConE'-URA3 (pSc-native-tSc)-URA3 3' Homology-KanR-ColE1 | Msn2-NLS plasmid for strain used in Supplementary Figures; medium expression of Msn2 and high expression of PM-LOVTRAP (expressed as control for LANSTrap strains) | |
| pLO480 | | URA3 | URA3 5' Homology-ConS-pRPL18B-Msn2-mScarlet-tADH1-Con1-pTDH3-Hs_RGS2(33-67)_13aa-iRFP713 (iRFP)-asLOV2(404-546)-tPGK1-ConE'-URA3 (pSc-native-tSc)-URA3 3' Homology-KanR-ColE1 | Msn2-mScarlet plasmid for strain used in Supplementary Figures; medium expression of Msn2 and high expression of PM-LOVTRAP (expressed as control for LANSTrap strains) | |

Supplementary Table 3

| Name | Background | Marker | Description | Notes | Addgene ID |
|---|---|---|---|---|---|
| pLO364 | | URA3 | URA3 5' Homology-ConS-pRPL18B-Zdk1-Gal4-mScarlet-yeLANS-tADH1-Con1-pTDH3-Hs_RGS2 (33-67)_13aa-iRFP713 (iRFP)-asLOV2(404-546)-tPGK1-ConE-URA3 (pSc-native-tSc)-URA3 3' Homology-KanR-ColE1 | Gal4-LANSTrap plasmid for strain used in Supplementary Figures; medium expression of Gal4 and high expression of PM-LOVTRAP | |
| pLO413 | | URA3 | URA3 5' Homology-ConS-pRPL18B-Zdk1-Gal4-mScarlet-tADH1-Con1-pTDH3-Hs_RGS2(33-67) _13aa-iRFP713 (iRFP)-asLOV2(404-546)-tPGK1-ConE-URA3 (pSc-native-tSc)-URA3 3' Homology-KanR-ColE1 | Gal4-mScarlet plasmid for strain used in Supplementary Figures; medium expression of Gal4 and high expression of PM-LOVTRAP (expressed as control for LANSTrap strain) | |
| pAN736 | | LEU2 | LEU2 5' Homology-p43_8(8x)-Venus-tPGK1-LEU2 (pSc-native-tSc)-LEU2 3' Homology-KanR-ColE1 | pSynTF reporter | |
| pLO363 | | URA3 | URA3 5' Homology-ConS-pRPL18B-Zdk1-Pho4-mScarlet-yeLANS-tADH1-Con1-pTDH3-Hs_RGS2 (33-67)_13aa-iRFP713 (iRFP)-asLOV2(404-546)-tPGK1-ConE-URA3 (pSc-native-tSc)-URA3 3' Homology-KanR-ColE1 | Pho4-LANSTrap plasmid for strain used in Figure 2 and Supplementary Figures; medium expression of Pho4 and high expression of PM-LOVTRAP | |
| pLO352 | | LEU2 | LEU2 5' Homology-ConS-pHsp12-Venus-tPGK1-ConE'-LEU2 (pSc-native-tSc)-LEU2 3' Homology-KanR-ColE1 | pHSP12 reporter | |
| YTK120 | | HIS3 | C.G. HIS3 | HIS3 cassette | |
| pAN160 | | LEU2 | LEU2 5' Homology-ConS-pGal1-Venus-tPGK1-ConE-LEU2 (pSc-native-tSc)-LEU2 3' Homology-KanR-ColE1 | pGAL1 reporter | |
| pSYC084 | pSYC081 | TRP1 | pAdh1-mCherry-PEF-NESLOVNLS-JSO179 | original LANS Yumerefendi 2015 | |
| pSYC085 | pSYC084 | TRP1 | pAdh1-mCherry-PEF-NESLOVNLSvar#3-JSO179 | NLS variant | |
| pSYC086 | pSYC084 | TRP1 | pAdh1-mCherry-PEF-NESLOVNLSvar#4-JSO179 | NLS variant | |
| pSYC087 | pSYC084 | TRP1 | pAdh1-mCherry-PEF-NESLOVNLSvar#5-JSO179 | NLS variant | |
| pSYC088 | pSYC084 | TRP1 | pAdh1-mCherry-PEF-NESLOVNLSvar#8-JSO179 | NLS variant | |
| pSYC089 | pSYC084 | TRP1 | pAdh1-mCherry-PEF-NESLOVNLSvar#9-JSO179 | NLS variant | |
| pSYC090 | pSYC084 | TRP1 | pAdh1-mCherry-PEF-NESLOVNLSvar#11-JSO179 | NLS variant | |
| pSYC091 | pSYC084 | TRP1 | pAdh1-mCherry-PEF-NESLOVNLSvar#14-JSO179 | NLS variant | |
| pSYC092 | pSYC084 | TRP1 | pAdh1-mCherry-PEF-NESLOVNLSvar#15-JSO179 | NLS variant | |
| pSYC093 | pSYC084 | TRP1 | pAdh1-mCherry-PEF-NESLOVNLSvar#20-JSO179 | NLS variant | |
| pSYC094 | pSYC084 | TRP1 | pAdh1-mCherry-PEF-NESLOVNLSvar#24-JSO179 | NLS variant | |
| pSYC095 | pSYC084 | TRP1 | pAdh1-mCherry-PEF-NESLOVNLSvar#27-JSO179 | NLS variant | |
| pSYC096 | pSYC084 | TRP1 | pAdh1-mCherry-PEF-NESLOVNLSvar#29-JSO179 | NLS variant | |
| pSYC081 | pJSO179 | TRP1 | PEF_NESLOVNLS_SacI-JSO179 | intermediate plasmid | |
| pJSO179 | pNH604 | TRP1 | p604; basic single integration vector from Lim lab | base plasmid | |
| pJSO529 | pNH604 | TRP1 | 604-Adh1-Dot6-mCherry | tagged TF to test for localization dynamics in response to environmental inputs | |
| pJSO568 | pNH604 | TRP1 | 604-pAdh1-Crz1-mcherry | tagged TF to test for localization dynamics in response to environmental inputs | |
| pJSO269 | pNH604 | TRP1 | 604-pAdh1-Stb3-mCherry | tagged TF to test for localization dynamics in response to environmental inputs | |
| pJSO392 | pNH604 | TRP1 | 604-pAdh1-MSN2-mCherry | tagged TF to test for localization dynamics in response to environmental inputs | |
| pJSO285 | pNH604 | TRP1 | 604-pAdh1-Msn4-mCherry | tagged TF to test for localization dynamics in response to environmental inputs | |
| pJSO569 | pNH604 | TRP1 | 604-pAdh1-Pho4-mcherry | tagged TF to test for localization dynamics in response to environmental inputs | |
| pSYC332 | pYTK143 | LEU2 | pYPS1-Venus | promoter fusion | |
| pSYC392 | pYTK143 | LEU2 | pGYP7-Venus | promoter fusion | |
| pSYC393 | pYTK143 | LEU2 | pPUT1-Venus | promoter fusion | |
| pSYC395 | pYTK143 | LEU2 | pCMK2-Venus | promoter fusion | |
| pSYC398 | pYTK143 | LEU2 | pENA1-Venus | promoter fusion | |
| pSYC400 | pYTK143 | LEU2 | pMEP1-Venus | promoter fusion | |
| pSYC331 | pYTK143 | LEU2 | pPUN1-Venus | promoter fusion | |
| pSYC295 | pYTK146 | URA3 | pADH1-crz1-mcherry-tPGK1 | crz1 WT characterization | |
| pSYC293 | pYTK146 | URA3 | pADH1-crz1(19A)-mcherry-tPGK1 | crz1 19A mutant characterization | |
| pSYC310 | pYTK146 | URA3 | pADH1-zdk-crz1-mcherry-lans-tPGK1; pTDH3-Rgs2 | lanstrap crz1 WT characterization | |
| pSYC308 | pYTK146 | URA3 | pADH1-zdk-crz1(19A)-mcherry-lans-tPGK1; pTDH3-Rgs2 | lanstrap crz1* WT characterization | |
| pSYC346 | pYTK146 | URA3 | pADH1-zdk-crz1(5A)-mcherry-lans; pTDH3-Rgs2 | lanstrap crz1 5A characterization | |
| pSYC410 | pYTK146 | URA3 | pTEF1-zdk-crz1(19A)-mcherry-lans; pTDH3-Rgs2 | crz1 dose response | |
| pSYC411 | pYTK146 | URA3 | pRPL18b-zdk-crz1(19A)-mcherry-lans; pTDH3-Rgs2 | crz1 dose response | |
| pYTK143 | | LEU2 | LS' LEU (YTK) KanR RE' | Pre-Assembled Integration Vector | |
| pYTK146 | | URA3 | LS' URA (HES) KanR RE' | Pre-Assembled Integration Vector | |
| pLO263 | | AmpR | pTDH3-Rgs2-iRFP713-asLOV2(404-546)-tPGK1 | plasma membrane LOV2 tag for multi-part assembly of final LANSTrap construct | |

Supplementary Table 3

| Name | Background | Marker | Description | Notes | Addgene ID |
|---|---|---|---|---|---|
| pAN451 | | AmpR | ConS-GFP dropout (+BsaI)-Con1-AmpR-ColE1 | intermediate cloning vector | |

Supplementary Table 3

| Light Dose | milliWattage | Std Dev | Date |
|---|---|---|---|
| 0 | -0.0001499 | 0.000200[illegible] | 20181205 |
| 0 | -0.0002513 | 0.000252 | 20181128 |
| 0 | -0.001819 | 0.000978 | 20181109 |
| 64 | 0.1011 | 0.008522 | 20181205 |
| 64 | 0.07921 | 0.007053 | 20181128 |
| 64 | 0.1279 | 0.01123 | 20181109 |
| 128 | 0.1944 | 0.02126 | 20181205 |
| 128 | 0.1962 | 0.01824 | 20181128 |
| 128 | 0.2633 | 0.02653 | 20181109 |
| 256 | 0.3961 | 0.0393 | 20181205 |
| 256 | 0.4012 | 0.0362 | 20181128 |
| 256 | 0.5107 | 0.04832 | 20181109 |
| 512 | 0.7849 | 0.07396 | 20181205 |
| 512 | 0.7776 | 0.09084 | 20181128 |
| 512 | 1.041 | 0.09508 | 20181109 |
| 1024 | 1.53 | 0.1346 | 20181205 |
| 1024 | 1.477 | 0.1491 | 20181128 |
| 1024 | 1.986 | 0.2349 | 20181109 |
| 2048 | 3.088 | 0.3326 | 20181205 |
| 2048 | 3.026 | 0.2563 | 20181128 |
| 2048 | 3.45 | 0.9 | 20181109 |
| 3072 | 4.302 | 0.4414 | 20181205 |
| 3072 | 4.235 | 0.4559 | 20181128 |
| 4095 | 5.407 | 0.5494 | 20181205 |
| 4095 | 5.486 | 0.4617 | 20181128 |
| 4095 | 6.56 | 0.587 | 20181109 |

Supplementary Table 4
